# Control of leaf shape through non-cell autonomous TCP4 regulation

**DOI:** 10.64898/2026.05.09.723987

**Authors:** Vishwadeep Mane, Amrita Saxena, Constance Le Gloanec, Abhishek Gupta, Ganesh Venkatesh, Simone Bovio, Daniel Kierzkowski, Olivier Hamant, Utpal Nath

## Abstract

Leaf development requires coordinated growth across distinct tissue layers, integrating genetic regulation with mechanical coupling to achieve proper organ shaping. However, the mechanisms coordinating differentiation and growth across layers to ensure coherent leaf morphogenesis remain poorly understood. Therefore, we investigated the role of TCP4 as a potential central regulator of interlayer communication coupling genetic and mechanical signals during *Arabidopsis* leaf development. Three-dimensional analyses revealed pronounced asymmetry in the expression of TCP4 and its negative regulator, MIR319C, across developing primordia, establishing spatially patterned differentiation cues. Tissue-specific perturbations demonstrated that TCP4 activity in either epidermis or subepidermis modulates cell proliferation and expansion across layers through both protein mobility and mobility-independent signalling mechanisms. Strong epidermal TCP4 induction arrested leaf development by prematurely suppressing proliferation and preventing symmetry breaking, thereby locking primordia in cylindrical states. Live imaging revealed fundamental reprogramming of growth anisotropy, with TCP4 driving cortical microtubule alignment, increasing cell wall stiffness, and establishing layer-specific pectin patterns. Together, these findings establish TCP4 as an integrator of genetic and mechanical signals across leaf tissue layers, coupling transcriptional programmes to cytoskeletal organisation, cell wall remodelling, and cross-layer mechanical feedback.

## Introduction

True leaves emerge as asymmetric finger-like primordia on the flanks of the dome-shaped shoot apical meristem (SAM) through coordinated genetic and mechanical signalling networks (Barkoulas et al., 2008; Kuhlemeier & Timmermans, 2016; De Carufel et al., 2026). Following emergence, these multilayered primordia, derived from clonally distinct SAM layers (Stewart & Dermen, 1970; Reinhardt et al., 2003), undergo progressive flattening to form the leaf blade. This morphogenetic programming coordinates growth across tissue compartments to establish final organ shape, size, and planarity (Efroni et al., 2008; De Carufel et al., 2026). Central to this programme is the spatiotemporal coordination of cell proliferation and differentiation, which generates a basipetal growth gradient essential for robust leaf morphology (Donnelly et al., 1999; Andriankaja et al., 2012; Le Gloanec et al., 2024).

Genetic circuits governing the proliferation-differentiation transition are central to establishing these growth patterns (Hamant & Traas, 2010; Sapala et al., 2018). In *Arabidopsis thaliana*, the JAW-TCP transcription factor family (TCP2, TCP3, TCP4, TCP10, and TCP24), which is negatively regulated by the MIR319 microRNA, plays a key role in this process (Palatnik et al., 2003; Schommer et al., 2008). Loss of JAW-TCP function through MIR319A overexpression (*jaw-D* mutant) results in enlarged leaves with ruffled margins and negative Gaussian curvature (Palatnik et al., 2003; Nath et al., 2003). At the cellular level, this phenotype reflects increased cell number and reduced cell size through cell-organ compensation (Tsuge et al., 1996; Horiguchi et al., 2005), indicating that JAW-TCPs typically restrict leaf growth by promoting differentiation (Ori et al., 2007). Although TCP function has been extensively characterised in the epidermis (Efroni et al., 2010; Koyama et al., 2010; Challa et al., 2019), how TCP-mediated differentiation signals coordinate growth across tissue layers remains unclear.

Along with genetic circuits, mechanical feedback influences tissue stress and growth behaviour, mediating coordination between tissue layers during development (Green, 1962; Zhao et al., 2020). Plant cells behave as pressurised vessels in which turgor-induced stresses orient cortical microtubules (CMTs) along the maximal stress directions (Cosgrove, 2005; Hamant et al., 2008; Zhao et al., 2020). These oriented CMTs guide cellulose microfibril (CMF) deposition, reinforcing growth anisotropy. Through coordinated CMT-CMF dynamics, mechanical signals propagate across shared cell walls, enabling tissues to sense and respond to local growth heterogeneity. Cell wall yielding and remodelling occur through crosstalk with genetic programmes (Paredez et al., 2006; Elsner et al., 2012), thereby integrating molecular and mechanical information across tissue compartments. Because plant cells are mechanically coupled through shared walls, stress-mediated signalling is essential for coordinating proliferation and expansion across the organ. (Peaucelle et al., 2011; Uyttewaal et al., 2012). Mechanical feedback also contributes to resolving global growth conflicts associated with proliferation-differentiation transitions and developmental-stage-dependent changes in cell wall properties (Kwiatkowska, 2004; Hervieux et al., 2016; Coen and Cosgrove, 2023; Lapointe et al., 2025).

During leaf development, growth and mechanical coupling between layers are critical for transforming the cylindrical primordium into a flat lamina (Boudaoud, 2010; Sampathkumar et al., 2014; Hervieux et al., 2017; Qi et al., 2017; Zhao et al., 2020). Growth of inner tissues places the epidermis under tension, while the epidermis constrains internal layers through tissue-wide CMT orientations and anisotropic CMF deposition (Boudaoud, 2010; Qi et al., 2017; Zhao et al., 2020; Silveira et al., 2025). Simultaneously, anisotropic wall reinforcement in subepidermal layers along the adaxial-abaxial axis promotes lamina expansion (Zhao et al., 2020; Guo et al., 2022). These reciprocal mechanical interactions enable continuous cross-layer communication that coordinates symmetry breaking and planar expansion (Zhao et al., 2020). Consequently, differentiation timing and wall stiffening must be precisely synchronised across layers to achieve coordinated planar growth (Zhao et al., 2020). Despite the importance of differentiation timing for mechanical coupling, how developmental regulators like TCP4 influence this process across tissue layers remains poorly understood. Given the role of TCP4 in promoting the proliferation-differentiation transition (Rodriguez et al., 2010; Challa et al., 2019) and its multilayered expression pattern, an important unresolved question is how TCP4 activity influences cross-layer communication and mechanical coupling during morphogenesis.

To address this question, we restricted TCP4 activity to specific tissue layers using the PDF1 and AN3 promoters. PDF1 drives highly specific expression in the epidermal L1 layer (Abe et al., 2003), whereas AN3 is expressed in the subepidermal L2 layer (Kawade et al., 2013). By restricting TCP4 to a single tissue layer, this approach enables dissection of cell-autonomous differentiation effects from non-cell-autonomous mechanical consequences, directly testing how premature differentiation in one tissue layer impacts differentiation and mechanical coupling within adjacent tissue layers.

Here, we investigate how TCP4-mediated differentiation influences cross-layer mechanical coupling and whether layered expression pattern of TCP4 modulates differentiation kinetics and intensity. Using layer-specific genetic perturbations, quantitative morphometric analyses, and mechanical measurements, we show that the spatial distribution of TCP4 across tissue compartments is essential for maintaining effective mechanical coupling during morphogenesis. Our findings establish that three-dimensional gene expression architecture governs cross-layer mechanical coupling, integrating molecular and physical signals to ensure robust organ morphogenesis.

## Results

### 1. The TCP4-MIR319C module exhibits asymmetrically layered expression during early leaf development

Previous studies have shown that expression of a microRNA-resistant version of TCP4 (rTCP4) under its endogenous promoter in the *jaw-D* mutant background restores organ shape and size beyond wild-type levels, accompanied by reduced cell number and compensatory increases in cell size (Nag et al., 2009; Das Gupta & Nath, 2015). At the organ scale, TCP4 protein and the MIR319C promoter display mutually exclusive expression domains. TCP4 accumulates in distal regions, whereas MIR319C is predominantly expressed proximally when leaf primordia reach approximately 120-150 μm in length (Palatnik et al., 2007; Shankar et al., 2023). TCP4 protein subsequently spreads basipetally across the lamina, coinciding with the progression of the growth arrest and cellular expansion (Danisman et al., 2012; Shankar et al., 2023). However, these dynamics have been primarily characterised in two dimensions and do not address whether this spatial organisation extends across tissue layers. Given the established roles of JAW-TCPs in regulating leaf size, shape, and curvature (Nath et al., 2003; Ori et al., 2007), and the importance of mechanical coupling between tissue layers for morphogenesis, we asked whether the TCP4-MIR319C regulatory module exhibits three-dimensional expression asymmetry across the developing leaf.

Whole-mount GUS staining of cleared leaf primordia (8^th^ leaf) showed that TCP4 promoter activity and TCP4 protein accumulation occupy distal domains, whereas MIR319C promoter activity is confined proximally (Fig. 1A), consistent with previous reports (Shankar et al., 2023). To determine how these spatial patterns extend across the different tissue layers of the leaf, we examined TCP4 and MIR319C activity in three dimensions. We used previously reported GUS reporter lines to monitor TCP4 promoter activity (*pTCP4::GUS*), TCP4 protein accumulation (*pTCP4::TCP4-GUS*) and MIR319C promoter activity (*pMIR319C::GUS*), all in the Col-0 background (Palatnik et al., 2007; Shankar et al., 2023). GUS staining was performed on the first pair of true leaves at 2-3 days after initiation (DAI), followed by modified pseudo-Schiff’s propidium iodide (mPS-PI) counterstaining to reveal tissue architecture (Truernit et al., 2008). Confocal reflection microscopy and three-dimensional image reconstruction in MorphoGraphX (Barbier de Reuille et al., 2015; Strauss et al., 2022) were used to generate optical sagittal and transverse sections spanning distal, middle, and proximal regions of the lamina.

**Fig. 1.**
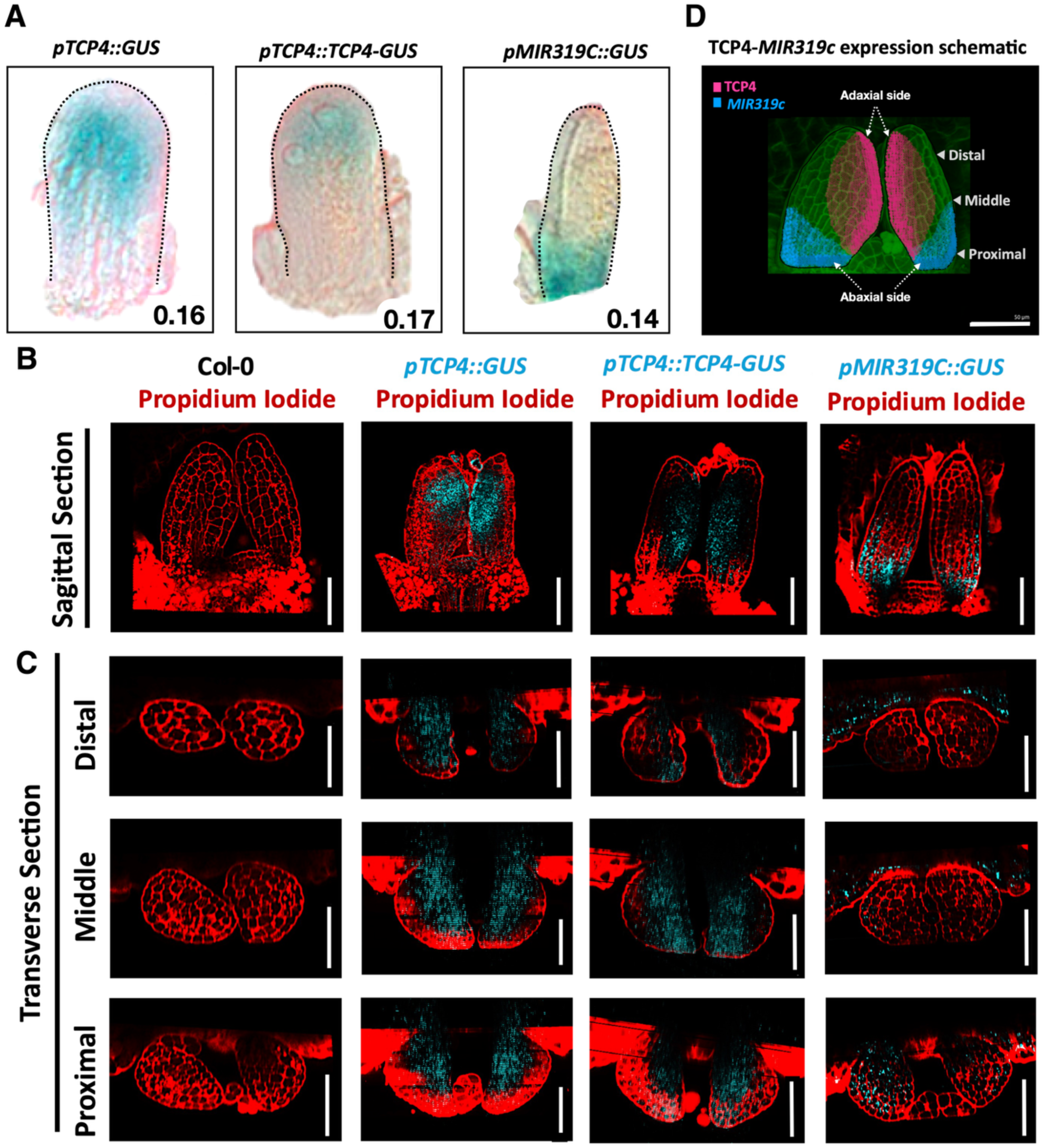
Expression pattern of the TCP4-miR319c module across three dimensions reveals an asymmetric gradient during early leaf development. (A) Bright-field images of whole-mount GUS reporter assays of the 8th incipient leaf primordium showing promoter activity of *TCP4* and *MIR319c*, together with TCP4 protein activity driven by the endogenous promoter. TCP4 promoter and protein activities are enriched at the distal end of the primordium, whereas miR319c promoter activity is confined to the proximal end, exhibiting mutually exclusive spatial domains in young leaves. Numbers below the leaf primordium images indicate the leaf length in mm, and outlines of the primordia are hand-drawn. (B) Optical mid-sagittal section (taken through the midrib) and (C) optical transverse section of modified pseudo-schiff’s-propidium iodide (mPS-PI) and GUS-stained leaves indicating promoter activities of TCP4 and miR319c, along with TCP4 protein localisation. Col-0 serves as a negative control. The pTCP4 domain extends from the distal toward the proximal end with an asymmetric distribution from the adaxial to the abaxial side across tissue layers. The spatial distribution of TCP4 protein closely mirrors its promoter activity. In contrast, the pMIR319c domain remains restricted to the proximal region, with an expression gradient extending from the abaxial toward the adaxial side. (Scale bar = 50 µm for all confocal images). (D) Schematic representation of a 3D reconstruction of the TCP4 (pink) and miR319c (blue) module, superimposed with data obtained from panels (B) and (C) onto a confocal image of a leaf primordium. TCP4 protein displays a gradient from the adaxial to the abaxial side, whereas miR319c expression is restricted to the proximal region and shows a subtle gradient from the abaxial to the adaxial side.

Analysis of mid-sagittal and transverse sections revealed that both TCP4 promoter activity and TCP4 protein accumulation exhibit a pronounced gradient from the adaxial to the abaxial side of the leaf (Fig. 1B, C). In primordia measuring approximately 150-200 μm, TCP4 protein levels were higher in the distal and middle regions than in the proximal region. In contrast, promoter activity remained relatively uniform along the longitudinal axis (Fig. 1B). In contrast, *pMIR319C::GUS* activity was predominantly confined to proximal regions. It displayed an opposing gradient from the abaxial toward the adaxial side (Fig. 1C). This pattern was mutually exclusive with TCP4 protein accumulation in mid-sagittal and proximal sections, consistent with their antagonistic regulatory relationship. This divergence between promoter activity and protein distribution is consistent with post-transcriptional restriction by MIR319C. No GUS signal was detected in control plants lacking reporter constructs (Fig. 1B, C).

Together, these observations demonstrate that the TCP4-MIR319C regulatory module is organised along both longitudinal and adaxial-abaxial axes and exhibits pronounced asymmetry across tissue layers during early leaf development (Fig. 1D). In particular, TCP4 expression is enhanced on the adaxial and distal side of the developing leaf.

### 2. Inducible and tissue-specific TCP4 expression suggests a layer-independent function for TCP4 at first order, and a primary role for the epidermis, at second order

TCP4 is expressed across epidermal and subepidermal tissues during early leaf development, raising the question of whether its activity in distinct tissue layers contributes equally to organ growth. The epidermis can exert a major influence on organ growth by integrating biochemical and mechanical signals and by constraining the expansion of underlying tissues (Kutschera & Niklas, 2007; Silvera et al., 2025). Tissue-specific perturbations of brassinosteroid, ethylene, and auxin-related pathways have further demonstrated that signalling initiated in the epidermis can channel growth at the organ scale (Savaldi-Goldstein et al., 2007; Vaseva et al., 2018; Kierzkowski et al., 2013). The graded, multilayered distribution of TCP4 therefore prompted us to investigate how its spatial activity contributes to growth coordination across tissue layers. To address this, we turned to an inducible, tissue-specific expression system that enables precise manipulation of TCP4 levels across tissue layers. We employed previously reported transgenic lines and generated multiple independent transgenic lines expressing dexamethasone-inducible rTCP4 under promoters with distinct spatial activities. To reproduce the endogenous TCP4 expression pattern, we used *pTCP4::rTCP4-GR* in Col-0 and *jaw-D* backgrounds (hereafter Col-0;GR and *jaw-D;GR,* respectively; Challa et al., 2019). For tissue-specific expression, we expressed rTCP4-GR under the epidermal-specific PDF1 promoter (*pPDF1::rTCP4-GR* in *jaw-D;* hereafter *PDF1;GR*; Abe et al, 2003; Fig. S1A, C) or the subepidermal AN3 promoter (*pAN3::rTCP4-GR* in *jaw-D;* hereafter, *AN3;GR*; Kawade et al; 2013; Fig. S1B, C; also see Table S1). This approach enables direct comparison of TCP4 activity in its native spatial domain and when restricted to the epidermal or subepidermal layers, while avoiding the developmental lethality associated with strong constitutive epidermal expression.

Under all conditions, upon dexamethasone (DEX) induction, we observed a reduction in leaf size (Fig. 2A, B), consistent with the known role of TCP4 in promoting cell differentiation. The magnitude of the response varied among tissue-specific lines and independent transgenic insertions. In Col-0;GR, DEX induction of rTCP4 under its endogenous promoter resulted in a significant reduction in the first leaf area (MOCK: 22.20 ± 3.40 mm²; DEX: 15.70 ± 3.70 mm²; ∼32% reduction; Fig. 2A, B; S1G). In *jaw-D;GR*, where endogenous TCP4 levels are reduced due to miR319A overexpression, DEX induction of rTCP4 rescued leaf area and shape, even exceeding wild-type levels (MOCK: 36.0 ± 7.0 mm²; DEX: 14.2 ± 2.7 mm²; ∼60% reduction; Fig. 2A, B, S1G).

**Fig. 2.**
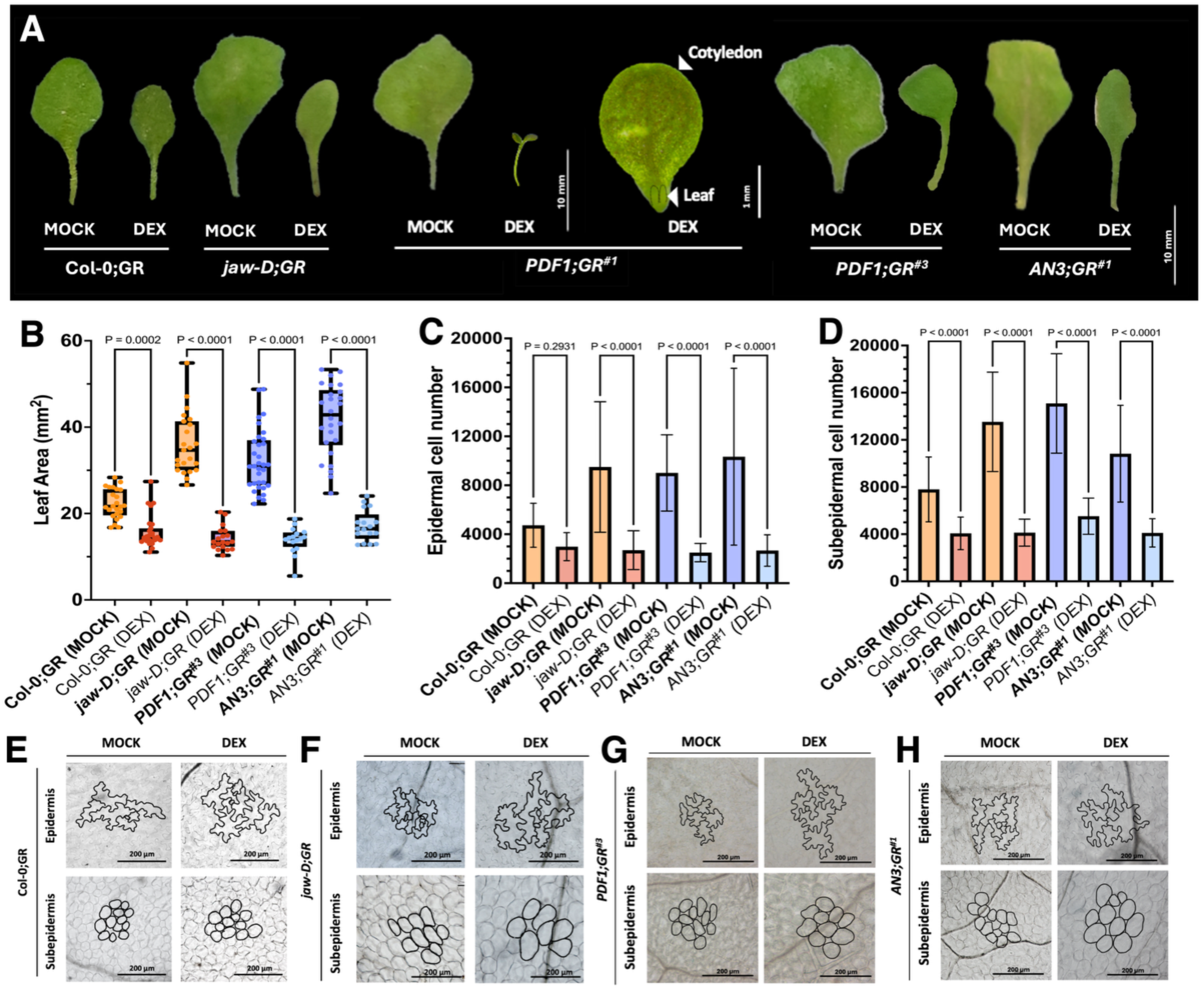
TCP4-dependent signalling integrates cell proliferation and expansion across tissue layers independently of spatial expression. (A) Representative images of the mature first leaf pair from 28-day-old plants of the indicated genotypes and chemical treatments. Scale bar, 10 mm. (B) Quantification of leaf area (Y-axis) for the mature first leaf pair of the indicated genotypes and treatments (X-axis). N = 14-34 leaves. Statistical significance was assessed by one-way ANOVA followed by a Šídák post hoc test (P-values indicated above the compared means). Error bars represent SD. (C, D) Estimated epidermal (G) and subepidermal (H) cell numbers for the indicated genotypes and treatments (X-axis). Cell number was calculated as the ratio of total leaf area to mean cell area (16-20 leaves per genotype). Statistical analysis as in (B); error bars represent SD. (E-H) Differential interference contrast (DIC) images of epidermal pavement cells and subepidermal cells from mature first leaves of 28-day-old plants under the indicated genotypes and chemical treatments. Cell outlines are shown in black. Scale bars, 200 µm.

Among the transgenic lines analysed, the most severe phenotypes were observed following epidermal TCP4 induction. *PDF1;GR*^#1^ and *PDF1;GR*^#2^ failed to form any true leaves when rTCP4 was induced by stratifying seeds on dexamethasone-containing medium (Fig. 2A; S1D, G). While seeds germinated and cotyledons developed, plants displayed elongated hypocotyls reminiscent of TCP4-induced cell elongation, and two arrested leaf primordia-like structures, with no subsequent visible leaf outgrowth (Fig. 2A; S1G). This complete developmental arrest underscores the severe consequences of high levels of epidermal TCP4 activity and the necessity to use weaker transgenic insertions to investigate its effects without abolishing leaf development. We therefore analysed weaker insertions to determine how phenotypic severity varied with transgene strength. *PDF1; GR*^#3^ showed robust rescue of the *jaw-D* leaf phenotype upon dexamethasone treatment (Fig. 2A; S1G), with leaf area reduced by ∼53% (MOCK: 32.2 ± 7.7 mm²; DEX: 15.1 ± 2.4 mm²), approaching the magnitude of rescue observed in *jaw-D;GR* (∼60%; Fig. 2B).

Last, in the stronger *AN3;GR*^#1^ insertion, dexamethasone-induced rTCP4 activity in the subepidermis fully rescued the *jaw-D* leaf phenotype (Fig. 2A; S3C), reducing leaf area by ∼58% (MOCK: 40.75 ± 8.21 mm²; DEX: 17.15 ± 3.58 mm²; Fig. 2B). This effect was comparable to that observed following epidermal TCP4 induction (*PDF1;GR*^#3^: ∼53%). However, because rTCP4 abundance was not quantitatively matched between the *PDF1;GR* and *AN3;GR* lines, differences between individual lines cannot be attributed solely to the tissue in which TCP4 was induced. We therefore examined multiple independent insertions of each construct to account for variation in transgene strength (Fig. S1 and S3).

Altogether, the ability of both epidermal and subepidermal TCP4 induction indicates that restricting TCP4 activity to either tissue compartment is sufficient to substantially alter leaf growth. Although the effects were generally more pronounced following epidermal induction, differences in transgene abundance between lines preclude a direct quantitative comparison of TCP4 activity between tissue layers. Moreover, the similar phenotypic responses produced by layer-specific induction could reflect non-cell-autonomous effects, including intercellular movement of TCP4 or secondary effects on mechanical coupling between tissue layers.

### 3. TCP4 hinders cell proliferation and promotes cell elongation in all cell layers, with a more pronounced effect in the epidermis

To determine how layer-specific TCP4 activity affects growth at the cellular level, we quantified cell number and size in the epidermal and subepidermal layers across the different transgenic lines. Across genotypes, DEX induction generally reduced cell number and increased final cell size in both tissue layers, although the magnitude of these responses differed between the epidermis and subepidermis (Fig. 2C-H).

In Col-0;GR, DEX induction resulted in a non-significant reduction in epidermal cell number (MOCK: 4734 ± 1797; DEX: 2986 ± 1146; ∼37% reduction; Fig. 2C) and a significant reduction in subepidermal cell number (MOCK: 7798 ± 2749; DEX: 4070 ± 1381; ∼48% reduction; Fig. 2D). Subepidermal cell numbers remained consistently higher than epidermal cell numbers under both conditions. Reduced cell number was accompanied by an increase in mean cell size in both layers, which was more pronounced in the subepidermis (∼36%; MOCK: 2850 ± 795 µm²; DEX: 3864 ± 907 µm²; Fig. 2E, S1F) than in the epidermis (∼12%; MOCK: 4692 ± 1645 µm²; DEX: 5267 ± 1799 µm²; Fig. 2E, S1E).

In *jaw-D;GR* background, DEX induction markedly reduced cell number in both the epidermis (∼71%; MOCK: 9494 ± 5337; DEX: 2698 ± 1584; Fig. 2C) and subepidermis (∼68%; MOCK: 13525 ± 4213; DEX: 4131 ± 1145; Fig. 2D), accompanied by increases in final cell size of ∼39% and ∼29% respectively (Fig. 2F; S1E, F). Compared with Col-0;GR under MOCK conditions, *jaw-D;GR* exhibited higher cell numbers in both layers, which were strongly reduced following rTCP4 induction (Tables S4 and S5).

In the MOCK condition, both *PDF1;GR*^#1^ and *PDF1;GR*^#2^ preserved the *jaw-D* background phenotype (Fig. 2A; S1D, G) with leaf areas indistinguishable from the *jaw-D;GR* control (*jaw-D;GR*: 37.4 ± 6.5 mm²; *PDF1;GR*^#1^: 34.7 ± 5.6 mm²; *PDF1;GR*^#2^: 43.1 ± 6.5 mm²; Fig. S1H).

Similarly, epidermal and subepidermal cell numbers (Fig. S1I, J) and cell sizes in MOCK-treated *PDF1;GR*^#1^ and *PDF1;GR*^#2^ were comparable to those in *jaw-D;GR* (Fig. S2A, C and D), indicating that the *jaw-D* phenotype was maintained in the absence of induction and providing no evidence of transgene leakiness (Tables S4 and S5).

In *PDF1;GR^#3^*, cell numbers decreased significantly in both the epidermis (∼71%; MOCK: 9010 ± 3117; DEX: 2498 ± 749; Fig. 2C) and subepidermis (∼63%; MOCK: 15088 ± 4218; DEX: 5525 ± 1538; Fig. 2D), demonstrating that epidermal rTCP4 induction affects cell proliferation across both tissue layers (Tables S4 and S5). In contrast, the increase in final cell size was strongly layer dependent: epidermal cell size increased by ∼58% (MOCK: 3578 ± 958 µm²; DEX: 5667 ± 1142 µm²; Fig. 2G; S1E), whereas subepidermal cell size increased by only ∼20% (MOCK: 2136 ± 489 µm²; DEX: 2562 ± 451 µm²; Fig. 2G; S1F) (Tables S4 and S5).

Together, these results show that epidermal rTCP4 induction affects cell number in both tissue layers, while its effect on final cell size is substantially stronger in the epidermis. However, because rTCP4 abundance was not quantitatively matched between the tissue-specific lines, comparisons between *PDF1;GR* and *AN3;GR* cannot by themselves establish differences in the intrinsic responsiveness of the epidermal and subepidermal tissues.

In the weakest insertion (*PDF1;GR*^#4^), DEX induction produced only modest effects across the measured cellular parameters (Fig. S1D, G). Leaf area decreased by ∼17% (MOCK: 30.8 ± 8.8 mm²; DEX: 25.6 ± 9.3 mm²; Fig. S1H), and leaves retained visible surface ripples characteristic of the *jaw-D* background. Epidermal cell number decreased by ∼29% (MOCK: 8687 ± 2979; DEX: 6139 ± 2558; Fig. S1I), whereas subepidermal cell number decreased by only ∼12% (MOCK: 12412 ± 3476; DEX: 10874 ± 3059; Fig. S1J). Similarly, final cell size increased by ∼17% in the epidermis and ∼5% in the subepidermis (Fig. S2B, C and D). Together with the stronger responses observed in the other *PDF1;GR* insertions, these results show that the magnitude of the cellular and organ-level phenotypes varies with transgene strength (Tables S4 and S5).

In *AN3;GR*^#1^, upon DEX induction, subepidermal rTCP4 expression resulted in substantial reductions in cell number in both the epidermis (∼73%; MOCK: 10334 ± 7223; DEX: 2666 ± 1285; Fig. 2C) and subepidermis (∼62%; MOCK: 10823 ± 4105; DEX: 4111 ± 1202; Fig. 2D).

These reductions were accompanied by increases in final cell size in both layers, with epidermal cell size increasing by ∼59% (MOCK: 4042 ± 2711 µm²; DEX: 6434 ± 2793 µm²; Fig. 2H; S1E) and subepidermal cell size increasing by ∼8% (MOCK: 3859 ± 1303 µm²; DEX: 4172 ± 920 µm²; Fig. 2H; S1F). Thus, restricting rTCP4 induction to the subepidermis affected both subepidermal and epidermal cellular parameters (Tables S4 and S5).

In a second strong insertion, *AN3;GR*^#2^, DEX induction similarly rescued leaf area by ∼62% (MOCK: 37.1 ± 6.7 mm²; DEX: 14.2 ± 3.4 mm²; Fig. S3B). Cell number decreased by ∼72% in the epidermis (MOCK: 9434 ± 2022; DEX: 2588 ± 1070; Fig. S3D) and ∼74% in the subepidermis (MOCK: 14095 ± 4225; DEX: 3573 ± 1313; Fig. S3E). These reductions were accompanied by increases in final cell size in both layers, of ∼38% in the epidermis (MOCK: 3939 ± 154 µm²; DEX: 5419 ± 207 µm²; Fig. S4C, E) and ∼49% in the subepidermis (MOCK: 2636 ± 723 µm²; DEX: 3923 ± 1157 µm²; Fig. S4C, F) (Tables S4 and S5). Thus, an independent strong AN3;GR insertion reproduced the reduction in cell number and increase in cell size observed in both tissue layers.

In the weaker insertion *AN3;GR*^#3^, DEX induction produced an attenuated rescue of the *jaw-D* leaf phenotype (Fig. S3A, C), with leaf area decreasing by ∼48% (MOCK: 29.4 ± 5.5 mm²; DEX: 15.3 ± 3.8 mm²; Fig. S3B). Epidermal and subepidermal cell numbers decreased by ∼61% (MOCK: 8593 ± 2088; DEX: 3403 ± 1200; Fig. S3D) and ∼60% (MOCK: 12908 ± 3318; DEX: 5199 ± 1509; Fig. S3E), respectively. Final cell size increase by ∼32% in the epidermis (MOCK: 3422 ± 113 µm²; DEX: 4508 ± 1191 µm²; Fig. S4B, E) and ∼33% in the subepidermis (MOCK: 2227 ± 379 µm²; DEX: 2952 ± 391 µm²; Fig. S4B, F). Thus, the weaker AN3;GR insertion produced qualitatively similar cellular responses in both layers, but with reduced phenotypic severity compared with the stronger insertions (Tables S4 and S5).

In the weakest insertion, *AN3;GR*^#4^, DEX induction produced negligible effects on leaf growth (Fig. S5A, C), with leaf area decreasing by only ∼2% (MOCK: 24.6 ± 5.6 mm²; DEX: 23.5 ± 4. 6 mm²; Fig. S3B). Cellular responses were similarly limited: epidermal cell number increased slightly (∼2%; Fig. S3D), whereas subepidermal cell number decreased by ∼12% (Fig. S3E), and change in final cell size were small and non-significant (Fig. S4, E and F). Together with the progressively stronger responses observed in *AN3;GR*^#3^, *AN3;GR*^#2^, and *AN3;GR*^#1^, these results show that phenotypic severity varies with transgene strength (Tables S4 and S5).

The graded phenotypic responses observed across independent *PDF1;GR* and *AN3;GR* insertions indicate that the magnitude of rTCP4-induced effects is strongly influenced by transgene strength in both tissue contexts. Consequently, quantitative differences between individual epidermal and subepidermal lines cannot be attributed solely to the tissue in which rTCP4 is induced. Nevertheless, a qualitative difference was observed among the strongest insertions: strong epidermal induction of *PDF1;GR^#1^* and *PDF1;GR^#2^* completely arrested true-leaf development, whereas strong subepidermal induction in *AN3;GR*^#1^ and *AN3;GR*^#2^ produced small but viable leaves. This distinction suggests that high levels of TCP4 activity may have different developmental consequences depending on the tissue in which it is induced. However, quantitative measurements of rTCP4 abundance would be required to determine whether this reflects intrinsic differences in tissue sensitivity rather than differences in transgene expression.

Altogether, these results show that rTCP4 induction reduces cell number and promotes cell expansion both within the tissue in which it is induced and in adjacent tissue layers. The effect of epidermis- and subepidermis-specific induction across tissue compartments suggest that TCP4 can influence cellular behavior non-cell-autonomously, although the mechanism underlying these cross-layer effects remain unclear.

### 4. The confined expression of *TCP4* to the epidermis can rescue the *jaw-D* phenotype

The non-cell-autonomous effects of epidermal TCP4 could result from movement of the TCP4 protein into adjacent tissue layers. To test this possibility, we examined the effects of constitutive epidermal rTCP4 under the PDF1 promoter and compared mobile and mobility-restricted forms of the protein.

We generated multiple independent *pPDF1::rTCP4-mVENUS* lines in the *jaw-D* background (hereafter *PDF1-mVENUS*). Consistent with the severe phenotypes observed in the strongest inducible PDF1;GR lines (upon TCP4 induction in epidermis), constitutive epidermal rTCP4 expression strongly impaired development: ∼69% of seedlings failed to germinate, while a further ∼20-25% displayed severe developmental defects, including distorted or fused cotyledons and arrested leaf growth (Fig. S5E). We therefore selected the weakest viable insertions for further analysis (Table S1).

In *PDF1-mVENUS*^#1^, the area of the first leaf was significantly reduced compared with *jaw-D* (35.4 ± 9.9 mm² vs 18.3 ± 4.4 mm²; 48% reduction; Fig. 3A, B), accompanied by rescue of the rosette phenotype (Fig. S5F) This reduction in leaf area was associated with marked decreases in cell number in both the epidermis (9661 ± 4298 vs 2264 ± 1141; -76%; Fig. 3C) and subepidermis (16092 ± 6817 vs 4979 ± 2207; -69%; Fig. 3D). Conversely, mean cell area increased in both the epidermis (3665 ± 1248 µm² vs 8090 ± 3266 µm²; +120%; Fig. S5G, H) and subepidermis (2202 ± 382 µm² vs 3678 ± 778 µm²; +66%; Fig. S5G, I), showing that constitutive rTCP4 expression affects cellular growth parameters across both tissue layers (Tables S4 and S5).

**Fig. 3.**
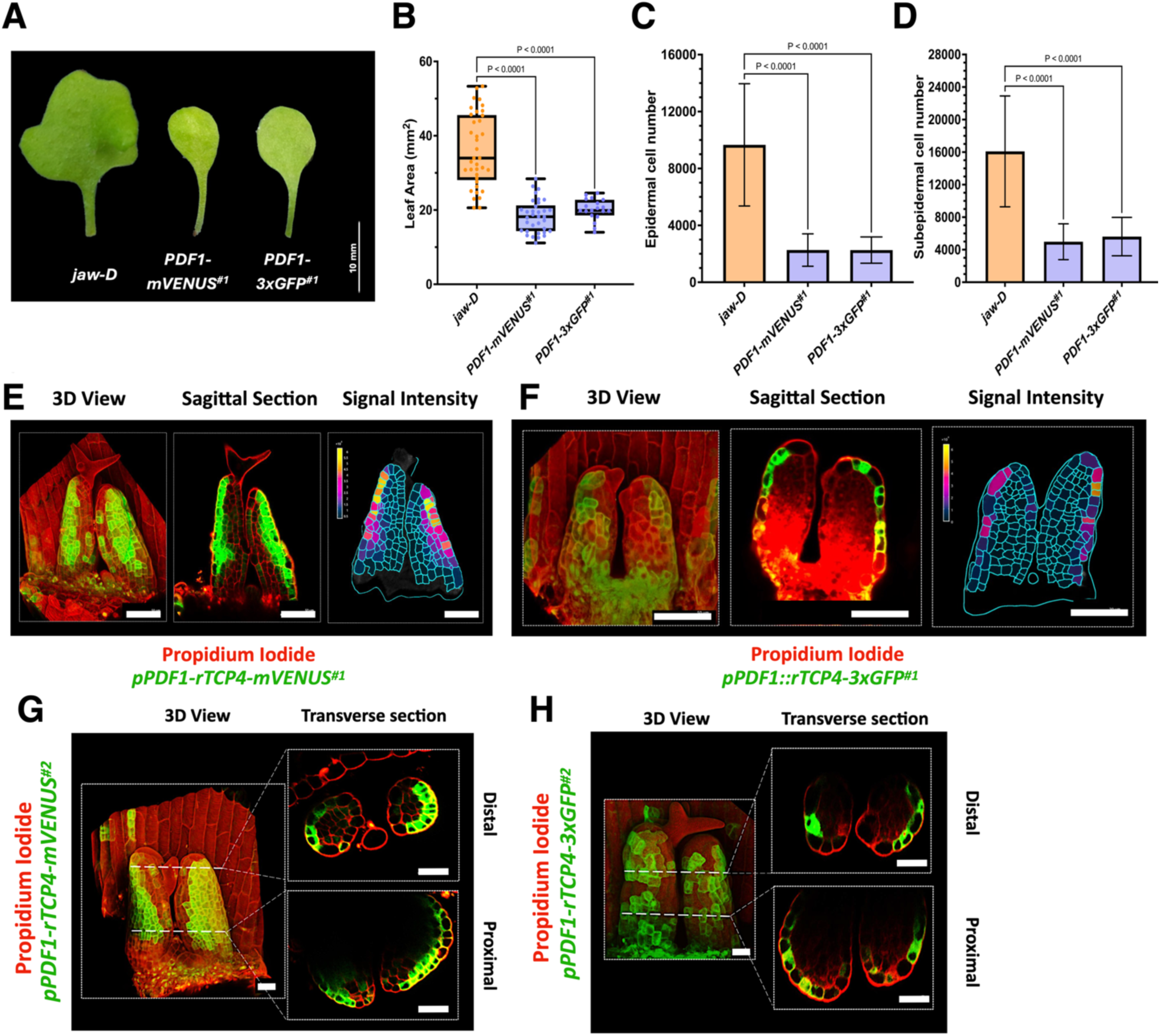
Epidermal TCP4 coordinates cross-layer growth through mobility-independent downstream signalling. (A) Representative images of the mature first leaf pair from 28-day-old plants of the indicated genotypes. Scale bar, 10 mm. (B) Quantification of leaf area (Y-axis) for the mature first leaf pair of the indicated genotypes (X-axis). N = 19-30 leaves. Statistical significance was assessed by one-way ANOVA followed by a Šídák post hoc test P-values indicated above the compared means). Error bars represent SD. (C, D) Estimated epidermal (C) and subepidermal (D) cell numbers for the indicated genotypes. Cell number was calculated as the ratio of total leaf area to mean cell area (8-10 leaves per genotype). Statistical analysis as in (B); error bars represent SD. (E, G) Confocal images of propidium iodide-stained leaf primordia (2-3 DAI) from two independent *pPDF1::rTCP4-mVENUS* insertion lines (#1 and #2). Three-dimensional views and optical mid-sagittal (E, left and middle panels) and transverse sections (G, distal and proximal) show fluorescence in the epidermis and adjacent subepidermal layers, indicating limited inter-layer mobility of TCP4-mVENUS. Scale bars in white, 50 µm. (F, H) Confocal images of propidium iodide-stained leaf primordia (2-3 DAI) from two independent *pPDF1::rTCP4-3xGFP* insertion lines (#1 and #2). Fluorescence is confined to the epidermis in three-dimensional views and optical sections (F, H), indicating restricted mobility of the TCP4-3xGFP fusion. Scale bars, 50 µm. (E, F) Signal intensity heat maps derived from the mid-sagittal sections of representative *pPDF1::rTCP4-mVENUS^#1^* (E, right panel) and *pPDF1::rTCP4-3xGFP^#1^* (F, right panel) primordia, illustrating fluorescence distribution across tissue layers. Warmer colours indicate higher signal intensity. Scale bars, 50 µm.

A second independent line, *PDF1-mVENUS*^#2^, showed a similar response. Leaf area was reduced by ∼49% (34.8 ± 7.7 mm² vs 17.9 ± 5.1 mm²; Fig. S5 A, B and F), accompanied by reductions in epidermal (∼ 72%; Fig. S5C) and subepidermal (∼ 69%; Fig. S5D) cell numbers. Mean cell area increased in both the epidermis (+93%) and subepidermis (+72%; Fig. S5G, J and K). Thus, two independent *PDF1-mVenus* insertions showed consistent effects on cell number and size across both tissue layers (Tables S4 and S5).

To determine whether TCP4 protein movement contributes to the observed cross-layer effects, we examined the spatial distribution of rTCP4-mVenus by confocal imaging using 3D reconstructions and transverse and sagittal optical sections. As controls, no fluorescence was observed in the WT leaves (Fig. S6C), whereas *p35S::GFP* showed fluorescence throughout all tissue layers (Fig. S6A). The *pPIN1::PIN1-GFP* showed signals in epidermis and vasculature/L3 (Abley et al., 2016; Fig. S6C) while *pPIN3::PIN3-GFP* was detected across all tissue layers (Le Gloanec et al., 2024; Fig. S6C). In contrast, *pPDF1::MBD-mCitrine* fluorescence was restricted to the epidermis (Trinh et al., 2024; Fig. S6B). In *PDF1-mVENUS* seedlings at 2-3 DAI, rTCP4-mVENUS fluorescence was detected not only in the abaxial epidermis, but also in underlying subepidermal layers in both sagittal (Fig. 3E; S6C, D) and transverse sections (Fig. 3G). MorphoGraphX-based signal quantification supported limited movement of rTCP4-mVenus across two to three cell layers (Fig. 3E, right panel). Given the ∼63 kDa size of the fusion protein, this limited intercellular movement is compatible with reported plasmodesmatal size-exclusion limits (Roberts & Oparka, 2003; Brunkard & Zambryski, 2017; Timney et al., 2016; Gundu et al., 2020). The patchy fluorescence pattern likely reflects the weak transgenic insertions required to maintain viability. Occasional fluorescence in both the adaxial and abaxial epidermis may similarly reflect variation in promoter activity and insertion-dependent expression strength.

These observations indicate that rTCP4 expressed in the epidermis can move into underlying tissues, providing a potential explanation for the cross-layer cellular effects observed in *PDF1- mVenus* lines. However, whether TCP4 protein movement is required for these non-cell-autonomous effects remain unclear. To address this, we generated lines expressing a mobility-restricted rTCP4-3xGFP fusion (*pPDF1::rTCP4-3xGFP*; hereafter *PDF1-3xGFP;* Table S1).

In *PDF1-3xGFP*^#1^, the area of the first leaf was reduced by ∼43% relative to *jaw-D* (35.4 ± 9.9 mm² vs 20.1 ± 2.2 mm²; -43%; Fig. 3A, B), accompanied by rescue of the rosette phenotype (Fig. S5F). Epidermal and subepidermal cell numbers decreased by ∼76% (Fig. 3C) and ∼17% (Fig. 3D), respectively, while mean cell area increased by ∼14% in the epidermis and ∼ 63% in the subepidermis (Fig. S5G, H and I). A second independent line, *PDF1-3xGFP*^#2^, showed a comparable reduction in leaf area (∼ 49%; Fig. S5A, B and F), together with reductions in epidermal (∼ 72%; Fig. S5C), and subepidermal (∼ 69%; Fig. S5D) cell numbers. Mean cell area increased in both the epidermis (∼ 146%; Fig. S5G, J and K) and subepidermis (∼ 71%; Fig. S5G, J and K). Thus, cellular responses across tissue layers persisted when rTCP4 was fused to a mobility-restricting 3xGFP tag (Tables S4 and S5).

Confocal imaging confirmed that *PDF1-3xGFP* fluorescence was restricted to the epidermis, with no detectable signal in underlying tissue layers in either sagittal (Fig. 3F; S6C, E) and transverse sections (Fig. 3H). MorphoGraphX-based signal quantification further supported the epidermis-restricted localisation of the fusion protein (Fig. 3F, right panel).

Altogether, these findings show that epidermal TCP4 is sufficient to alter cell proliferation and expansion across multiple tissue layers. Importantly, comparable cross-layer cellular responses were observed when rTCP4 was mobile or restricted to the epidermis, indicating that TCP4 protein movement is not required for its non-cell-autonomous effects. Instead, TCP4 activity in the epidermis must influence neighbouring tissues through downstream intercellular signals, which could involve biochemical and/or mechanical coupling between tissue layers. Therefore, we next examined how strong epidermal TCP4 activity affects the progression of leaf growth and morphogenesis.

### 5. Epidermal TCP4 allows normal primordium initiation but suppresses proliferation and locks leaf primordia in an axisymmetric state

To investigate the developmental consequences of strong epidermal TCP4 activity, we selected the two strongest inducible lines, *PDF1;GR^#1^* and *PDF1;GR^#2^*(Fig. 2A; S1D), for further analysis. Constitutive lines producing similarly severe phenotypes could not be used because they failed to survive beyond early developmental stages. In contrast, weaker viable lines, including *PDF1;GR^#3^* (Fig. 2A) and the constitutive *PDF1-mVENUS^#^*^1^ line (Fig. 3A), showed only partial phenotypic rescue. The strong inducible lines therefore provide a system to examine the pronounced developmental effects of epidermal TCP4 activity while retaining control over the timing of its induction.

To determine when strong epidermal TCP4 activity begins to affect leaf development, we examined *PDF1;GR*^#1^ primordia by scanning electron microscopy at 1 and 3 days after initiation (DAI) following continuous MOCK or DEX treatment. At 1 DAI, leaf primordia were present under both conditions and were comparable in length (50-60 µm; Fig. 4A), indicating that epidermal TCP4 induction did not prevent primordium initiation. By 3 DAI, however, their growth had markedly diverged: MOCK-treated primordia had elongated by a further 100-150 µm, reaching 150-210 µm in length, whereas DEX-treated primordia remained only 60-70 µm long (Fig. 4A, B). Thus, strong epidermal TCP4 activity permits leaf primordium initiation but severly restricts subsequent organ growth.

**Fig. 4.**
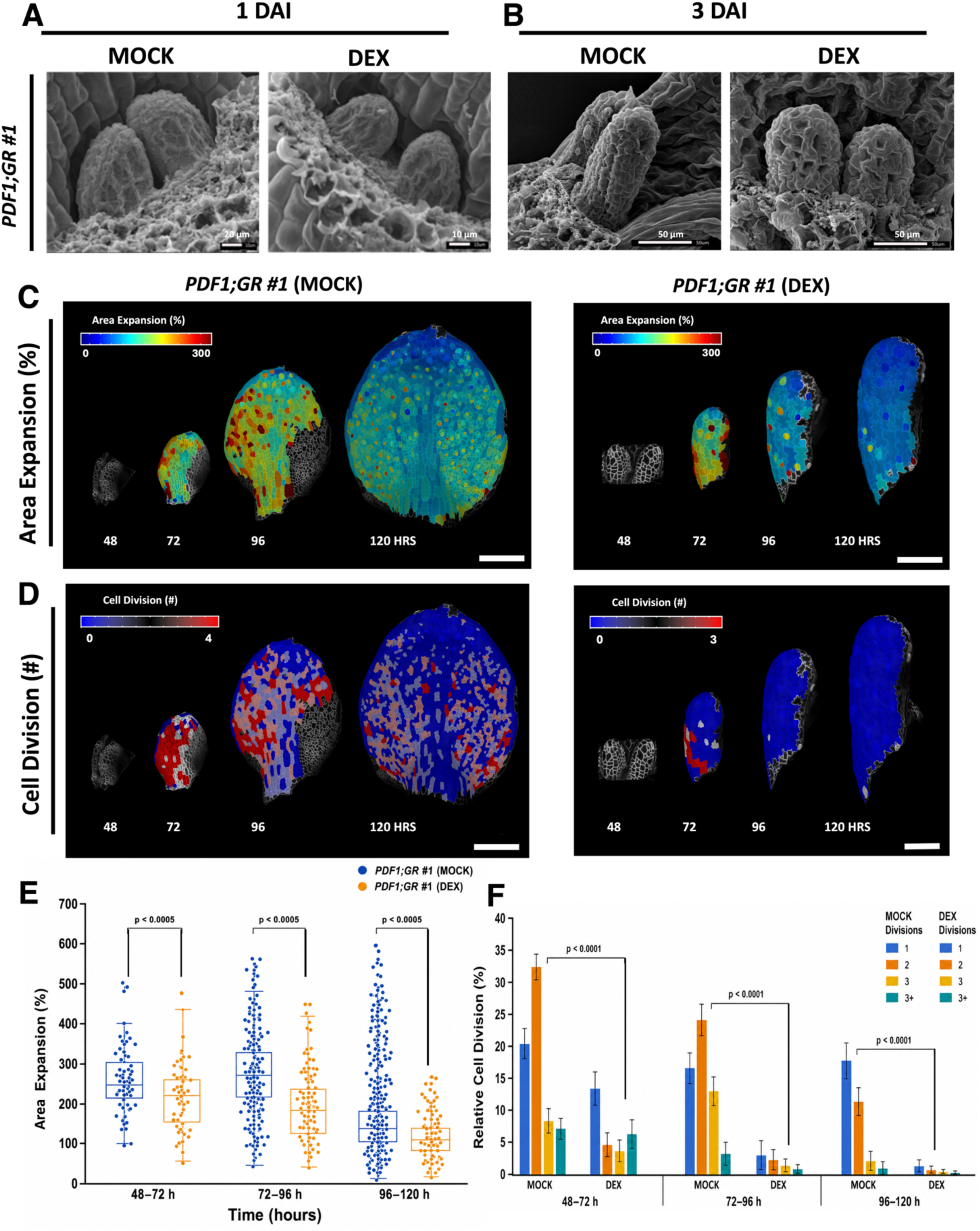
Epidermal TCP4 induction promotes precocious differentiation and suppresses growth dynamics during leaf development. (A, B) Scanning electron micrographs of leaf primordia from the inducible *pPDF1::rTCP4-GR* #1 (*PDF1;GR*#1) line under MOCK and continuous dexamethasone (DEX; 12 μM) treatment at 1 and 3 days after initiation (DAI). N = 4-6 primordia. Scale bars, 10 µm (A) and 50 µm (B). MOCK-treated primordia display normal outgrowth at 3 DAI, whereas DEX-treated primordia exhibit reduced lamina expansion following epidermal TCP4 induction. (C) Heat maps showing relative area expansion (%) in *PDF1;GR*#1 crossed to *pUBQ10::Myr-YFP* (plasma membrane marker) under MOCK and DEX conditions. Primordia were live-imaged from 48 to 120 hours after initiation. Scale bar, 100 µm. (D) Heat maps depicting the spatial distribution of cell division events across successive time intervals during live imaging under MOCK and DEX treatments (48-120 hours after initiation). Scale bar, 100 µm. (E) Box plots quantifying area expansion from three independent leaf primordia of the *PDF1;GR*#1 line under MOCK (blue) and DEX (orange) conditions across the time course. Statistical comparisons between MOCK and DEX treatments at corresponding time points were performed using a two-tailed Student’s t-test; P values are indicated above the box plots. (F) Grouped bar graphs showing the relative frequency of cell division events (percentage of cells dividing once, twice, or more) across time points under MOCK and DEX treatments. Colour legends indicate the number of divisions per cell. Statistical comparisons between MOCK and DEX treatments (grouped mean) at corresponding time points were performed using a two-tailed Student’s t-test; P values are indicated above the box plots.

To determine the cellular basis of this growth arrest, we performed live confocal imaging of the leaf epidermis at 24-hour intervals throughout early leaf development. *PDF1;GR*^#1/2^ were crossed with the plasma membrane reporter *pUBQ10::myr-YFP* (Willis et al., 2016) to visualize cell boundaries, and F1 leaf primordia were imaged from 2 to 5 DAI (48-120 hrs) under continuous MOCK or DEX treatment. Using MorphoGraphX, we then quantified cellular growth, cell division patterns, and cell size over time (Kuchen et al., 2012; Barbier de Reuille et al., 2015; Kierzkowski et al., 2019; Strauss et al., 2022).

At the organ level, MOCK-treated leaf primordia expanded substantially over time, reaching ∼600 µm in length and ∼320 µm in width by 5 DAI (Fig. 4C, D). Although the *jaw-D* background produced moderate negative Gaussian curvature at the leaf tip, the developing lamina underwent overall blade flattening. In contrast, DEX-treated primordia reached ∼500 µm in length but only ∼150 µm in width (Fig. 4C, D). Consequently, DEX-treated primordia retained a narrow, approximately axisymmetric morphology rather than developing a flattened lamina. These observations indicate that TCP4 activity preferentially restricts mediolateral expansion during blade development.

To determine how these morphological differences arise from changes in growth dynamics, we quantified cellular area expansion over successive 24-hour intervals expressed at the percentage increase in cell area increase independently of cell divisions (Kierzkowski et al., 2019). During the 48-72 h, both MOCK- and DEX-treated primordia showed high area expansion across the developing lamina, with elevated growth at the margins (Fig. 4C, E). However, the spatial distribution of growth already differed between treatments: in DEX-treated primordia, regions of high area expansion were shifted proximally, while the distal tip entered a slow-growth phase earlier than in MOCK. Despite these spatial differences, mean area expansion remained comparable between treatments at this stage (∼200%; Fig. 4E).

During the 72-96 h interval, MOCK-treated primordia showed reduced area expansion at the distal tip while growth remained high across the more proximal the lamina, establishing a clear basipetal growth gradient as the blade flattened (∼200% mean area expansion; Fig. 4C, E). In contrast, DEX-treated primordia exhibited a marked reduction in area expansion across the organ, with mean values falling to ∼100%, approximately half those of MOCK (Fig. 4C, E). By 96-120 h, MOCK-treated leaves maintained a pronounced basipetal gradient, with growth concentrated proximally and mean area expansion remaining at ∼175%. DEX-treated leaves showed further growth attenuation, with a broad slow-growth domain extending across the lamina and mean area expansion decreasing to ∼69% (Fig. 4C, E). Together, these growth maps show that epidermal TCP4 induction causes an earlier distal decline in growth followed by a strong attenuation of growth across the developing lamina.

Cell division patterns showed a similarly pronounced divergence between treatments. MOCK-treated primordia underwent sustained cell division throughout the 48-120 h imaging period, with divisions following a basipetal distribution and becoming progressively restricted to the proximal region of the expanding lamina (Fig. 4D, F). In contrast, divisions in DEX-treated primordia were largely confined to the 48–72 h interval, with virtually no proliferation detected thereafter (Fig. 4D, F). Thus, strong epidermal TCP4 activity causes a premature cessation of cell proliferation, consistent with an accelerated transition from proliferation to differentiation.

The premature decline in cell proliferation was accompanied by a pronounced increase in cell size. Under MOCK conditions, mean cell area increased gradually during primordium development, from ∼120-150 µm² at 48 h to ∼250-300 µm² at 120 h (Fig. S7A, C). In DEX-treated, cell enlargement occurred substantially earlier and reached much greater values. At 48 hours, mean cell area was already slightly higher than in MOCK (∼150-180 µm²), and by 72 h, had increased to ∼350-450 µm², approximately twice the ∼150-200 µm² observed in MOCK (Fig. S7A, C). The difference increased further at 96 h, when DEX-treated cells reached ∼600- 800 µm² compared with ∼200-250 µm² in MOCK (Fig. S7A, C). By 120 h, DEX-treated cells exceeded ∼1000 µm², with mean cell area reaching ∼800-1200 µm², whereas MOCK-treated cells remained largely below ∼300 µm² (Fig. S7A, C). Overall, epidermal TCP4 induction resulted in an approximately three- to four-fold increase in mean cell area over the course of development. Because this pronounced cell enlargement coincided with premature proliferation arrest and an overall reduction in area expansion, it is more consistent with the accumulation of enlarged, non-dividing cells following early proliferative exit than with enhanced compensatory growth. Spatial heat maps further showed that cell enlargement emerged first distally and progressed basipetally, consistent with an earlier progression of differentiation along the proximodistal axis.

To determine how epidermal TCP4 induction affects the directionality of growth, we quantified growth anisotropy and the principal directions of growth (PDGs), which describe complementary properties of the local deformation field. Growth anisotropy reflects the relative difference between the maximal and minimal principal growth components, with values approaching 1 indicating more isotropic growth and higher values indicating increasingly anisotropic growth, whereas PDGs describe the orientation of the maximal growth component across the tissue (Strauss et al., 2022). Under MOCK conditions, cellular growth anisotropy progressively increased during lamina outgrowth, reaching ∼1.5-1.7 at 72 h and ∼1.6-1.8 at 96 h (Fig. S7B, D). Following DEX induction, anisotropy transiently increased at 72 h (∼1.8-2.0), before declining to values comparable to MOCK at 96 h (∼1.4-1.6). By 120 h, anisotropy decreased further in DEX-treated primordia (∼1.2-1.4), indicating that late cellular expansion became increasingly isotropic (Fig. S7B, D).

Analysis of PDG orientation revealed a distinct difference in the spatial organisation of growth. Despite the decline in cellular growth anisotropy at later stages, PDGs remained spatially coherent in DEX-treated primordia, with the maximal growth component predominantly align along the proximodistal axis (Fig. S7B). Thus, the increasingly isotropic growth of individual cells was not accompanied by a loss of tissue-level directional organisation: although the two principal growth components became more similar in magnitude, their maximal component remained coherently oriented across neighbouring cells. In contrast, MOCK-treated primordia displayed more heterogeneous PDG orientations across the lamina, reflecting greater spatial diversification of local growth directions during blade expansion. Collectively, these findings show that strong epidermal TCP4 activity profoundly alters the spatiotemporal organisation of leaf growth rather than simply reducing its magnitude. TCP4 induction promotes premature proliferative exit, followed by the accumulation of enlarged, non-dividing cells and an earlier basipetal progression of growth attenuation. These changes are accompanied by dynamic alterations in cellular growth anisotropy and sustained spatial coherence of proximodistally oriented PDGs, while mediolateral expansion of the primordium remains strongly restricted. Consequently, DEX-treated primordia fail to undergo normal blade flattening and retain a narrow, approximately axisymmetric morphology. Notably, these pronounced developmental effects were observed in F1 seedlings carrying a single copy of the *PDF1;GR* transgene is present, highlighting the strong response to epidermal TCP4 activity.

### 6. Temporal induction reveals a pronounced and persistent response to epidermal TCP4 activity

To assess the temporal sensitivity of developing leaves to TCP4 activity, we transferred *jaw-D;GR* and *PDF1;GR^#1^* seedlings between non-inductive (MOCK) and inductive (DEX) media at 4 days after stratification (DAS), generating four treatment regimes: MOCK-to-MOCK (MtM), DEX-to-DEX (DtD), MOCK-to-DEX (MtD), and DEX-to-MOCK (DtM). To directly assess the temporal sensitivity of developing leaves to TCP4 activity, we transferred seedlings of *jaw-D;GR* and *PDF1;GR^#1^* between non-inductive (MOCK) and inductive (DEX) media at 4 days after stratification (DAS), generating four treatment regimes: MOCK-to-MOCK (MtM), DEX-to-DEX (DtD), MOCK-to-DEX (MtD), and DEX-to-MOCK (DtM). We selected 4 DAS because the first leaf pair remains actively proliferating at this stage, whereas proliferation becomes progressively restricted during subsequent development (Challa et al., 2019). This experimental design allowed us to determine whether TCP4 induction initiated during this proliferative window remains sufficient to alter subsequent leaf growth and whether removal of the inducing signal permits developmental recovery.

In the *jaw-D;GR* control, the continuous-treatment conditions (MtM and DtD) produced the expected divergence (Fig. S8A, E). Continuous DEX treatment significantly reduced leaf area by ∼67% relative to continuous MOCK (MtM: 48.2 ± 7.5 mm²; DtD: 15.4 ± 3.9 mm²; Fig. S8A, C). This reduction was accompanied by decreases in cell number in both the epidermis (∼9%; Fig. S8G) and subepidermis (∼72%; Fig. S8H) cell number, together with increase in final cell area of ∼41% and ∼22%, respectively (Fig. S9A, C, D; Tables S6, S7). Transfer from MOCK to DEX at 4 DAS (MtD) produced a similarly strong phenotype, reducing leaf area by ∼65% relative to MtM (MtD: 17.0 ± 4.8 mm²; Fig. S8A, C, E). Epidermal and subepidermal cell numbers were reduced by ∼76% and ∼66%, respectively (Fig. S8G, H; Tables S6, S7), accompanied by increases in final cell area in both layers (Fig. S9A, C, D; Tables S6, S7). Notably, the reduction in leaf area was comparable to that observed under continuous DEX treatment, indicating that TCP4 induction initiated at 4 DAS is sufficient to produce near-maximal effects on organ size under these conditions.

By contrast, transfer from DEX to MOCK at 4 DAS (DtM) resulted in partial recovery of leaf growth, although final leaf area remained ∼47% lower than in MtM controls (DtM: 25.4 ± 3.9 mm²; Fig. S8A, C, E). Cell-number reductions were intermediate between the continuous MOCK and DEX conditions, reaching ∼68% in the epidermis and about ∼41% in the subepidermis (Fig. S8G, H; Tables S6, S7). Final cell area increased by ∼43% in the epidermis but only ∼4% in the subepidermis (Fig. S9A, C, D; Tables S6, S7). Thus, removal of DEX at 4 DAS partially alleviated the effects of earlier TCP4 induction, indicating that the developmental response remained at least partially reversible at this stage.

We next performed the same transfer experiments in *PDF1;GR^#1^*to determine whether the effects of strong epidermal TCP4 induction remained reversible at 4 DAS. Continuous DEX treatment (DtD) resulted in a complete failure of leaf development, with no detectable outgrowth beyond cotyledon-stage structures (Fig. S8B, F). Strikingly, seedlings transferred from DEX to MOCK at 4 DAS (DtM) showed a similarly severe phenotype, with no subsequent leaf outgrowth despite removal of the inducing signal. This contrasts with *jaw-D;GR*, in which the same DtM treatment permitted partial recovery, indicating that the developmental effects of strong epidermal TCP4 induction persist after DEX removal at 4 DAS.

In *PDF1;GR^#1^*, induction initiated at 4 DAS (MtD) strongly restricted subsequent leaf development, reducing final leaf area by ∼81% relative to MtM (MtM: 42.8 ± 7.4 mm²; MtD: 7.3 ± 2.5 mm²; Fig. S8B, D). This was accompanied by pronounced reductions in cell numbers in both the epidermis (∼88%) and subepidermis (∼87%; Fig. S8I, J), together with increases in final cell area of ∼84% and ∼57%, respectively (Fig. S9B, E, F; Tables S6, S7). The reduction in leaf area was greater than that observed following MtD treatment of *jaw-D;GR* (∼65%), as were the reductions in epidermal and subepidermal cell number (∼76% and ∼66%, respectively).

Together, the live-imaging and transfer experiments show that strong epidermal TCP4 activity rapidly suppresses cell proliferation and induces developmental changes that persist after removal of DEX at 4 DAS. These effects extend across tissue layers, resulting in pronounced reductions in both epidermal and subepidermal cell number. The concomitant increase in final cell size, despite the reduced cellular area expansion observed by live imaging, is consistent with premature exit from proliferation and the accumulation of enlarge, non-dividing cells rather than enhanced cellular growth rates. Thus, TCP4 induction in the epidermis during an early proliferative window is sufficient to impose a persistent, organ-wide restriction of leaf growth.

Notably, induction from the epidermal PDF1 promoter produced a more severe and less reversible developmental phenotype than induction from the endogenous TCP4 promoter under the conditions analysed. However, because TCP4 abundance was not quantitatively matched between the *PDF1;GR* and *jaw-D;*GR lines, these differences cannot be attributed solely to the spatial domain of TCP4 induction. Nevertheless, the strong and persistent response to PDF1-driven TCP4 demonstrates that epidermal TCP4 activity is sufficient to profoundly alter leaf development across tissue layers. Together with the effects observed when TCP4 mobility was restricted to the epidermis, these results further support a non-cell-autonomous influence of epidermal TCP4 activity on leaf morphogenesis.

These findings raise the question of how epidermal TCP4 activity is transmitted across tissue layers and translated into the altered growth geometry of developing primordia. Because the epidermis integrates biochemical and mechanical cues that contribute to organ morphogenesis, we hypothesised that TCP4-induced changes in epidermal growth could modify the mechanical coordination of the tissue. In particular, the persistent proximodistal organisation of growth and restricted mediolateral expansion observed following TCP4 induction suggested that changes in growth directionality might involve altered cytoskeletal organisation and cell wall mechanics. We therefore examined cortical microtubule (CMT) orientation, a key regulator of anisotropic cell expansion, together with epidermal cell wall mechanics under MOCK and DEX conditions (Hamant et al., 2008).

### 7. Epidermal TCP4 induction promotes CMT co-alignment to reinforce directional growth

Given the altered growth directionality observed following epidermal TCP4 induction, we next examined cortical microtubules (CMT) organisation during early leaf development. To visualise CMTs, we crossed *PDF1;GR*^#1^ with the microtubule reporter *p35S::MBD-GFP* (Hamant et al., 2008) and performed high-resolution Airyscan confocal imaging of first –leaf primordia at 3 DAI under continuous MOCK or DEX treatment. In MOCK-treated primordia, CMTs displayed heterogeneous orientations across the epidermis, with transverse and oblique arrays present in both distal and proximal regions (Fig. 5A). At the cellular level, neighbouring cells frequently exhibited different dominant CMT orientations (Fig. 5A, zoomed insets), revealing limited tissue-scale co-alignment. This heterogeneous organisation parallels the spatially diverse PDGs orientations observed in MOCK-treated primordia during lamina expansion (Fig. S7B).

**Fig. 5.**
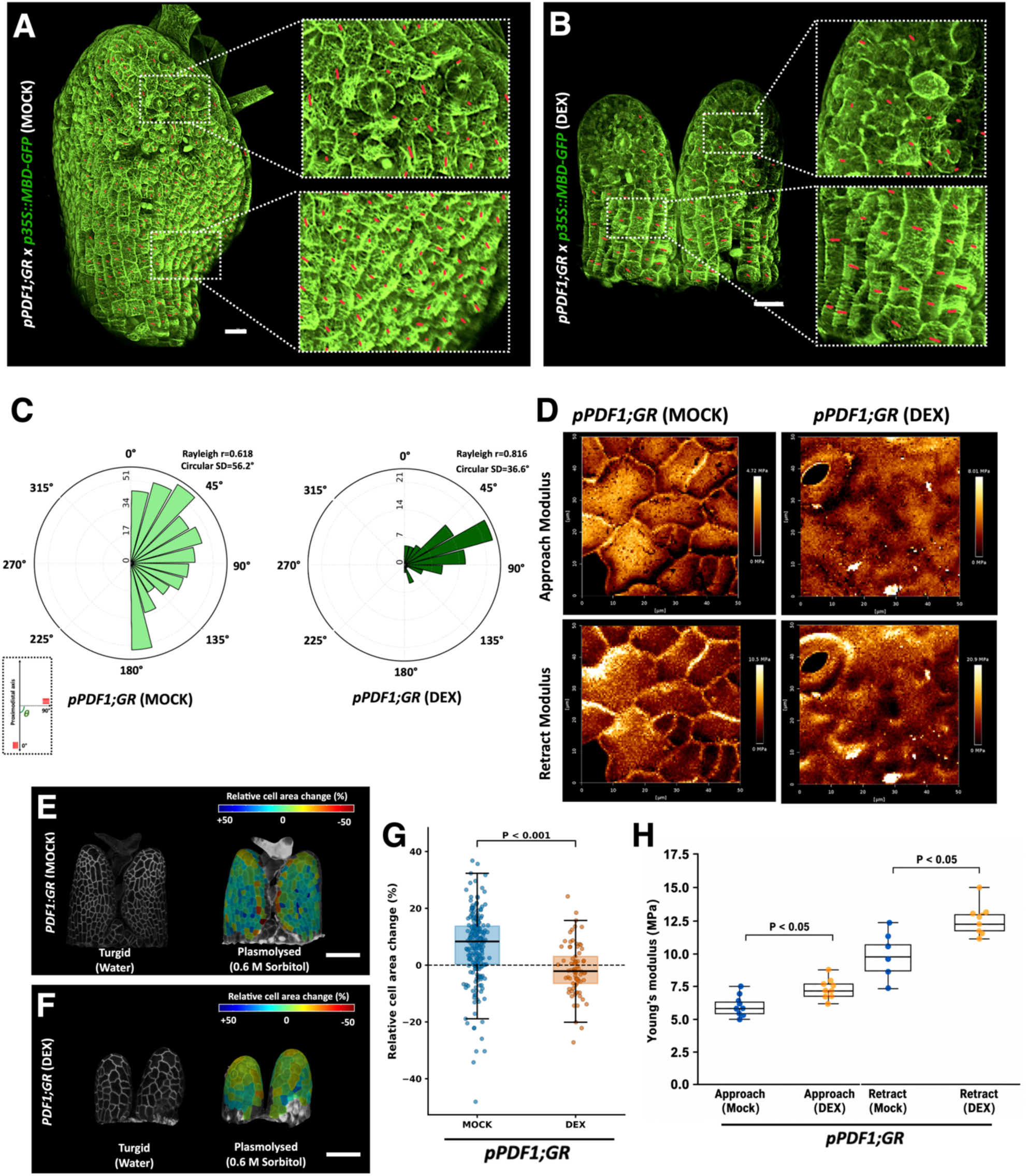
TCP4-mediated molecular signals reprogram epidermal mechanical cues to coordinate leaf development. (A, B) Airyscan confocal images of first leaf primordia from the inducible *pPDF1::rTCP4-GR* line crossed to *pPDF1::MBD-mCitrine* (cortical microtubule marker) at 3 days after initiation (DAI) under MOCK (A) and dexamethasone (DEX) (B) conditions. Images show the entire primordium with zoomed views of distal and proximal regions. Red lines indicate dominant cortical microtubule orientation in each cell, quantified using FibrilTool. Scale bars, 20 µm. (C) Rose plots showing distributions of cortical microtubule orientations from three independent biological replicates, quantified using FibrilTool, under MOCK (light green) and DEX (dark green) conditions. Plots show microtubule angle distributions (-90° to +90°, projected as 0° to 180°), with bar lengths corresponding to the number of cells exhibiting each orientation (n = 120-1240 cells). Both conditions show significant non-random alignment (Rayleigh test, p < 10⁻²⁵). DEX-treated primordia display increased directional coherence (r = 0.816) and reduced angular dispersion (circular SD = 36.6°) compared with MOCK (r = 0.618; SD = 56.2°). The inset schematic illustrates microtubule orientation relative to the proximodistal axis (0-180°) and the orthogonal mediolateral axis (90°). (D) Representative Atomic Force Microscopy (AFM) stiffness maps of epidermal cells (50 × 50 µm² regions) from *pPDF1::rTCP4-GR* leaves at 3 DAI under MOCK and DEX treatments. Heat maps show Young’s modulus values obtained during approach (upper) and retract (lower) indentation phases. Colour scales indicate stiffness values in MPa. (E, F) Representative heat maps of Relative cell area shrinkage of epidermal cells projected on plasmolysed samples from 2 DAI *pPDF1::rTCP4-GR* seedlings under MOCK (E) and DEX (F) treatments in fully turgid (water) and plasmolysed conditions (0.6 M sorbitol, 30 min). Scale bars, 50 µm. (G) Relative change in projected cell area following sorbitol treatment in MOCK- and DEX-treated samples, calculated for matched cells as relative cell area change (%). Positive and negative values indicate a decrease and an apparent increase in area, respectively. The dashed line indicates zero change. Boxes show the interquartile range, centre lines the median, and whiskers 1.5× the interquartile range; points represent individual cells (MOCK, n = 224; DEX, n = 77). Welch’s t-test, P < 0.001. (H) Box plots showing mean apparent Young’s modulus values derived from AFM approach and retract measurements across 4-6 independent leaf primordia (50 × 50 µm² sectors per leaf) under MOCK and DEX treatments. Statistical significance between treatments was assessed using a two-tailed Student’s t-test, with p-values indicated above comparisons.

**Fig. 6.**
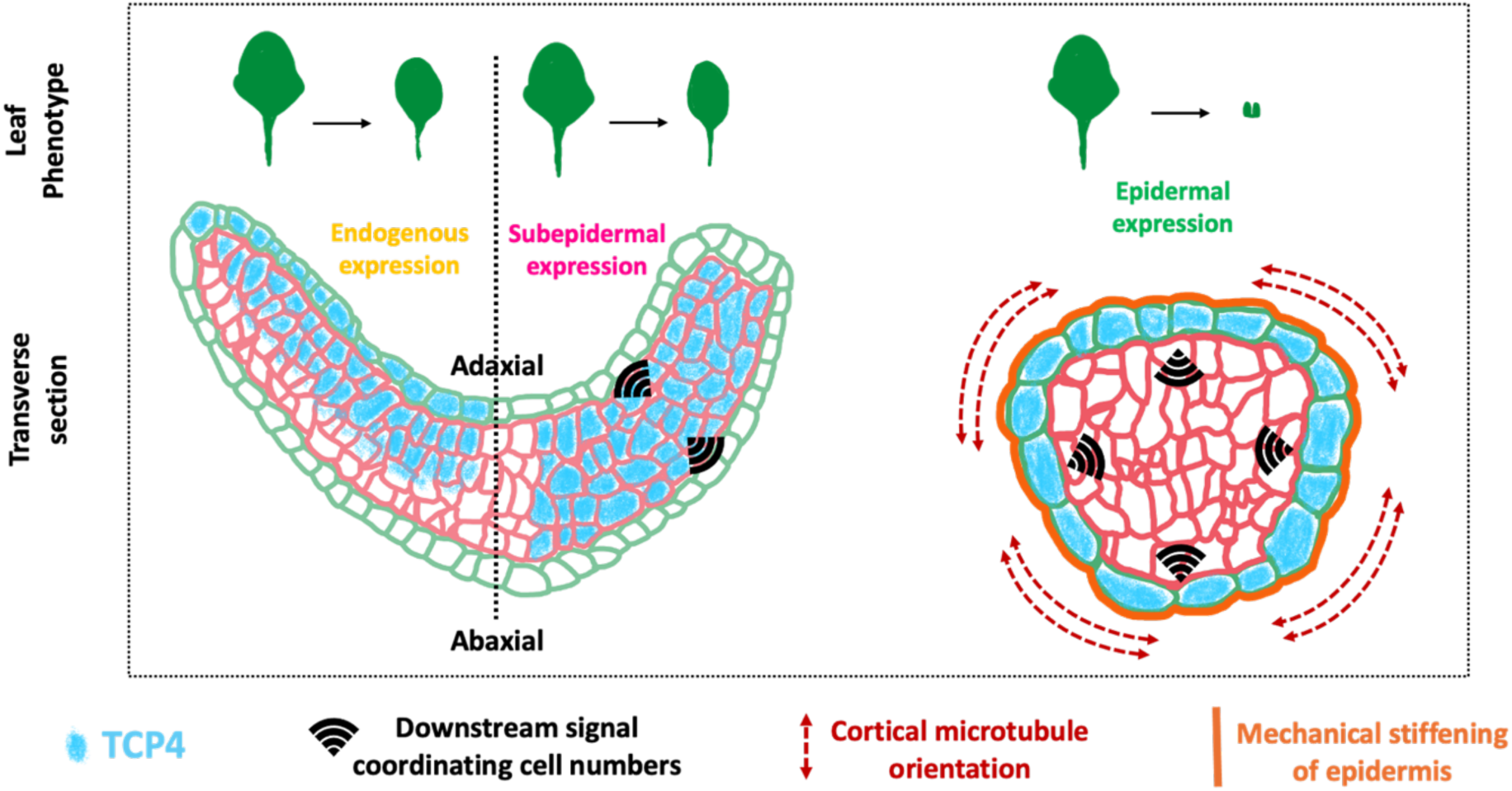
Working model of TCP4-mediated interlayer communication coordinating robust leaf shape and size through genetic and mechanical signalling. Schematic representation illustrating how layer-specific activity of TCP4 regulates leaf morphogenesis by coordinating genetic regulation, biochemical cell wall remodelling, and tissue mechanics across the developing primordium. *Top panels:* Representative leaf phenotypes resulting from distinct spatial domains of TCP4 expression. *Left panel (Endogenous expression):* Native TCP4 activity across the developing leaf coordinates proliferation between the epidermis and inner tissues. In transverse sections, balanced cell numbers across layers maintain laminar integrity and produce a normally shaped leaf. *Middle panel (Subepidermal expression):* Subepidermal induction of TCP4 similarly restores leaf morphology by coordinating cell proliferation across adjacent layers through non-cell-autonomous signalling. *Right panel (Epidermal expression):* Epidermis-restricted TCP4 activity primarily modulates tissue mechanics. TCP4 induction reorganises cortical microtubules, promotes cell wall reinforcement, and increases epidermal stiffness. This stiffened epidermal layer mechanically constrains underlying tissues and influences organ growth patterns. At the molecular level, TCP4 induction sustains auxin signalling that promotes differentiation. At the biochemical level, this signalling coordinates layer-specific pectin methylesterification, with increased methylesterified pectin in the epidermis enhancing cell wall rigidity and demethylesterified pectin in subepidermal layers transmitting mechanical constraints non-cell-autonomously. At the cytoskeletal level, cortical microtubules align in transverse arrays that restrict mediolateral expansion and reinforce proximodistal growth. Together, these molecular, biochemical, cytoskeletal, and biophysical processes elevate cell wall stiffness and generate a rigid epidermal corset that channels growth directionality across the leaf primordium, thereby ensuring robust control of organ shape and size.

In contrast, DEX-treated primordia displayed predominantly transverse CMT arrays across the epidermis (Fig. 5B). This organisation was apparent in both distal and proximal regions, where CMTs were preferentially oriented perpendicular to the proximodistal axis and showed greater alignment between neighbouring cells than under MOCK conditions (Fig. 5B, zoomed insets). Thus, epidermal TCP4 induction is associated with increased tissue-scale CMT co-alignment and a predominantly transverse orientation.

To quantify differences in CMT organisation, we measured CMT orientation using FibrilTool (Boudaoud et al., 2014) across three independent biological replicates (n = 120-1240 cells per condition). CMT orientations were significantly non-random under both MOCK and DEX conditions (Rayleigh test, p < 10⁻²⁵; Fig. 5C). However, DEX-treated primordia showed substantially greater directional coherence, with a higher mean resultant length (Rayleigh r = 0.816) and lower circular standard deviation (36.6°) than MOCK-treated primordia (r = 0.618; circular SD = 56.2°; Fig. 5C). Consistent with this increased co-alignment, most CMT arrays in DEX-treated cells were oriented within ±45° of the transverse axis, whereas MOCK-treated cells displayed a broader distribution of orientations (Fig. 5C).

The increased transverse co-alignment of CMTs in DEX-treated primordia is consistent with the sustained proximodistal PDG alignment and restricted mediolateral expansion observed by live imaging (Fig. 4, S7). Because CMT organisation influences the orientation of cellulose microfibril deposition and, consequently, anisotropic cell wall properties, the predominance of transverse CMT arrays could favour proximodistal over mediolateral expansion. Thus, the tissue-scale CMT co-alignment induced by epidermal TCP4 provides a potential cellular basis for the directional growth bias associated with the narrow, axisymmetric morphology of DEX-treated primordia.

### 8. Epidermal TCP4 induction alters pectin methylesterification patterns across tissue layers

In addition to cellulose organisation, pectin composition and methylesterification can strongly influence the mechanical properties of the cell wall. The consequences of pectin demethylesterification are context dependent: de-esterified homogalacturonan can undergo Ca^2+^-mediated cross-linking, potentially increasing wall rigidity, but can also become susceptible to pectin-degrading enzyme that promote wall remodelling (Micheli, 2001; Wolf et al., 2009; Jolie et al., 2010; Hocq et al., 2017). To determine whether epidermal TCP4 induction is associated with changes in pectin methylesterification, we performed immunolabelling on sections of 6-DAI leaves from Col-0 and *PDF1;GR* seedlings grown under MOCK or DEX conditions, using LM20 and LM19 antibodies to detect highly methylesterified and low-methylesterified /de-esterified homogalacturonan epitopes, respectively.

In Col-0 and MOCK-treated *PDF1;GR* controls, LM20 labelling was relatively uniformly across cell walls in both the epidermal and subepidermal layers, with moderate signal intensity throughout the tissue (Fig. S10A, left and middle panels). LM19 was weak and diffuse across both tissue layers, with similar patterns observed in Col-0 and MOCK-treated *PDF1;GR* leaves (Fig. S10B, left and middle panels).

DEX-treated *PDF1;GR* primordia displayed pronounced, layer-specific changes in pectin labelling. LM20 signal was increased along epidermal cell walls compared with MOCK-treated controls (Fig. S10A, C, arrows), indicating an increased abundance or accessibility of highly methylesterified homogalacturonan epitopes in the TCP4-expressing layer. In contrast, LM19 labelling was enhanced predominantly in the subepidermal layer of DEX-treated primordia (Fig. S10B, C, right panel, arrows), a pattern not observed in either Col-0 or MOCK-treated *PDF1;GR* controls. Thus, epidermal TCP4 induction is associated with distinct changes in pectin methylesterification-related epitopes in the epidermal and subepidermal layers.

The complementary, layer-specific changes in LM20 and LM19 labelling indicate that epidermal TCP4 induction differentially alters pectin organisation across tissue layers. Increased LM20 labelling in the epidermis is consistent with an enrichment of highly methylesterified homogalacturonan epitopes, whereas the increased LM19 signal in the subepidermis indicates greater abundance or accessibility of low-methylesterified/de-esterified homogalacturonan epitopes. However, the mechanical consequences of these changes cannot be inferred directly from immunolabelling. Demethylesterified homogalacturonan can undergo Ca²⁺-mediated cross-linking, which may promote wall stiffening, but can also become susceptible to polygalacturonase- and pectate lyase-mediated degradation and wall remodelling (Wolf et al., 2009; Jolie et al., 2010). Moreover, LM19 and LM20 labelling can be influenced by epitope accessibility and overall wall architecture in addition to pectin methylesterification status. We therefore interpret these results as evidence that TCP4 induction is associated with layer-specific changes in pectin composition or organisation, rather than as direct evidence of altered wall stiffness or mechanical transmission between tissue layers.

### 9. Epidermal TCP4 induction increases cell wall stiffness

To determine whether the cytoskeletal and pectin changes associated with epidermal TCP4 induction are accompanied by altered cell wall mechanical properties, we performed atomic force microscopy (AFM) on epidermal cells of *PDF1;GR* leaf primordia at 3 DAI under MOCK and DEX conditions (Peaucelle et al., 2011; Routier-Kierzkowska & Smith, 2013; Bidhendi & Geitmann, 2018; Koyama et al., 2025). Apparent Young’s modulus was measured over 50 × 50 µm² regions of individual leaves (Fig. 5D) from both approach and retract force curves. The approach modulus reports the resistance of the cell surface during loading, whereas the retract modulus characterises the mechanical response during unloading and can additionally reflect elastic recovery, viscoelastic behaviour, and tip-sample interactions (Vinckier & Semenza, 1998). Together, these measurements provide complementary estimates of the mechanical response of the epidermal cell wall.

In MOCK-treated primordia, AFM heat maps revealed spatial variation in apparent Young’s modulus across the epidermis (Fig. 5D, left panels). The approach measurements yielded values of ∼6-7 MPa, whereas the retract measurements were higher, at ∼8-9 MPa. Thus, the retract modulus exceeded the approach modulus by approximately 2 MPa under MOCK conditions. This difference between loading and unloading responses is consistent with the viscoelastic and adhesive behaviour expected for hydrated biological materials, although these measurements do not distinguish the individual contributions of wall composition, hydration, and tip-sample interactions.

In contrast, DEX-treated primordia showed higher apparent Young’s modulus values across the epidermis (Fig. 5D, right panels). AFM heat maps revealed a broad increase in indentation stiffness across the scanned regions compared with MOCK-treated primordia. Quantification across 4-6 independent leaf primordia confirmed this difference, with the mean approach modulus increasing from ∼6-7 MPa under MOCK conditions to ∼10-11 MPa following DEX treatment, corresponding to an increase of ∼50-60% (Fig. 5H). Thus, epidermal TCP4 induction is associated with a substantial increase in the apparent indentation stiffness of the epidermal surface.

Retract measurements showed a similar response, with mean apparent Young’s modulus increasing from ∼8-9 MPa in MOCK-treated primordia to ∼12-13 MPa following DEX-treatment, corresponding to an increase of ∼44-55% (Fig. 5H). The difference between retract and approach moduli remained approximately 2 MPa under both conditions, indicating that the increase following TCP4 induction was detected consistently during both loading and unloading phases. Thus, two complementary AFM measurements independently revealed an increase in the apparent mechanical resistance of the epidermal surface following TCP4 induction.

Because AFM indentation primarily probes the mechanical response normal to the epidermal surface, we next assessed wall deformation in the plane of the tissue by measuring changes in epidermal cell area following removal or turgor. *PDF1;GR* seedlings at 2 DAI were imaged under fully turgid conditions in water and after plasmolysis in 0.6 M sorbitol for 30 minutes. The relative change in cell area between these conditions provide a complementary measure of the deformation sustained by the cell wall under turgor.

Under MOCK conditions, epidermal cells showed a broad distribution of area changes following plasmolysis, with a median relative change of ∼8% (Fig. 5E, G). In contrast, DEX-treated cells showed markedly less change in area, with a median close to 0% and a narrower distribution (Fig. 5F, G). Because the change in cell area following plasmolysis reflects the deformation sustained by the wall under turgor, the reduced response following DEX treatment indicates altered mechanical behaviour of the epidermal cells. This reduced deformation is consistent with decreased wall compliance, although differences in turgor pressure could also contribute to the observed response. Together with the AFM measurements, these results support a substantial alteration of epidermal mechanical properties following TCP4 induction.

Together, the increased apparent indentation stiffness, reduced turgor-associated cell deformation, and enhanced transverse CMT co-alignment indicate that epidermal TCP4 induction is accompanied by substantial changes in both the mechanical properties and cytoskeletal organisation of the epidermis. These changes are consistent with an epidermal state that favours proximodistal over mediolateral expansion, providing a potential mechanical basis for the persistent directional growth and narrow, axisymmetric geometry observed by live imaging experiments (Fig. 4, S7). Thus, strong epidermal TCP4 activity is associated with coordinated changes in growth directionality and wall mechanical behaviour that may contribute to the failure of blade flattening.

### 10. Epidermal TCP4 sustains auxin response during leaf development

The cross-layer effects of epidermal TCP4 activity raise the question of what possible downstream signals contribute to coordinating growth between tissue layers. Auxin represents a potential candidate, as TCP transcription factors have been implicated in the regulation of auxin biosynthesis, transport, and response (Koyama et al., 2010; Nicolas & Cubas, 2016; Challa et al., 2019; Koyama et al., 2025) Auxin signalling can also influence processes relevant to the phenotype observed here, including cytoskeletal organisation and pectin remodelling (Fu et al., 2005; Braybrook & Peaucelle, 2013; Ambrose & Wasteneys, 2014). We therefore investigated whether epidermal TCP4 induction is associated with altered auxin response during leaf development.

To examine auxin response following epidermal TCP4 induction, we crossed *PDF1;GR*^#1^ with a dual-marker line carrying the plasma membrane marker *pUBQ10::Lti6b-TdTomato* and the auxin-response reporter *pDR5VII::nls-3XmVENUS* (Le Gloanec et al., 2024). The membrane marker was used to outline cell boundaries, while *pDR5VII::nls-3XmVENUS* enabled visualisation of auxin response patterns during leaf development. We then performed time- lapse confocal imaging of first-leaf primordia from 1 to 5 DAI under continuous MOCK or DEX treatment.

In MOCK-treated primordia, *pDR5VII::nls-3XmVENUS* activity was readily detected at 1-2 DAI (Fig. S10D, MOCK). As development progressed, reporter activity gradually decreased and became more spatially restricted, with only weak signal remaining by 4-5 DAI (Fig. S10D). Thus, under MOCK conditions, the auxin-response reporter displayed a developmentally dynamic pattern, with strong activity during early primordium development followed by a progressive decline at later stages.

In contrast, DEX-treated primordia maintained *pDR5VII::nls-3XmVenus* activity throughout the developmental time course (Fig. S10D, DEX). At 1-2 DAI, reporter activity was comparable to or slightly stronger than under MOCK conditions. From 3-4 DAI onwards, however, the patterns diverged: while reporter activity progressively declined in MOCK-treated primordia, it remained readily detectable throughout DEX-treated leaves and persisted through 5 DAI (Fig. S10D). Moreover, reporter activity in DEX-treated primordia was broadly distributed across the organ, including inner tissue layers, rather than becoming spatially restricted as observed under MOCK conditions. Thus, epidermal TCP4 induction is associated with prolonged and spatially expanded auxin response during leaf development.

Because TCP4 was continuously induced throughout the reporter experiment, these data establish that epidermal TCP4 activity is associated with a sustained auxin response but do not resolve the timing or immediate dynamics of this response following TCP4 induction.

Collectively, these findings identify sustained auxin response as an additional component of the developmental changes associated with strong epidermal TCP4 activity. Whereas auxin reporter activity progressively declined during normal leaf development, epidermal TCP4 induction prolonged and spatially expanded this response, including into inner tissue layers. Together with the layer-specific changes in pectin organisation, increased mechanical resistance, and enhanced CMT co-alignment described above, these results reveal coordinated molecular, cell-wall, cytoskeletal, and mechanical changes accompanying TCP4-nduced growth restriction. Although the causal relationships between these responses remain to be resolved, their coordinated alteration provides a framework through which epidermal TCP4 activity could propagate across tissue layers to reorganise growth and ultimately reshape the developing leaf primordia.

Together, these findings reveal that epidermal TCP4 activity is associated with coordinated changes in auxin response, cell-wall reorganisation, cytoskeletal alignment, and tissue mechanical properties across the developing primordium. Their convergence provides a framework through which an epidermally initiated developmental signal can be translated into cross-layer changes in growth organisation, and ultimately, organ geometry.

## Discussion

Our results identify the spatial distribution of TCP4 activity across the developing leaf as an important component of its control over the timing and coordination of differentiation. TCP4 displays an asymmetric distribution, enriched in the adaxial epidermis, mesophyll and extending toward the abaxial epidermis, while altering its spatial domain of activity produces markedly different developmental outcomes. Induction from the endogenous TCP4 domain (*jaw-D;GR*) reduces leaf area and cell number while preserving organ development, whereas strong epidermal induction (*PDF1;GR*) causes rapid proliferative arrest and severely disrupt the lamina outgrowth. These observations suggest that the native spatial distribution of TCP may constrain the timing and magnitude of its activity across tissue layers. One possibility is that mesophyll-enriched TCP4 acts as a spatially buffered source or a rheostat from which activity progressively reaches neighbouring tissues, whereas direct strong expression in the epidermis bypasses this spatial regulation and prematurely engages differentiation-associated responses throughout the developing primordium.

The relationship between TCP4 mobility and its non-cell-autonomous activity provides an additional layer of spatial regulation. Mobile TCP4-mVENUS was detected beyond the epidermal expression domain, extending across two to three cell layers, yet mobility-restricted TCP4-3xGFP produced comparable cellular responses in both epidermal and subepidermal tissues. Thus, TCP4 protein movement is not required for cross-layer growth responses when TCP4 activity is strongly induced in the epidermis. This raises a paradox: if local epidermal TCP4 activity is sufficient, what function might protein mobility serve under native conditions? We propose that mobility and local activity fulfil complementary roles. In the endogenous expression domain, progressive movement of TCP4 between tissue layers could contribute to the spatial and temporal distribution of TCP4 activity, whereas once sufficient activity is established within a given layer, downstream signals may transmit their effects to neighbouring tissues without requiring further TCP movement. In this model, protein mobility contributes to the distribution of TCP4 activity, while non-cell-autonomous downstream signalling extends its developmental consequences beyond the cells containing TCP4 itself. The strong epidermal induction experiments. therefore, establish the sufficiency of TCP4 activity, but do not determine the contribution of protein mobility at endogenous expression levels. Distinguishing these possibilities will require quantitative comparison of TCP4 abundance and mobility across tissue layers under native and induced conditions, together with manipulation of TCP4 mobility without altering its expression levels or spatial domain.

The pronounced response to epidermal TCP4 activity also raises the possibility that tissue context influences how TCP4-dependent signals are translated into organ growth. The epidermis occupies a distinctive position in organ morphogenesis: it participates in auxin transport and signalling, and, as the outer mechanically continuous tissue layer, contributes strongly to the coordination of growth across the organ (Bhalerao & Bennett, 2003; Adamowski & Friml, 2015; Hervieux et al., 2017; Sapala et al., 2018). In our experiments, strong epidermal TCP4 induction was associated with sustained auxin response, increased epidermal mechanical resistance, and coordinated changes in growth across both epidermal and subepidermal layers. These observations suggest that TCP4 activity initiated in the epidermis may be particularly effective at coupling biomechanical and mechanical responses across the developing primordium. However, whether these cross-layer effects are transmitted predominantly through biochemical signals, mechanical coupling between tissues, or interaction between the two remains unresolved. The mechanosensitivity of auxin transport provides one potential point of interaction between these processes (Nakayama et al., 2012). Experimentally uncoupling auxin signalling from changes in wall mechanics will therefore be important for determining how these pathways contribute to non-cell-autonomous effects of epidermal TCP4.

Auxin provides one plausible link between TCP4 activity and the cytoskeletal and cell-wall changes observed here. Auxin can influence both cortical microtubule organisation through ROP-dependent pathways and pectin remodelling through regulation of cell-wall modifying processes (Fu et al., 2005; Peaucelle et al., 2011; Braybrook & Peaucelle, 2013). In our system, sustained auxin response coincided with enhanced CMT co-alignment, layer-specific changes in pectin epitopes, and increased epidermal mechanical resistance. We therefore propose a working model in which these responses form an interconnected network downstream of strong TCP4 activity, with auxin signalling, cytoskeletal organisation and wall mechanics potentially reinforcing one another during the transition from proliferation to differentiation. Such coupling could provide a mechanism by which an initial epidermal perturbation becomes stabilised across the developing primordium and persists after TCP4 induction is withdrawn. The causal ordering of these events remained to be determined; time-resolved analysis following acute TCP4 induction, combined with independent perturbation of auxin signalling, CMT organisation and pectin remodelling, should distinguish whether these responses occur sequentially or in parallel.

The mechanical changes observed following epidermal TCP4 induction are also consistent with models in which tissue stiffness and initial geometry jointly determine leaf flattening. Finite-element modelling has shown that increasing epidermal stiffness can restrict the buckling required for lamina formation, whereas greater compliance favours symmetry breaking and blade expansion (Zhao et al., 2020). In our experiments, TCP4 induction increased epidermal apparent Young’s modulus from ∼6-7 to ∼10-13 MPa and was accompanied by the persistence of a narrow, axisymmetric primordium. Initial geometry is likely to be equally important: the cylindrical phenotype of *wox1;prs;as2* illustrates how reduced primordial asymmetry can compromise subsequent flattening even in the absence of the TCP4 perturbation observed here. Together, these observations lead us to propose that the spatial and temporal regulation of TCP4 contributes to a transient window of epidermal compliance during early leaf development. Low TCP4 activity in the epidermis during early primordium growth may permit the mechanical deformation and symmetry breaking required for blade outgrowth, whereas subsequent TCP4 activity could contribute to the progressive restriction of growth as differentiation proceeds. Premature strong epidermal TCP4 activity would therefore close this compliance window too early, stabilising an axisymmetric growth state before lamina expansion is established.

More broadly, the diversification of TCP regulatory networks across plant lineages raises the possibility that changes in the spatial and temporal deployment of TCP activity have contributed to the evolution of leaf form (Ori et al., 2007; Nicolas & Cubas, 2016). By coupling the timing of proliferative exit to cytoskeletal organisation, cell-wall mechanics and tissue growth, TCP activity could tune the duration of the morphogenetic window during which the developing lamina remains permissive to symmetry breaking and blade expansion. Evolutionary changes in the timing, level or tissue distribution of TCP activity could therefore shift this developmental window, providing a mechanism through which a conserved growth-regulatory network generates diversity in leaf architecture.

## Materials and Methods

### Growth conditions and plant material

All transgenic lines used in this study were generated in the *jaw-D* background; therefore, *jaw-D* was used as the primary control for phenotypic comparisons. In selected early experiments, *Arabidopsis thaliana* ecotype Columbia-0 (Col-0) was included as a wild-type reference. A complete list of plant lines used and generated in this study is provided in Supplementary Table S1.

Seeds were surface sterilised in 70% ethanol containing 0.05% SDS for 10 min, followed by two washes with 100% ethanol (Shankar et al., 2023). Sterilised seeds were sown on Murashige and Skoog (MS) medium (PT011, HiMedia Laboratories, India) supplemented with either 0.025% ethanol (MOCK) or 12 μM dexamethasone (DEX; D4902, Sigma-Aldrich, USA). After sowing, plates were stratified for 3 days in the dark at 4 °C to synchronise germination.

Following stratification, plates were transferred to a plant growth chamber (I-41LL, Percival Scientific, USA) maintained at 22 °C under a 16 h light/8 h dark photoperiod with a light intensity of approximately 100 μmol m⁻² s⁻¹. Twelve-day-old seedlings were transplanted to soil composed of Soilrite and perlite (Keltech Energies, India) in a 3:1 ratio and grown to maturity under the same environmental conditions.

### DNA constructs and generation of transgenic lines

To generate the epidermis-specific inducible construct *pPDF1::rTCP4-GR*, a 1.5 kb fragment of the PDF1 promoter (*AT2G42840*) was amplified from genomic DNA and cloned into a TA cloning vector using the InsTAclone PCR Cloning Kit (Thermo Scientific, #K1213). The promoter fragment was subsequently subcloned into the binary vector pCAMBIA1390 using *HindIII* and *SalI* restriction sites. The *rTCP4-GR* fragment was excised from clone PUTN97 using *SalI* and *NcoI* and inserted downstream of the PDF1 promoter to generate the final construct *pPDF1::rTCP4-GR*.

For the subepidermal construct *pAN3::rTCP4-GR*, a 1.9 kb fragment of the AN3 promoter was excised from clone PUTN210 using *EcoRI* and inserted into pCAMBIA1390. The *rTCP4-GR* fragment was subsequently cloned downstream using *NcoI* and *HpaI* restriction sites.

For fluorescent localisation studies, the *mVENUS* construct was amplified by PCR and cloned into pCAMBIA1390 using *BamHI* and *NcoI*. The rTCP4 coding sequence was excised from clone PUTN97 using *SalI* and *BamHI,* and inserted upstream of *mVENUS,* together with an NLS sequence, to generate *rTCP4-mVENUS*. This fragment was then transferred into the pCAMBIA vector containing the PDF1 promoter using *SalI* and *NcoI*, yielding the final construct *pPDF1::rTCP4-mVENUS*.

To generate the mobility-restricted construct *pPDF1::rTCP4-3×GFP*, the *SV40NLS-3×GFP* fragment was excised from plasmid 129 (*pAtML1::3×GFP*, a gift from Gerd Jürgens and Ulrike Hiller) using *BamHI* and *SacI* and cloned into the binary vector pCAMBIA1300. The fragment was subsequently subcloned into pCAMBIA1390 using *EcoRI* and *BamHI*. The *pPDF1::rTCP4* fragment was excised from pCAMBIA1390;*pPDF1::rTCp4* using *ApaI* and *BamHI* and cloned upstream of *SV40NLS-3×GFP* to obtain the final construct *pPDF1::rTCP4- 3×GFP*.

Details of all constructs generated in this study and primer sequences used for promoter amplification and genotyping are provided in Supplementary Table S2.

### Plant transformation and selection of transgenic lines

All binary constructs were introduced into the *jaw-D* background using the *Agrobacterium tumefaciens*-mediated floral dip method. For transformation, Silwet L-77 was added to the infiltration solution at a concentration of 50 μl L⁻¹.

T₁ seeds were selected on MS medium supplemented with 20 mg L⁻¹ hygromycin. For each construct, at least ten independent transgenic lines were obtained and advanced to the T₃ generation to establish homozygous lines. Genotyping and phenotypic analyses were used to confirm transgene integration and expression.

From the homozygous T₃ populations, representative *pPDF1::rTCP4-GR* and *pAN3::rTCP4- GR* lines displaying a range of phenotypic severities were selected for detailed analysis. For *pPDF1::rTCP4-mVENUS* and *pPDF1::rTCP4-3×GFP*, viable lines were taken forward for analysis after homozygosity was confirmed.

### Histochemical GUS staining and three-dimensional imaging

To examine the spatial distribution of TCP4 and MIR319C expression during early leaf development, previously described GUS reporter lines were used: *pTCP4::GUS* (Shankar et al., 2023) and *pTCP4::TCP4-GUS* (Challa et al., 2019) to monitor TCP4 promoter activity and TCP4 protein accumulation, respectively, and *pMIR319C::GUS* (Shankar et al., 2023) to assess MIR319C promoter activity. All reporter lines were in the Col-0 background.

Seedlings were grown under standard growth conditions, and the first pair of true leaves were exposed by excising one cotyledon with forceps at 2-3 days after initiation (DAI). Tissues were fixed in ice-cold 90% acetone for 20 minutes at room temperature and then washed with ice-cold staining buffer containing 50 mM sodium phosphate (pH 7.0), 0.2% Triton X-100, 5 mM potassium ferrocyanide, and 5 mM potassium ferricyanide. For the GUS reaction, samples were incubated in staining solution containing 50 mM sodium phosphate buffer (pH 7.0), 0.5 mg mL⁻¹ 5-bromo-4-chloro-3-indolyl-β-D-glucuronide (X-Gluc), 5 mM potassium ferricyanide, 5 mM potassium ferrocyanide, 0.1% Triton X-100, and 10 mM EDTA. Samples were vacuum infiltrated for 10-15 min and incubated at 37 °C for 3 h. The reaction was terminated by replacing the staining solution with 70% ethanol, followed by sequential ethanol washes to remove chlorophyll and rinsing with water.

GUS-stained samples were subsequently processed using the modified pseudo-Schiff propidium iodide (mPS-PI) method as previously described (Truernit et al., 2008). Briefly, tissues were oxidised with periodic acid, followed by incubation in Schiff’s reagent containing 100 mM sodium metabisulfite and 0.15 M HCl, to which propidium iodide was freshly added to a final concentration of 100 μg/mL. Samples were mounted in chloral hydrate solution containing 4 g chloral hydrate, 1 mL glycerol, and 2 mL water

Samples were imaged using Zeiss LSM 980 Airyscan2 upright confocal microscope (MechanoDevo team, RDP, ENS de Lyon) with a 40× oil-immersion objective (NA 1.4). Propidium iodide fluorescence was excited at 561 nm and detected between 580 and 650 nm using a GaAsP detector (laser intensity, 2-10%). GUS precipitate was visualised by confocal reflection microscopy using 488 nm illumination and a T80/20 reflection filter, with signal collected between 480 and 490 nm using a GaAsP detector (laser intensity, 5-30%).

Optical stacks encompassing the entire leaf primordium were acquired at 512×512 pixels and 16-bit depth with 1 µm z-step. Image stacks were imported into MorphoGraphX (Barbier de Reuille et al., 2015) for three-dimensional reconstruction. Virtual sections were generated using clipping planes to obtain mid-sagittal sections along the midrib axis and transverse sections across distal, middle, and proximal regions of the lamina.

Control samples lacking GUS reporter constructs were processed in parallel using identical staining and imaging conditions to assess background signal.

### Measurement of leaf area, epidermal and subepidermal cell size, and cell number

The first pair of true leaves was collected from 29-day-old plants. Because several genotypes exhibited curved or crinkled leaf morphology, leaves were gently flattened before imaging. Digital images were acquired with a Google Pixel 7a camera (*f/*1.9, 64 MP rear sensor, 1/25 shutter speed, 5.43 mm focal length, and ISO 222), leaf outlines were isolated in Windows 3D Paint, and leaf area was measured in ImageJ (National Institutes of Health; https://imagej.nih.gov/ij/).

For cellular analyses, leaves were first fixed in 70% ethanol and subsequently transferred to ethanol:glacial acetic acid (7:1) for 12-14 h at room temperature. Tissues were incubated in 1 M KOH for up to 30 min with gentle rotation, washed twice with double-distilled water and treated with lactic acid until fully transparent (Challa et al., 2019).

Cleared leaves were mounted and imaged by differential interference contrast (DIC) microscopy (Olympus, USA) using a 20× objective. Images were acquired from the abaxial side to visualise both epidermal pavement cells and the underlying subepidermal bundle sheath mesophyll cells. Multiple fields of view were captured from distal, medial, and proximal regions of each leaf.

For epidermal cell size analysis, a total of approximately 200-600 pavement cells were measured across 7-12 independent leaves per genotype or treatment using ImageJ. Subepidermal cell size was measured from approximately 200-800 bundle sheath mesophyll cells across 8-12 leaves. Mean cell size was calculated separately for each tissue layer.

The total number of cells per leaf in each layer was estimated by dividing measured leaf area by the corresponding mean cell area.

### Temporal TCP4 induction and transfer assays

To assess temporal effects of TCP4 induction on leaf growth, *jaw-D;GR* and *PDF1;GR^#1^* seedlings were subjected to reciprocal MOCK and DEX transfer treatments. Seedlings were initially grown on MOCK- or DEX-containing medium and, at 4 days after stratification (DAS), were transferred to fresh medium containing either the same or reciprocal treatment.

Four regimes were established: continuous MOCK (MtM; MOCK-to-MOCK), continuous DEX (DtD; DEX-to-DEX), MOCK-to-DEX transfer at 4 DAS (MtD), and DEX-to-MOCK transfer at 4 DAS (DtM). Seedlings were subsequently grown under identical environmental conditions until 29 days. Projected leaf area was quantified from calibrated images and expressed in mm². Epidermal and subepidermal cellular parameters were analysed by DIC microscopy as described above. Relative changes in leaf area, cell number, and cell area were calculated with respect to the continuous MOCK control (MtM).

### Confocal imaging and analysis of protein mobility

Fluorescent reporter lines were used as controls to establish tissue-specific localisation patterns. Wild-type Col-0 seedlings served as a negative control. *p35S::GFP* was used as a control for broadly distributed fluorescence, whereas *pPIN1::PIN1-GFP* and *pPIN3::PIN3- GFP* (Le Gloanec et al., 2024) provided membrane-localised controls with characteristic tissue distributions. The *pPDF1::MBD-mCitrine* line (Trinh et al., 2024) served as a control for epidermis-restricted localisation. *pPDF1::rTCP4-mVENUS* and *pPDF1::rTCP4-3×GFP* seedlings were analysed as test samples.

The first pair of true leaves was exposed by excising one cotyledon at 2-3 or 4-5 DAI, representing early and later developmental stages, respectively. Seedlings were stained with 1 mg.mL^-1^ propidium iodide in water for 2 minutes, immediately rinsed, placed in 60-mm Petri dishes containing ½ MS medium, and immersed with water for imaging.

Samples were imaged using either a Zeiss LSM 800 confocal microscope (Kierzkowski lab, IRBV, Université de Montréal) or a Zeiss LSM 980 Airyscan2 microscope (MechanoDevo team, RDP, ENS de Lyon), using long-working-distance water-immersion objectives. Propidium iodide fluorescence was excited at 561 and detected between 580 and 650 nm using a GaAsP detector (laser intensity, 2-10%). mVENUS and GFP were excited at 488 nm and detected between 490-550 nm (laser intensity, 5-8%).

Optical stacks encompassing the entire leaf primordium were acquired at 512×512 pixels and 16-bit depth with a 1 µm z-step. Stacks were imported into MorphoGraphX for three- dimensional reconstruction. Virtual sagittal and transverse sections were generated using clipping planes.

For quantitative analysis, a sagittal clipping plane containing the PI and reporter channels was extracted from the reconstructed primordium. The PI channel was filtered three times using a Gaussian Blur of 0.3, and the surface was detected with the “Edge detect” tool with thresholds between 6,000 and 13,000, adjusted according to PI signal intensity. An initial 5 µm cube-size mesh was generated and subdivided three times. The PI membrane signal was projected over a depth of 1-3 µm, and the mesh was segmented using auto-seeding followed by manual correction.

Reporter fluorescence was projected onto the cellular mesh using Mesh > Signal > Project Signal function, with MinDist = 1 μm and MaxDist = 3 μm. Fluorescence per cell was quantified using Mesh > Heat Map > Measures > Signal with Signal Total as the quantification metric. Heat maps therefore represent total reporter fluorescence associated with individual segmented cells.

### Scanning electron microscopy

Scanning electron microscopy (Maeda et al., 2014) was performed to examine early leaf primordia. Tissues were collected in fixation buffer containing 0.1 M sodium phosphate buffer (pH 7.0) and 3% glutaraldehyde and vacuum infiltrated in a desiccator for 30 min at 400-500 mmHg. The fixation buffer was replaced with fresh solution, and samples were incubated for 12-24 h at 4 °C.

Samples were subsequently transferred to post-fixation buffer containing 1% osmium tetroxide in 0.1 M sodium phosphate buffer (pH 7.0) and incubated for 36-48 hours at 4 °C. Tissues were rinsed once with 1×PBS and dehydrated through a graded ethanol series (30%, 40%, 50%, 60%, 70%, 80%, 90%, and 100%) with each step performed for 30 min on ice. Samples were then subjected to critical point drying.

Dried samples were sputter-coated with gold for 38-60 s under vacuum and imaged using a JEOL SEM-300 scanning electron microscope (AFMM facility, IISc) to visualise the shoot apical meristem and developing leaf primordia.

### Preparation of seedlings for live confocal imaging

For live-imaging experiments, seeds were stratified and germinated on ½ MS medium supplemented with 0.7% agar, 1% sucrose, and 0.1% Plant Preservative Mixture (PPM). One cotyledon was carefully removed at the experiment-specific developmental stage to expose the developing leaf primordia. Seedlings were transferred to Ø60 mm plates containing ½ MS medium supplemented with 1.5 % agar.

During imaging, plates were positioned horizontally and seedlings were immersed in sterile water supplemented with 0.1% PPM. Between repeated imaging sessions, seedlings were returned to a vertical orientation and maintained under long-day conditions (16 h light / 8 h dark) at 22 °C.

Unless otherwise specified, live imaging was performed using a Zeiss LSM980 Airyscan2 upright confocal microscope equipped with a 25× long-working-distance water-dipping objective (NA 1.2).

### Time-lapse imaging of epidermal growth

To quantify epidermal growth dynamics, *pUBQ10::myr-YFP* (Willis et al., 2016) was crossed into the *PDF1;GR* background. Seedlings were prepared as described above, with cotyledon removal performed at 2 DAI.

The first pair of true leaves was imaged from 2 to 5 DAI at 24-hour intervals until under continuous MOCK or DEX treatment. At each time point, at least half of the abaxial epidermal surface was imaged.

YFP was excited at 514 nm and detected between 520 and 560 nm. Image stacks were acquired at 512×512 pixels and 16-bit depth with a 1 µm z-step. When the primordium exceeded the field of view, overlapping stacks were acquired and subsequently stitched in MorphoGraphX.

### Cell lineage tracking and growth quantification

Confocal image stacks were analysed in MorphoGraphX. The *pUBQ10::myr-YFP* plasma membrane signal was used to reconstruct the surface geometry of the leaf primordium and segment individual epidermal cells.

Surface extraction was performed using the “Edge Detect” function, with thresholds adjusted between 6,000 and 13,000 according to signal intensity, followed by refinement using the “Edge Detect Angle” thresholds of 3,000-6,500. An initial 5 µm cube-size surface mesh was generated, subdivided three times and smoothed. The membrane signal was projected onto the mesh over a depth of 2-6 µm.

Cell segmentation and lineage tracking were performed manually. Parent-daughter relationships between successive time points were verified using the MorphoGraphX “Check Correspondence” function.

Area expansion was calculated from changes in cellular surface area between consecutive time points. For cells undergoing division, the area of the mother cell at time *t* was compared with the combined area of its daughter cells at time *t + 1*, providing a measure of local growth independent of cell division (Barbier de Reuille et al., 2015).

Cell division were identified by the appearance of two daughter cells derived from a single parent cell between successive imaging frames, and their spatial distribution was mapped across the primordium.

Growth anisotropy was calculated from the principal components of the local deformations between successive time points using MorphoGraphX growth-analysis tools. Principal directions of growth (PDGs) were calculated from the local deformation tensor, with the maximal principal growth direction visualized as vectors across the primordium.

Area expansion, cell division, growth anisotropy, and PDG distributions were visualized on the reconstructed mesh.

### Auxin-response reporter imaging

To monitor auxin-response dynamics during early leaf development, *PDF1;GR* plants were crossed with the plasma membrane reporter *pUBQ10::myr-tdTomato* and the nuclear auxin- response reporter *pDR5VII::nls-3xmVENUS* (Le Gloanec et al., 2024).

Seedlings were prepared and imaged under continuous MOCK or DEX treatment using the live-imaging procedure described above. tdTomato and mVENUS were excited at 561 and 514 nm, respectively. Emission was collected between 580 and 610 nm for tdTomato and between 520 and 550 nm for mVENUS. Image stacks were acquired at 512 × 512 pixels and 16-bit depth with a 1 μm z-step.

### Cortical microtubule imaging and orientation quantification

To visualise cortical microtubule organisation in the epidermis, seedlings expressing *pPDF1::MBD-mCitrine* (Trinh et al., 2023) were analysed. For TCP4-induction experiments, the reporter was crossed into the *pPDF1::rTCP4-GR* background.

Seedlings were prepared using the live-imaging protocol described above, except that cotyledon removal and imaging were performed at 3 DAI. High-resolution imaging was performed using the Zeiss LSM980 Airyscan detector in Airyscan Multiplex SR-4Y mode.

mCitrine was excited at 488 nm and detected between 518-550 nm. Z-stack were acquired using the 25× water-immersion objective (NA 1.2) with a 0.3–0.5 μm z-step, 2.3× zoom, a 1024 × 1024-pixel frame, and 1× line averaging. Laser power, detector gain, and acquisition settings were kept identical between treatments.

Raw Airyscan data were reconstructed in Zeiss ZEN software. Reconstructed images were imported into MorphoGraphX for image assembly. When multiple overlapping fields were required, stacks were aligned and merged before export as high-resolution PNG files.

CMT orientation was quantified using FibrilTool in batch mode (Boudaoud et al., 2014) in ImageJ/FIJI. Individual epidermal cells were manually delineated as regions of interest (ROIs). FibrilTool was used to calculate the mean CMT orientation and anisotropy within each ROI, with anisotropy ranging from 0 (isotropic organisation) to 1 (perfect alignment).

Orientation data were used to generate rose plots with custom Python scripts. Directional organization was evaluated using circular statistics, including the Rayleigh test, mean resultant length (*r*), and circular standard deviation.

### Atomic force microscopy (AFM) measurements

Atomic force microscopy (AFM) experiments (Beauzamy et al., 2015) were performed using a stand-alone JPK Nanowizard III microscope equipped with a CellHesion module and controlled by JPK Nanowizard software version 6.1. The CellHesion module provides an extended 100 µm Z-piezo range on the microscope stage, increasing the available vertical displacement several-fold compared with the head Z-piezo (∼15 µm). Measurements were performed in Quantitative Imaging (QI) mode, which records a force-distance curve at each pixel.

Experiments were conducted in distilled water at room temperature. A silica spherical probe (Nanotools Biosphere B300 NCH, NanoAndMore) mounted on a silicon cantilever with a nominal spring constant of 26 N m⁻¹ (pre-calibrated by the manufacturer) was used for all measurements. The probe had a radius of 300 ± 10 nm. AFM scans were acquired over a 50 µm × 50 µm area with a 500 nm pixel size. The force trigger was set to 500 nN, corresponding to an indentation depth of approximately 150-200 nm in MOCK samples. Force curves were recorded with a 2 µm ramp size and approach and retract speeds of 50 µm s⁻¹.

The deflection sensitivity of the cantilever was determined in contact mode by performing a linear fit of the contact region of a force curve acquired on a sapphire reference surface in PBS. The same cantilever was used for all measurements included in this study to ensure experimental consistency.

To minimise force offsets introduced during repeated calibrations, particularly during deflection-sensitivity measurements, the photodiode sum signal was used as a proxy for the laser’s position on the cantilever. The laser was first aligned near the end of the cantilever using the instrument’s top-view camera. The photodiode sum signal was then maximised by adjusting the laser position perpendicular to the cantilever’s longitudinal axis, ensuring the laser spot was close to the cantilever axis. The resulting sum signal value was recorded and used as a reference for subsequent experiments, enabling reproducible laser alignment during remounting of the cantilever.

For the experiment, Seeds were stratified and germinated on ½ Murashige and Skoog (MS) medium supplemented with 0.7% agar, 1% sucrose, and 0.1% Plant Preservative Mixture (PPM). After stratification and germination, a cotyledon from a 4 DAI seedling was dissected meticulously to expose the leaf primordium. For *PDF1;GR* MOCK, leaves were completely excised, and for DEX, the seedling with leaves and the cotyledon as a support was retained. For AFM, seedlings were mounted horizontally in Ø60 mm Petri dishes containing the polymer resin, and the resin was allowed to polymerise to stabilise tissue movements. The leaves were in such a position that the cantilever reached the abaxial surface.

### AFM data analysis

Force-distance curves were analysed using JPK Data Processing software version 8.1. Raw force vs height curves were first flattened by subtracting a linear fit calculated from the non- contact region of the approach curve, thereby setting the baseline force to zero.

An initial estimate of the point of contact (POC) was obtained by identifying the first point at which the force crossed zero as the approach curve was traced from the setpoint force position. The tip-sample distance was then calculated using the relation:

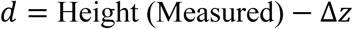

where Δ*z* represents the cantilever deflection.

The apparent Young’s modulus was determined by fitting the entire force-indentation curve using the Hertz contact model implemented in JPK Data Processing. The model is derived from the axisymmetric indentation solution described by Sneddon (Sneddon, 1965). The relationship between indentation depth *δ* and applied force *F* is given by:

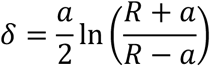

where *R* is the tip radius, *a*is the radius of the contact area, and *E*^∗^ is the reduced Young’s modulus, defined as:

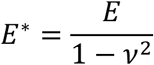

*E* representing the Young’s modulus of the sample and *ν* the Poisson’s ratio. Expanding the solution for small indentation relative to the tip radius (*δ*/*R*) yields:

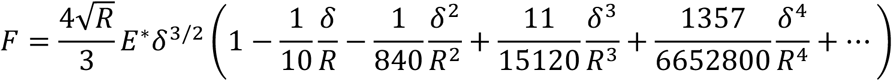

In many AFM studies, the Hertz model corresponds to the first term of this expansion, whereas JPK Data Processing uses the full spherical solution, implemented as the paraboloid model. Differences between these models become significant only when the indentation depth approaches the tip radius, which lies near the limits of their applicability.

For the fitting procedure, the tip radius was set to 300 nm, and Poisson’s ratio was fixed at 0.5, a commonly used value for biological materials. The Young’s modulus, point of contact, and force offset were treated as free parameters during the fitting procedure. The same analysis protocol was applied to both approach and retract curves.

AFM-derived mechanical parameters were visualised using heat maps and quantitative graphs to compare tissue stiffness across genotypes and treatments. Elastic modulus values calculated from individual force-indentation curves were mapped onto the corresponding AFM scan grid to generate spatial stiffness heat maps, allowing visualisation of local variations in mechanical properties across the leaf surface. Heat maps were generated from the Young’s modulus values obtained at each pixel of the Quantitative Imaging (QI) scan, and a representative image was used.

For statistical analysis, Young’s modulus values from multiple scans and biological replicates were compiled and plotted as distribution plots and summary graphs. Mean values and sample variability were represented using box plot graphs, with appropriate statistical descriptors. These visualisations enabled comparison of epidermal mechanical properties between MOCK and dexamethasone-treated samples, highlighting changes in tissue stiffness associated with TCP4-mediated differentiation.

All graphical representations were generated from processed AFM datasets using standard data analysis software following export from the JPK Data Processing platform.

### Osmotic treatment and quantification of cell deformation

To assess epidermal mechanical responses to osmotic perturbation (Ma et al., 2022), *pUBQ10::myr-YFP* was crossed into the *pPDF1::rTCP4-GR* background. Seedlings were prepared at 2 DAI using the live-imaging procedure described above.

Seedlings were first incubated in distilled water for 30 min. Tissues were stained with propidium iodide (1 mg mL⁻¹) for 3 min and immediately imaged using the Zeiss LSM980 upright confocal microscope with a 25× water-immersion objective. PI was excited at 561 nm and detected between 580 and 650 nm.

Following initial imaging, the same samples were exposed to 0.6 M sorbitol for 30 min and re- imaged using identical acquisition settings. Overlapping z-stacks were stitched when necessary.

Matched-cell analysis was performed in MorphoGraphX. Surface meshes were generated for images acquired in water and after sorbitol treatment, and individual epidermal cells were segmented and matched between conditions.

For each matched cell, the relative decrease in cell area was calculated as: 1- (A_Sorbitol/A_water) x 100, where A_water and A_sorbitol represent cell area before and after sorbitol treatment, respectively. Positive values indicate a decrease in cell area following osmotic treatment, whereas negative values indicate an apparent increase.

Cell-wise area changes were projected onto the sorbitol-treated mesh, and heat maps were displayed using identical scales across experimental conditions.

### Immunohistochemistry

Immunohistochemistry was performed to examine pectin composition and methylesterification using monoclonal antibodies recognising homogalacturonan epitopes. The procedure was adapted from Haas et al. (2020) with minor modifications.

6 DAI *pPDF1::rTCP4-GR* seedlings grown under MOCK or DEX conditions were fixed in FAA (formalin-acetic acid-alcohol). Fixed tissues were dehydrated through a graded ethanol series, cleared using Histoclear (Sigma-Aldrich), embedded in Paraplast, and sectioned at 4 μm using a Leica microtome. Sections were dewaxed before immunolabelling.

Sections were incubated overnight with LM19 or LM20 primary antibodies (Agrisera; 1:1000 dilution), recognising low-/un-methylesterified and highly methylesterified homogalacturonan epitopes, respectively. After washing, sections were incubated for 4 h with Alexa Fluor 647- conjugated secondary antibody (Thermo Fisher Scientific; 1:10,000 dilution). Sections were subsequently counterstained with Calcofluor White containing Evans blue and mounted using antifade mounting reagent (Thermo Fisher Scientific).

Fluorescence imaging was performed using a Zeiss LSM800 confocal laser-scanning microscope (Departmental Confocal Facility, MCB Department, IISc) with a 40× oil- immersion objective (NA 1.4). Alexa Fluor 647 was excited at 642 nm and detected between 660 and 760 nm. Calcofluor was excited at 405 nm and detected between 415 and 500 nm. Imaging parameters were kept constant between samples. Z-stacks were acquired with a 0.5 μm z-step, stitched in MorphoGraphX, and the antibody channel was overlaid on the Calcofluor channel.

Four independent leaves were analysed per genotype and antibody (n = 4).

### Statistical analysis

Graphical representations in Figs. 2, 3, S1, S3, S4, S5, S6, S8, S9 and S11 and the corresponding statistical analyses were performed using GraphPad Prism 9 (GraphPad, USA). Data were first tested for normality using the Shapiro-Wilk test. For normally distributed data, pairwise comparisons were performed using unpaired t-tests, with Welch’s correction when variances were unequal. Differences among multiple genotypes were assessed using ordinary one-way ANOVA followed by Šídák’s multiple-comparisons test.

Graphical representations in Figs. 4, 5 and S7 were generated using custom Python scripts. Statistical comparisons were performed using Student’s t-tests where indicated. Circular CMT orientation data were analysed using the Rayleigh test, mean resultant length (r), and circular standard deviation. Statistical significance is reported as exact p-values.

### Image assembly

Image assembly was done using Apple Keynote.

## Accession Numbers

The Arabidopsis genes analysed or used as markers in this study correspond to the following locus identifiers from TAIR (The Arabidopsis Information Resource, www.arabidopsis.org). The epidermal marker *PROTODERMAL FACTOR1 (PDF1/ATML1)* corresponds to *AT2G42840* and is commonly used to drive epidermis-specific expression. The transcriptional co-activator *ANGUSTIFOLIA3 (AN3/GIF1)*, corresponding to *AT5G28640*, is predominantly expressed in subepidermal tissues and regulates cell proliferation. Auxin transport markers *PIN-FORMED1 (PIN1)* and *PIN-FORMED3 (PIN3)* correspond to *AT1G73590* and *AT1G70940*, respectively. The TCP transcription factors examined in this study belong to the CIN-TCP/JAW-TCP clade, including *TCP2 (AT4G18390)*, *TCP3 (AT1G53230), TCP4 (AT3G15030), TCP10 (AT2G31070), and TCP24 (AT1G30210),* which are negatively regulated by miR319 and function in controlling the transition from cell proliferation to differentiation during leaf development. In addition, the cortical microtubule marker MBD used in reporter constructs (e.g., MBD-GFP) corresponds to the microtubule-binding domain of mammalian MAP4. It therefore does not represent an endogenous Arabidopsis locus.

## Acknowledgements

We acknowledge the contribution of SFR Biosciences (Université Claude Bernard Lyon 1, CNRS UMR3444, Inserm US8, ENS de Lyon) and the PLATIM-LyMIC facility for assistance with AFM, the Departmental Confocal Facility (supported by DAE-FIST) for imaging experiments, and the Advanced Facility for Microscopy and Microanalysis (AFMM) at IISc for SEM experiments. We thank Annalisa Bellandi, Denise Arico, Charlotte Kirchhelle, Christophe Trehin, Duy-Chi Trinh, Adrienne Roeder and Monalisha Rath for helpful discussions during the development of the project. We thank Marjolaine Martin for providing the reporter lines *pPDF1::MBD-mCitrine* and pUBQ*10::Lti6b-tdTomato*, and Claire Lionnet and the MechanoDevo team for microscopy training. We thank Gerd Jürgens and Ulrike Hiller for providing the *pAtML1::3×GFP* vector. Confocal imaging was conducted using an instrument supported by Canada Foundation for Innovation grants (37805) to D.K.

## Author contributions

A.S. initiated the project by generating multiple transgenic lines and preliminary results, assisted with organ and cellular morphometry experiments, and contributed to revisions. V.M. subsequently led the experimental work, designed and performed most experiments, analysed and interpreted the results, organised the figures, wrote the first draft of the manuscript, and contributed to its finalisation and revision. C.L.G. assisted V.M. with confocal image acquisition (protein fusions), MorphoGraphX analysis, and manuscript revision and editing.

A.G. assisted with the SEM and Immunohistochemistry experiments. G.V. assisted with organ and cellular morphometrics. S.B. assisted with performing and analysing the AFM experiments. D.K. contributed to imaging experiments, image-analysis infrastructure, editing, and proofreading. O.H. and U.N. contributed to experimental design and data interpretation, supervised the work, and corrected, revised, and finalised the manuscript.

## Source data

All the raw images and source data shown in this study are deposited in Biostudies database record and can be downloaded following this link: https://www.ebi.ac.uk/biostudies/studies/S-BSST3250

## SUPPLEMENTAL FIGURES AND TABLES

**Fig. S1.**
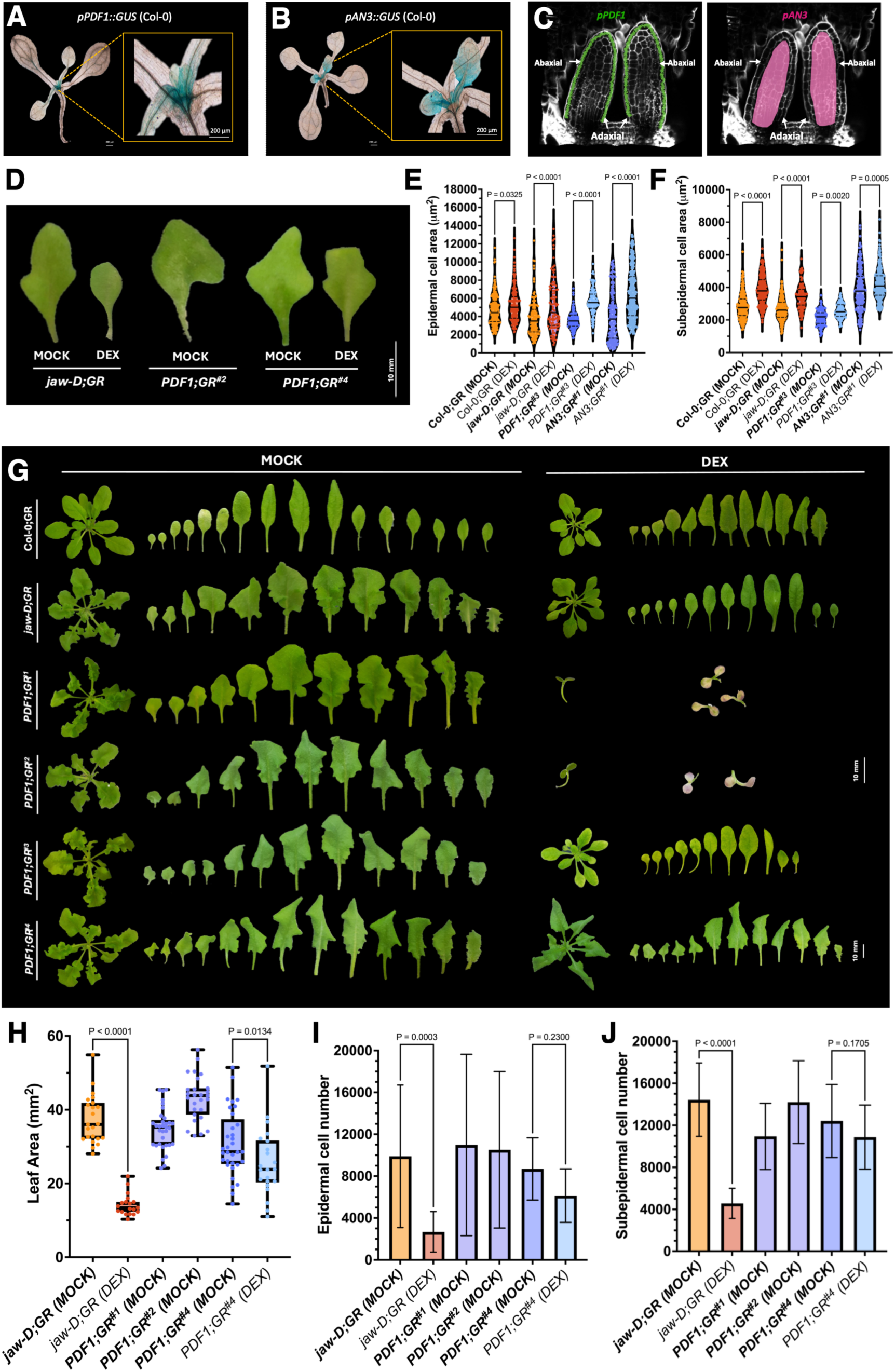
Promoter-specific TCP4 induction reveals layer-dependent control of cell proliferation and expansion in leaf size rescue. (A, B) Bright-field images of whole-mount leaves with corresponding magnified insets showing GUS reporter activity driven by the epidermis-specific promoter *pPDF1* (A) and the mesophyll-associated promoter *pAN3* (B) in 8-day-old seedlings. Scale bars, 200 µm. (C) Schematic representation of *pPDF1* (green) and *pAN3* (magenta) expression domains superimposed on an optical sagittal section of a developing leaf (grey), illustrating their relative tissue-layer localisation. (D) Representative images of the mature first leaf pair from 28-day-old plants of the indicated genotypes and chemical treatments. Scale bar, 10 mm. (E, F) Violin plots showing the distribution of epidermal (E) and subepidermal (F) cell areas for the indicated genotypes and treatments. N = 180-600 cells. Statistical significance was assessed by one-way ANOVA followed by a Šídák post hoc test (P-values indicated above the compared means). (G) Representative images of whole rosettes and individual rosette leaves from the indicated genotypes under MOCK and dexamethasone (DEX) treatments, corresponding to inducible TCP4 expression from the endogenous pTCP4 promoter or the epidermis-specific pPDF1 promoter. Scale bar, 10 mm. (H) Quantification of leaf area (Y-axis) for the mature first leaf pair of the indicated genotypes and treatments (X-axis). N = 20-34 leaves. Statistical analysis as in (E, F). Error bars represent SD. (I, J) Estimated epidermal (I) and subepidermal (J) cell numbers for the indicated genotypes and treatments. Cell number was calculated as the ratio of total leaf area to mean cell area (10-13 leaves per genotype). Statistical analysis as in (E, F); error bars represent SD. P-values are indicated above the compared means.

**Fig. S2.**
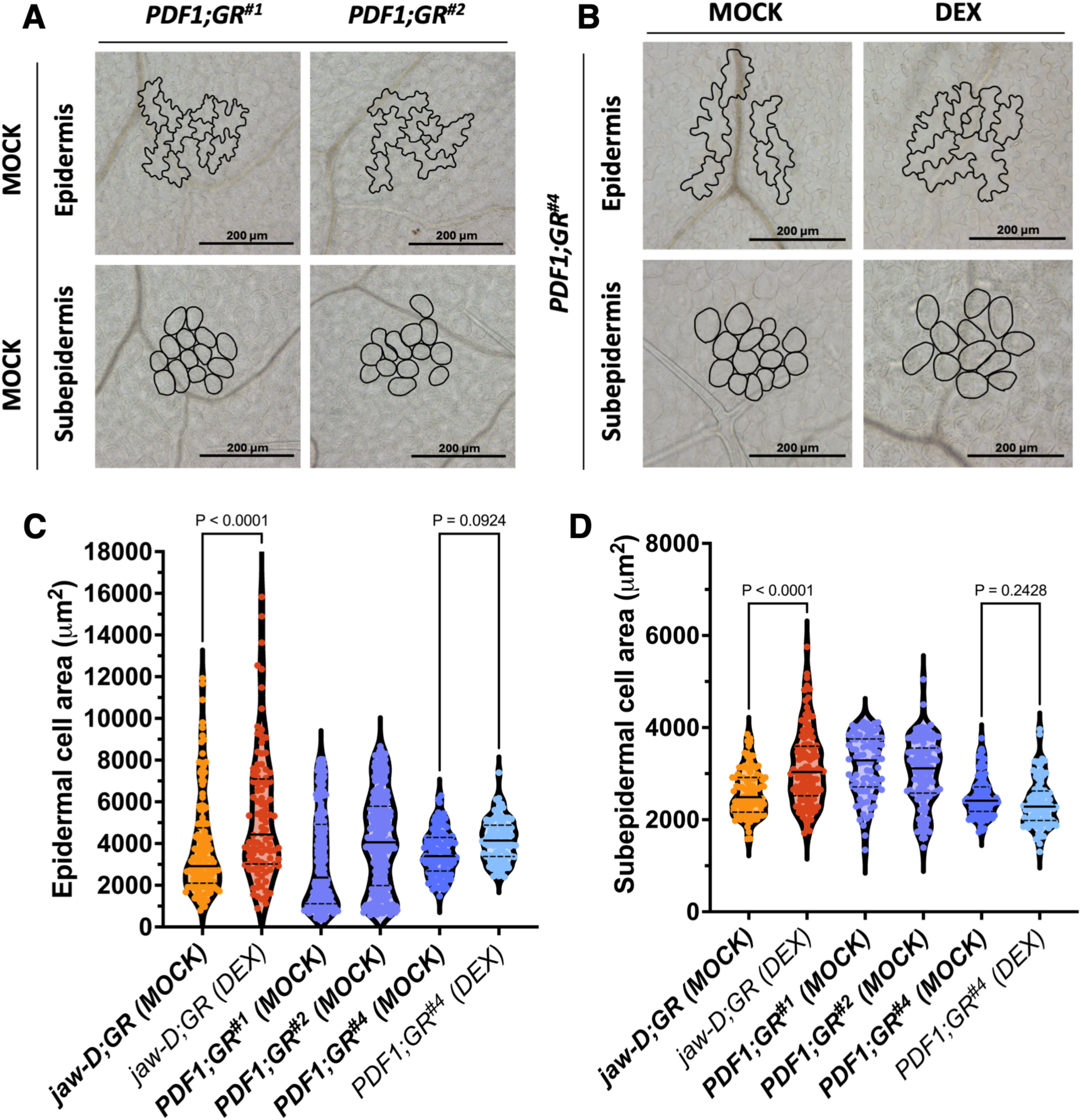
Epidermal TCP4 expression alters leaf development by modulating cell number and cell size. (A, B) Differential interference contrast (DIC) images of epidermal pavement cells and subepidermal cells from mature first leaves of 28-day-old plants of the indicated genotypes and chemical treatments. Cell outlines, including pavement cells and stomata, are shown in black. Scale bars, 200 µm. (C, D) Violin plots showing the distribution of epidermal (C) and subepidermal (D) cell areas (Y-axis) for the indicated genotypes and treatments. N = 160-600 cells. Statistical significance was assessed by one-way ANOVA followed by a Šídák post hoc test (P-values indicated above the compared means).

**Fig. S3.**
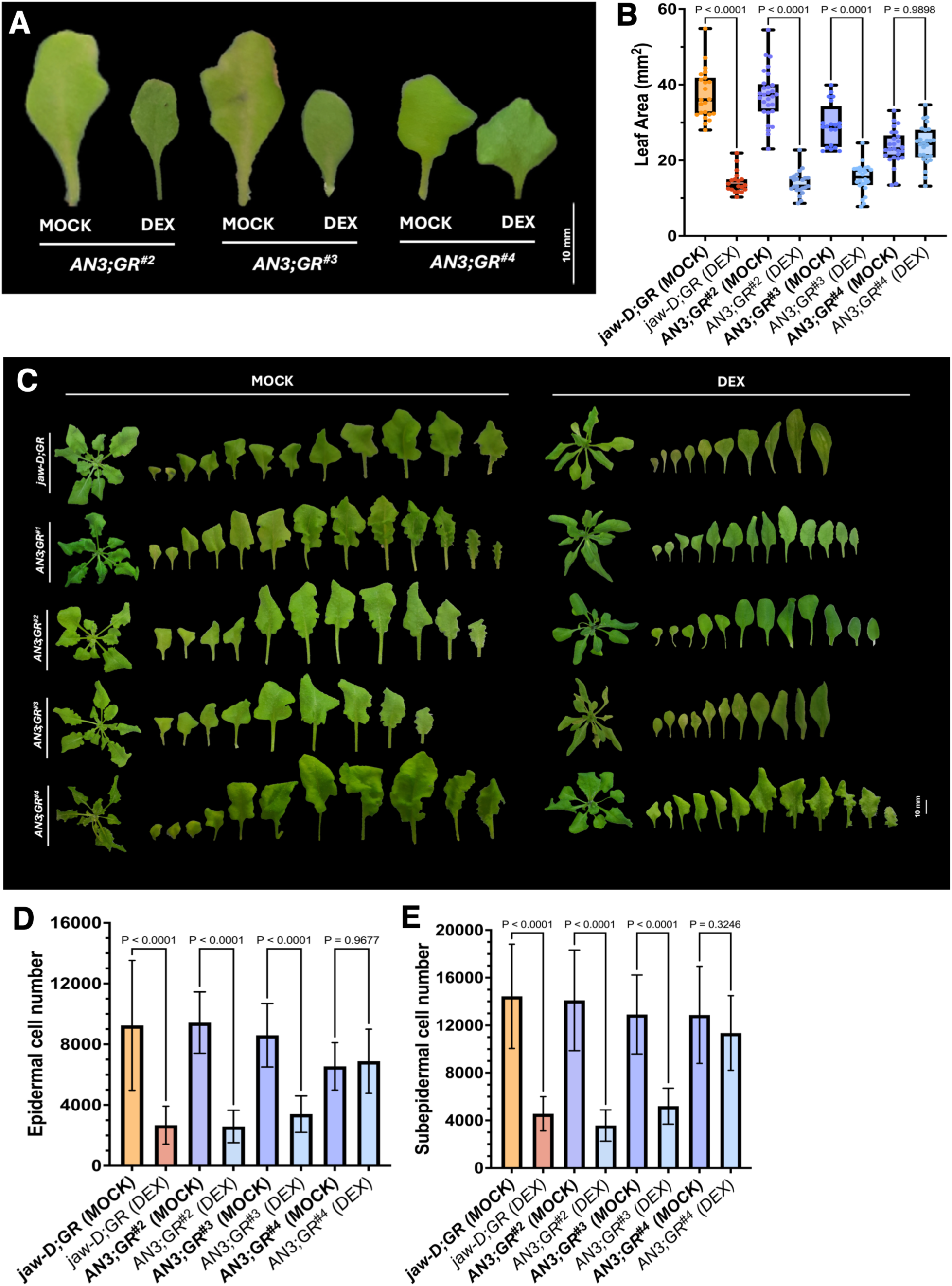
TCP4 signalling coordinates growth bidirectionally between epidermal and subepidermal layers. (A) Representative images of the mature first leaf pair from 28-day-old plants of the indicated genotypes and chemical treatments. Scale bar, 10 mm. (B) Quantification of leaf area (Y-axis) for the mature first leaf pair of the indicated genotypes and treatments (X-axis). N = 15-28 leaves. Statistical significance was assessed by one-way ANOVA followed by a Holm-Šídák post hoc test (P-values indicated above the compared means). Error bars represent SD. (C) Representative images of whole rosettes and individual rosette leaves from the indicated genotypes under MOCK and dexamethasone (DEX) treatments, corresponding to inducible TCP4 expression from the endogenous pTCP4 promoter or the subepidermal (L2) pAN3 promoter. Scale bar, 10 mm. (D, E) Estimated epidermal (D) and subepidermal (E) cell numbers for the indicated genotypes and treatments (X-axis). Cell number was calculated as the ratio of total leaf area to mean cell area (8-15 leaves per genotype). Statistical analysis as in (B); error bars represent SD.

**Fig. S4.**
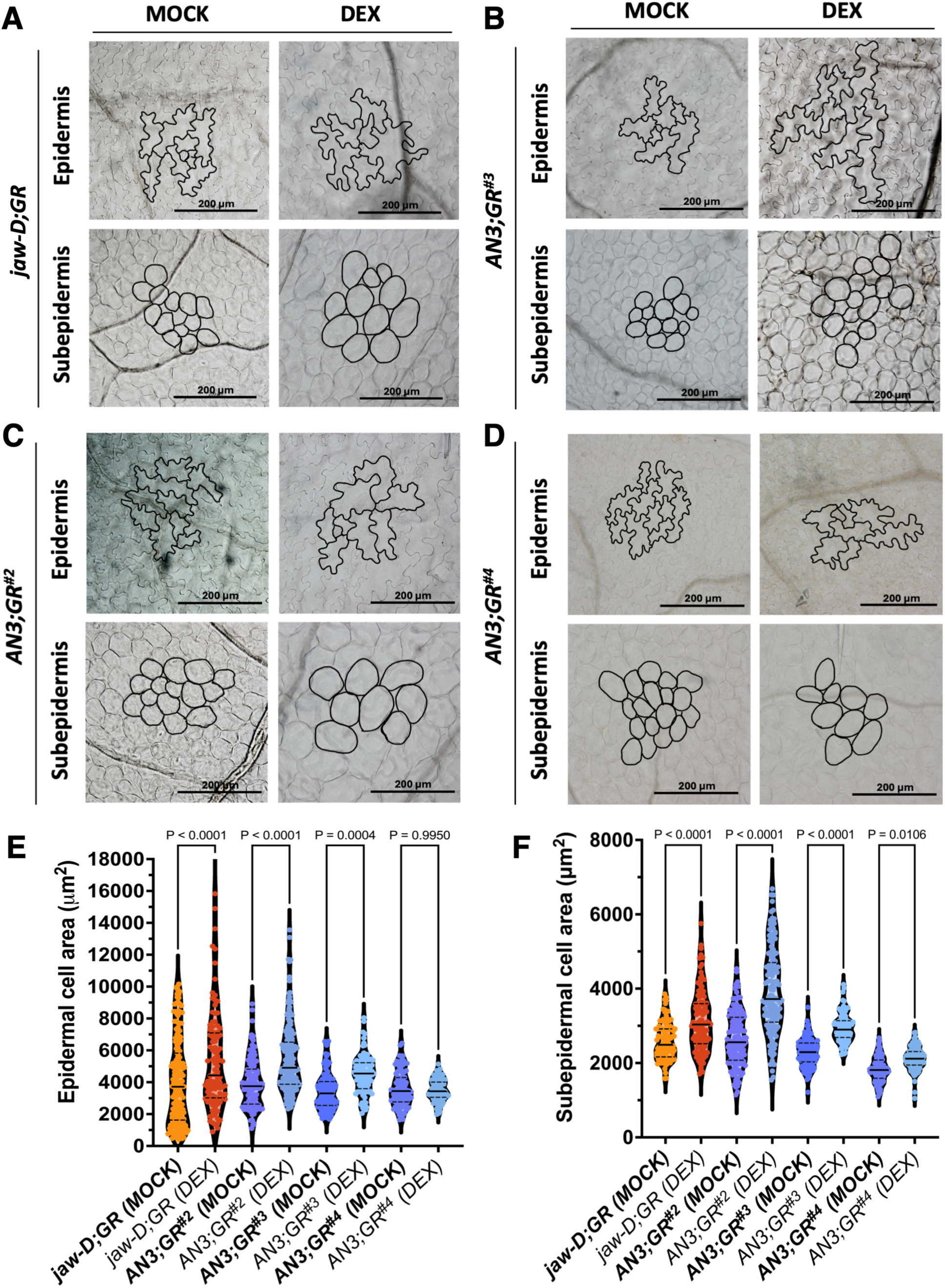
Subepidermal TCP4 expression non-autonomously regulates cell proliferation and expansion across tissue layers at near-endogenous levels. (A-D) Differential interference contrast (DIC) images of epidermal pavement cells and subepidermal cells from mature first leaves of 28-day-old plants of the indicated genotypes and chemical treatments. Cell outlines, including pavement cells and stomata, are shown in black. Scale bars, 200 µm. (E, F) Violin plots showing the distribution of epidermal (E) and subepidermal (F) cell areas (Y-axis) for the indicated genotypes and treatments. N = 120-400 cells. Statistical significance was assessed by one-way ANOVA followed by a Holm-Šídák post hoc test (P-values indicated above the compared means).

**Fig. S5.**
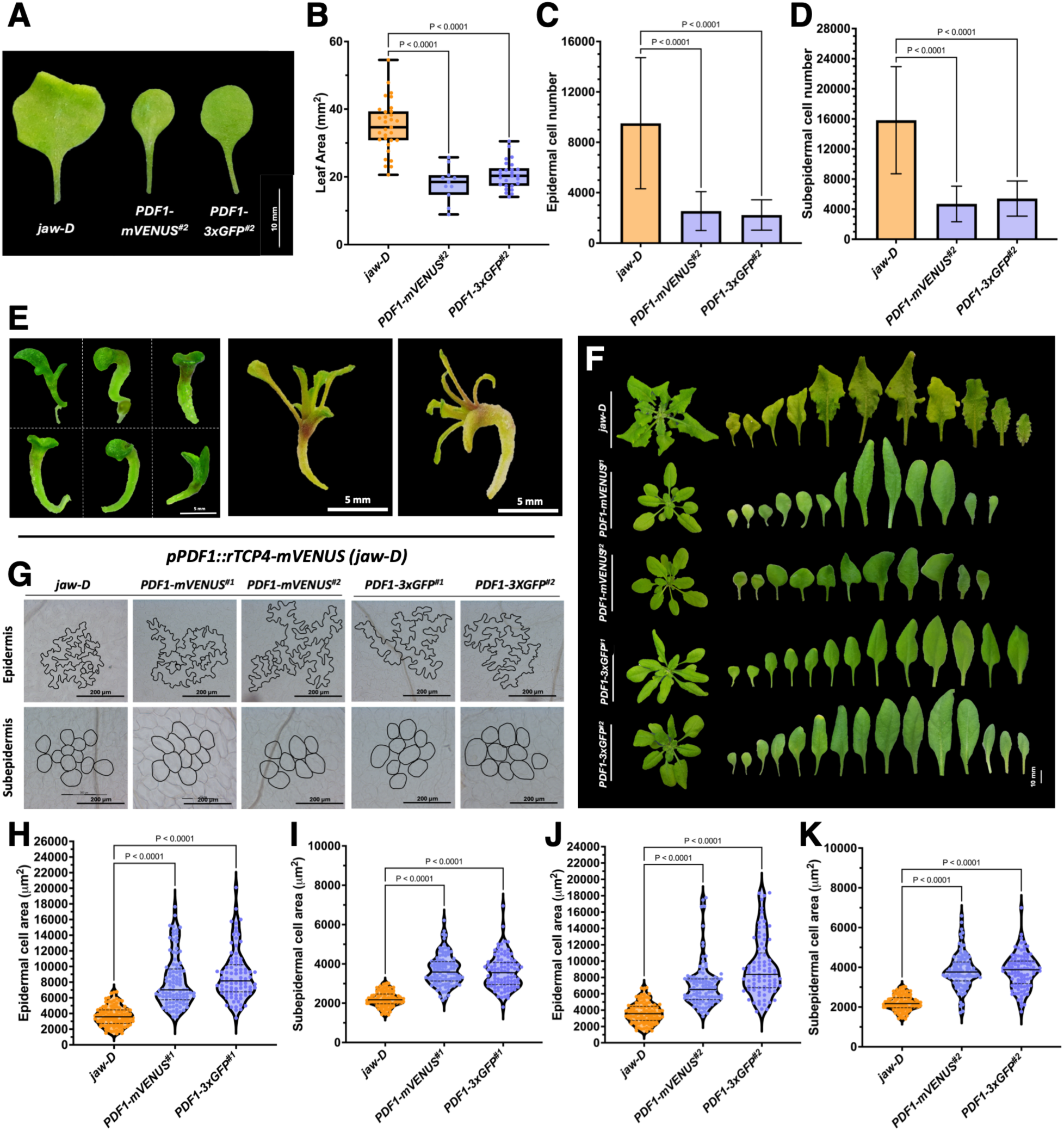
TCP4-mediated downstream signalling operates in a non-cell-autonomous manner. (A) Representative images of the mature first leaf pair from 28-day-old plants of the indicated genotypes. Scale bar, 10 mm. (B) Quantification of leaf area (Y-axis) for the mature first leaf pair of the indicated genotypes (X-axis). N = 14-26 leaves. Statistical significance was assessed by one-way ANOVA followed by a Šídák post hoc test (P-values indicated above the compared means). Error bars represent SD. (C, D) Estimated epidermal (C) and subepidermal (D) cell numbers for the indicated genotypes and chemical treatments (X-axis). Cell number was calculated as the ratio of total leaf area to mean cell area (8-10 leaves per genotype). Statistical analysis as in (B); error bars represent SD. (E) Representative T1 seedlings expressing *pPDF1::rTCP4-mVENUS* showing fused cotyledons, arrested leaf development, and seedling lethality upon constitutive epidermal TCP4 expression. Scale bar, 5 mm. (F) Representative whole rosettes and rosette leaves of homozygous weakest insertion lines *pPDF1::rTCP4-mVENUS* (#1 and #2) and *pPDF1::rTCP4-3xGFP* (#1 and #2), shown alongside the *jaw-D* control. (G) Differential interference contrast (DIC) images of epidermal pavement cells and subepidermal cells from mature first leaves of 28-day-old plants of the indicated genotypes. Cell outlines are shown in black. Scale bar, 200 µm. (H-K) Violin plots showing the distribution of epidermal (H, J) and subepidermal cell (I, K) areas for the indicated genotypes (Y-axis). N = 120-400 cells. Statistical significance was assessed by one-way ANOVA followed by a Šídák post hoc test (P-values indicated above the compared means).

**Fig. S6.**
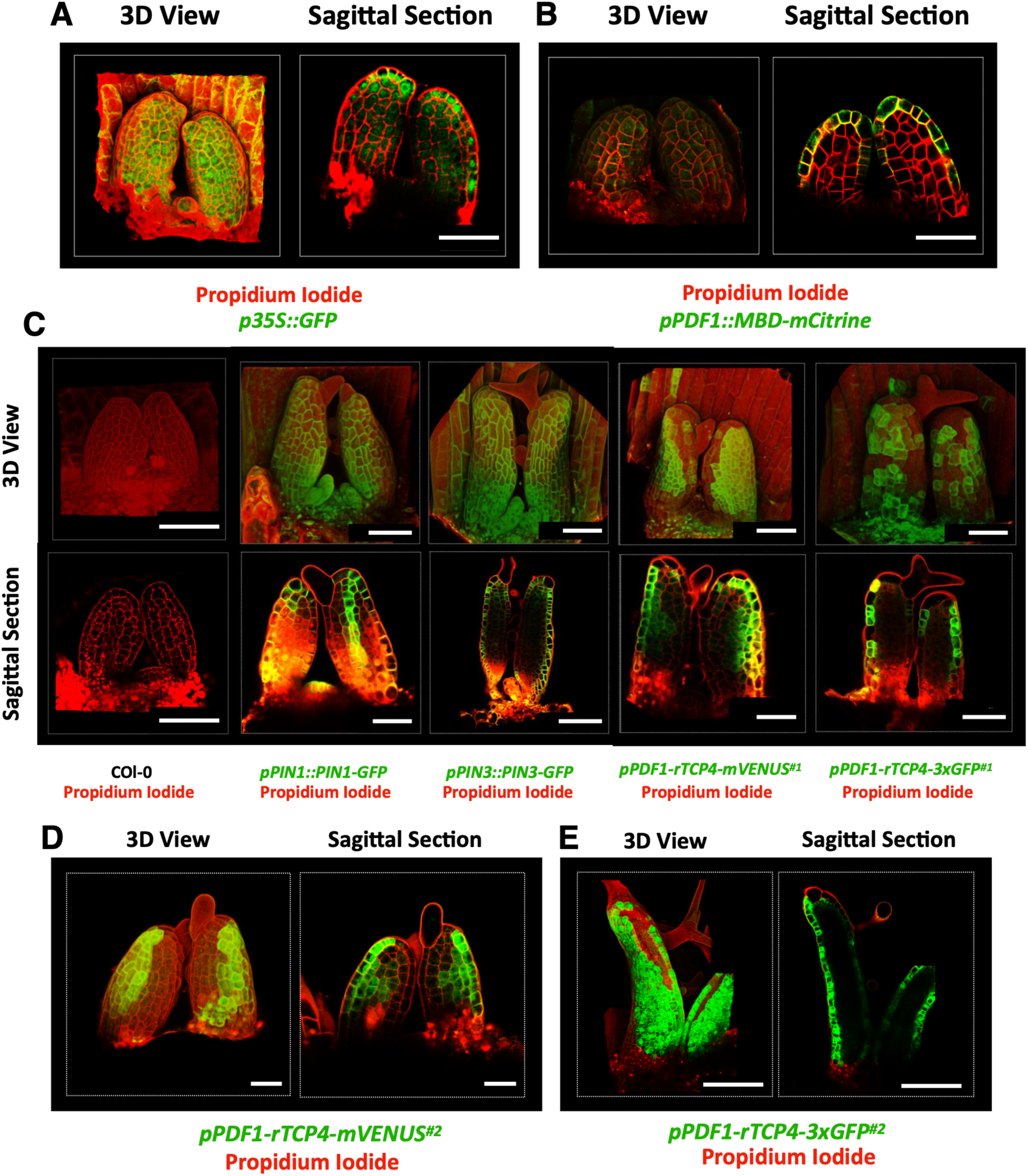
TCP4 protein is mobile across tissue layers, whereas a bulky fusion restricts its movement. (A, B) Confocal images of propidium iodide-stained leaf primordia (2-3 DAI) from control lines. (A) *p35S::GFP* showing fluorescence in all tissue layers. (B) *pPDF1::MBD-mCitrine* showing fluorescence restricted to the epidermis. Three-dimensional reconstructions and optical mid-sagittal sections are shown. Scale bars, 50 µm. (C) Propidium iodide-stained confocal images of first leaf primordia (2-3 DAI) are shown as three-dimensional reconstructions and optical sagittal sections. Negative control (Col-0) shows no signal. Positive controls *pPIN1::PIN1-GFP* and *pPIN3::PIN3-GFP* display expected localisation patterns in L1/vasculature and all layers, respectively. In comparison*, pPDF1::rTCP4-mVENUS* (#1 and #2) shows fluorescence in the epidermis and adjacent subepidermal layers, whereas *pPDF1::rTCP4-3xGFP* (#1 and #2) fluorescence is confined to the epidermis. Scale bars, 50 µm. (D) Confocal images of propidium iodide-stained leaf primordia (2-3 DAI) from *pPDF1::rTCP4-mVENUS* insertion lines (#2). Three-dimensional views and optical sagittal show fluorescence in the epidermis and adjacent subepidermal layers, indicating limited inter-layer mobility of TCP4-mVENUS. Scale bars, 50 µm. (E) Propidium iodide-stained confocal images of first leaf primordia (5-6 DAI) from *pPDF1::rTCP4-3xGFP* (#2) are shown as three-dimensional views and optical mid-sagittal sections, confirming epidermis-restricted localisation at later developmental stages. Scale bars, 100 µm.

**Fig. S7.**
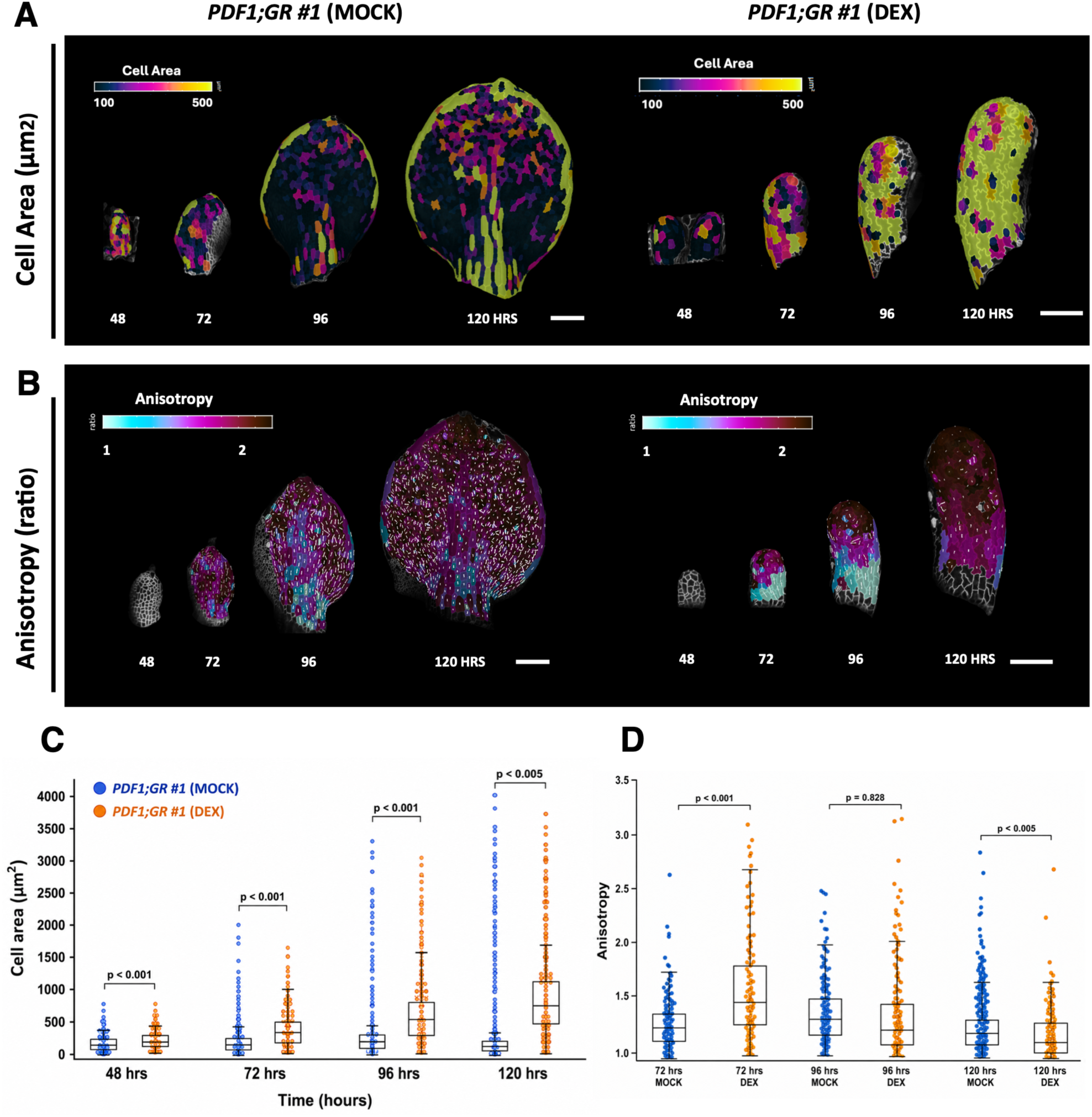
Epidermal TCP4 induction remodels cell area expansion and growth anisotropy during leaf morphogenesis. (A) Heat maps depicting the area extension cellular area profile of the leaf primordia (*pPDF::rTCP4-GR #1* line in MOCK and DEX conditions) live-imaged from 48-120 hours after initiation. (B) Heat maps of growth anisotropy superimposed with vectors depicting principal directions of growth (max PDGs) calculated using live imaging (*pPDF1; GR-*MOCK and DEX) from 48-120 hours post leaf initiation. (C) Boxplots of area extension cell area (µm²) of three replicates of leaf primordium of *pPDF::rTCP4-GR #1* line in MOCK (blue) and DEX (orange) conditions. A Student’s t-test was performed between the respective MOCK and DEX treatments across the time course, and p-values are indicated above the box plots. (D) Boxplots showing growth anisotropy of three replicates of leaf primordium of *pPDF::rTCP4-GR* line in MOCK (blue) and DEX (orange) conditions. A Student’s t-test was performed between the respective MOCK and DEX treatments across the time course, and p-values are indicated above the box plots.

**Fig. S8.**
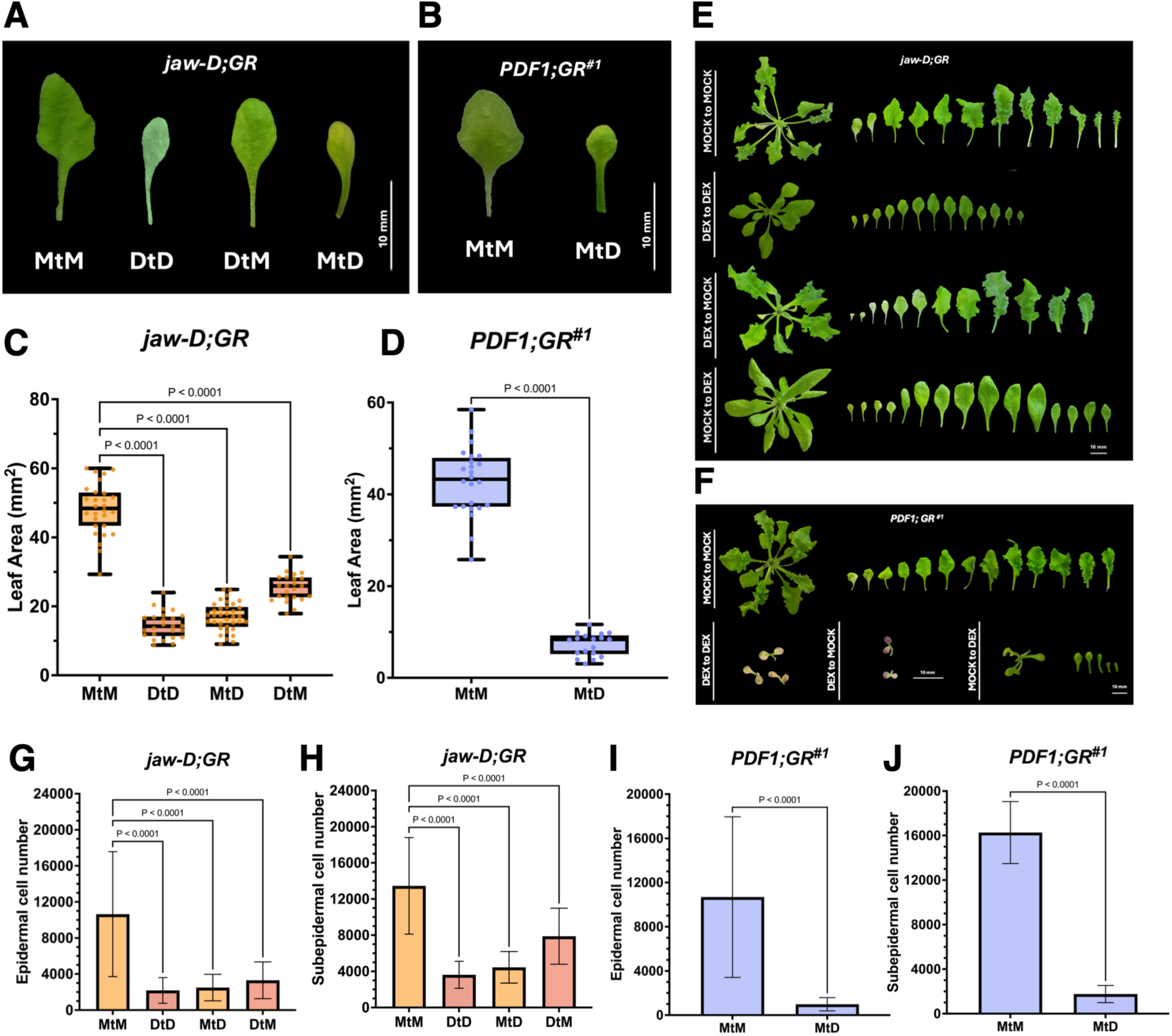
Ectopic epidermal TCP4 induction inhibits leaf formation in a temporally sensitive manner. (A, B) Mature first leaves from 28-day-old plants of *jaw-D;GR* (*pTCP4::rTCP4-GR*) and *pPDF1;GR* (#1; *pPDF1::rTCP4-GR*) grown under staged dexamethasone (DEX; 12 μM) treatments. Seedlings were grown under MOCK conditions for the first 4 days and then transferred to MOCK (MtM) or DEX (MtD). Alternatively, seedlings were grown in DEX for the first 4 days and then transferred to DEX (DtD) or MOCK (DtM). Scale bar, 10 mm. (C, D) Quantification of leaf area (Y-axis) of the mature first leaf pair for *jaw-D;GR*, *pPDF1;GR* #1 under the indicated transfer treatments (X-axis). N = 20-34 leaves. Statistical significance was assessed by one-way ANOVA followed by a Šídák post hoc test (P-values indicated over the compared treatments). Error bars represent SD. (E) Representative whole rosettes and individual rosette leaves of *jaw-D;GR* plants under MOCK and continuous DEX treatment, corresponding to induction of TCP4 under its endogenous promoter (pTCP4) in the jaw-D background. Scale bar, 10 mm. (F) Representative whole rosettes and individual rosette leaves of *pPDF1;GR* #1 plants under MOCK and DEX and transfer treatments, corresponding to epidermis-specific induction of TCP4 in the *jaw-D* background. Scale bar, 10 mm. (G-I) Estimated epidermal and subepidermal cell numbers (Y-axis) for the indicated genotypes under the respective chemical transfer treatments (X-axis). Cell number was calculated as the ratio of total leaf area to mean cell area, based on 12-20 leaves per genotype. Statistical significance was assessed by one-way ANOVA followed by a Šídák post hoc test (P-values indicated over the compared treatments). Error bars represent SD.

**Fig. S9.**
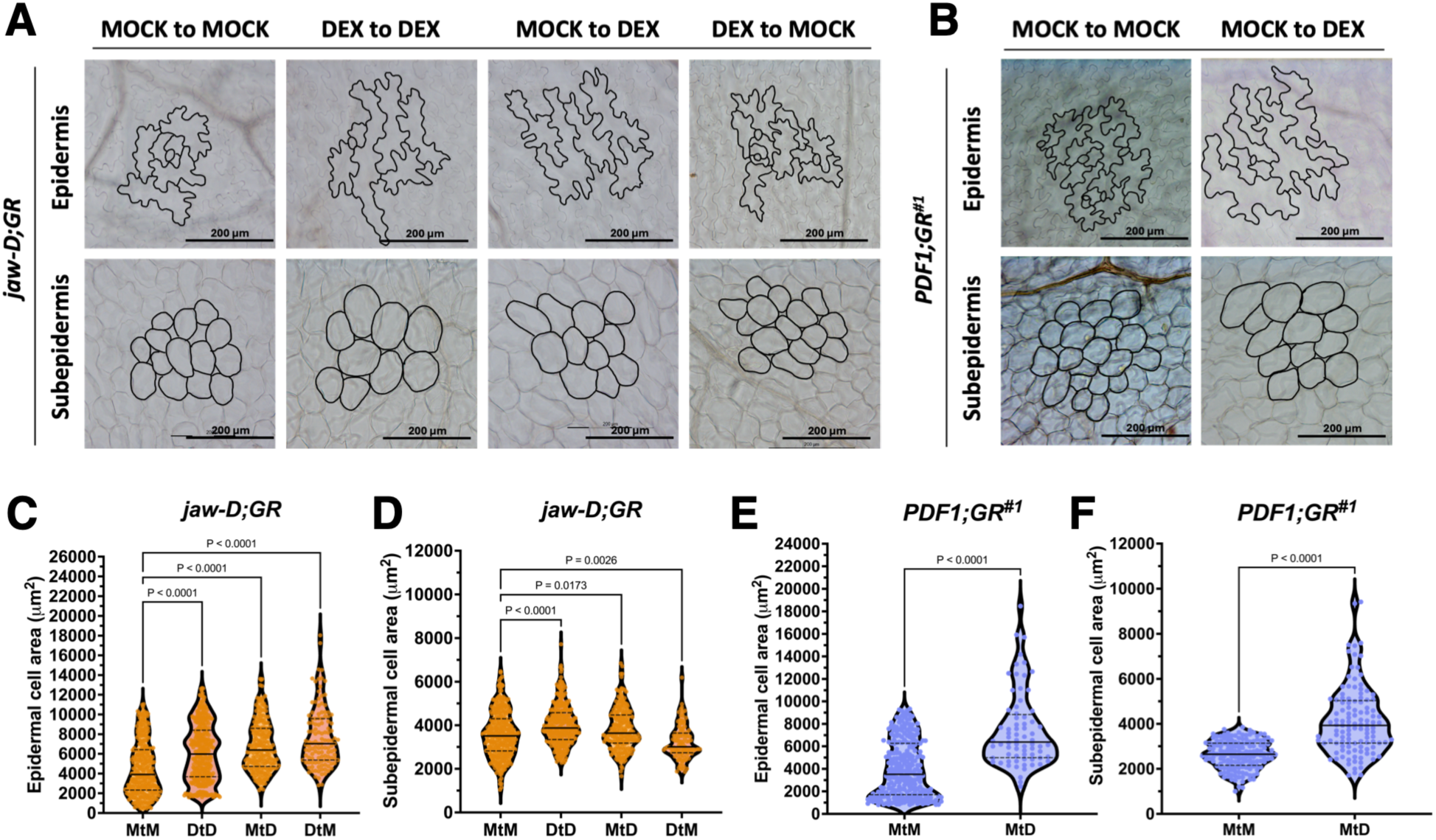
Epidermal TCP4 expression dominantly shapes cellular area through non-autonomous regulation across tissue layers. (A, B) Differential interference contrast (DIC) images of epidermal pavement cells and subepidermal cells from mature first leaves of 28-day-old plants of the indicated genotypes and chemical transfer treatments. Cell outlines, including pavement cells and stomata, are shown in black. Scale bars, 200 µm. (C-F) Violin plots showing the distribution of epidermal and subepidermal cell areas (Y-axis) for the indicated genotypes and transfer treatments. N = 120-400 cells. Statistical significance was assessed by one-way ANOVA followed by a Šídák post hoc test (P-values indicated over the compared treatments). Error bars represent SD.

**Fig. S10.**
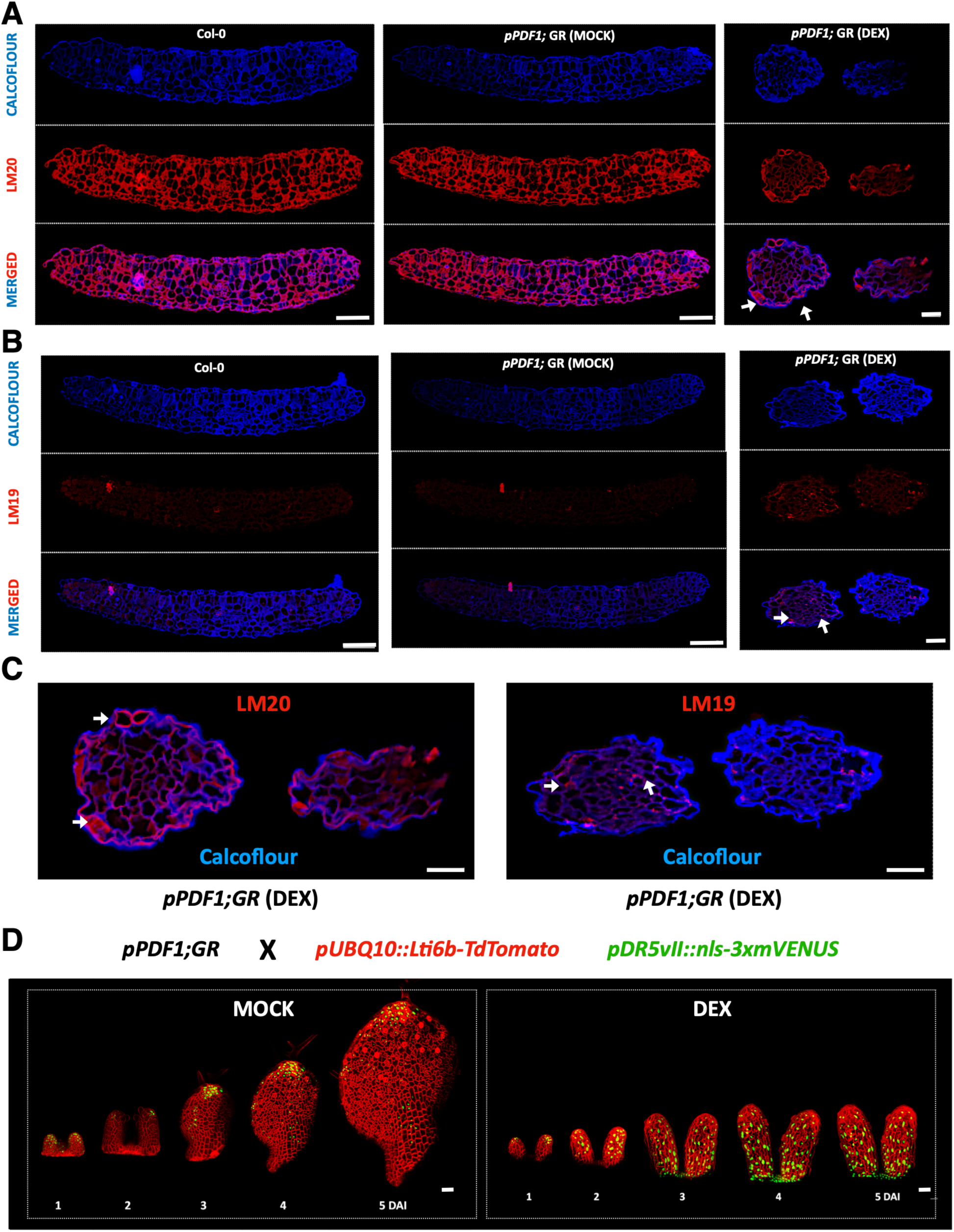
Epidermal TCP4 induction triggers sustained auxin signalling and alters pectin-associated mechanical signatures. (A, B) Confocal images of transverse sections of 6 DAI leaves (N=4) from Col-0, and *pPDF1::rTCP4-GR* leaves under MOCK and DEX conditions, immunolabeled with calcofluor (cell wall marker) and antibodies recognising pectin modifications. LM20 labels highly methylesterified pectin, whereas LM19 labels demethylesterified pectin. Merged images with calcofluor are shown. Arrows indicate ectopic deposition of LM20 in the epidermis of *pPDF1::rTCP4-GR DEX* samples (A) and increased LM19 signal in the subepidermal layer (B). Scale bars, 20 µm. (C) Zoomed inset of *pPDF1::rTCP4-GR* (DEX) depicting localisation of LM19 and LM20 antibodies/calcofluor. Arrows indicate ectopic deposition of LM20 in the epidermis of *pPDF1::rTCP4-GR DEX* samples and increased LM19 punctate signal in the subepidermal layer. Scale bars, 20 µm. (D) Time-lapse confocal imaging of the first leaf from *pPDF1::rTCP4-GR* crossed to *pUBQ10::Lti6b-TdTomato* (plasma membrane marker) and *pDR5vII::3xmVENUS* (auxin response reporter). Seedlings were imaged every 24 hours from 1 to 5 days after initiation (DAI) under MOCK and dexamethasone (DEX) treatments. DEX-treated primordia display sustained auxin response compared with MOCK controls. Scale bar, 50 µm.

**Supplementary Table S1:**
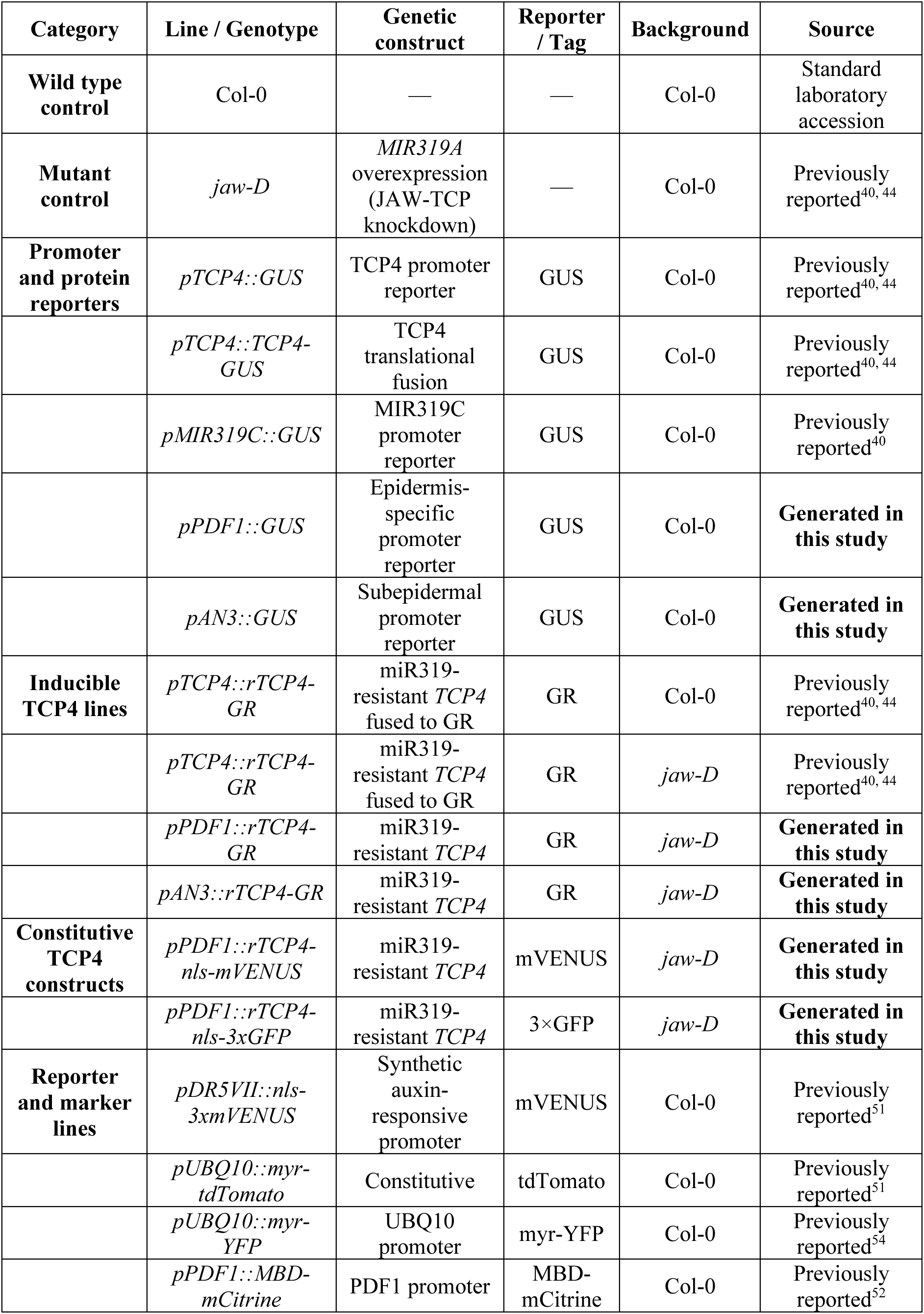
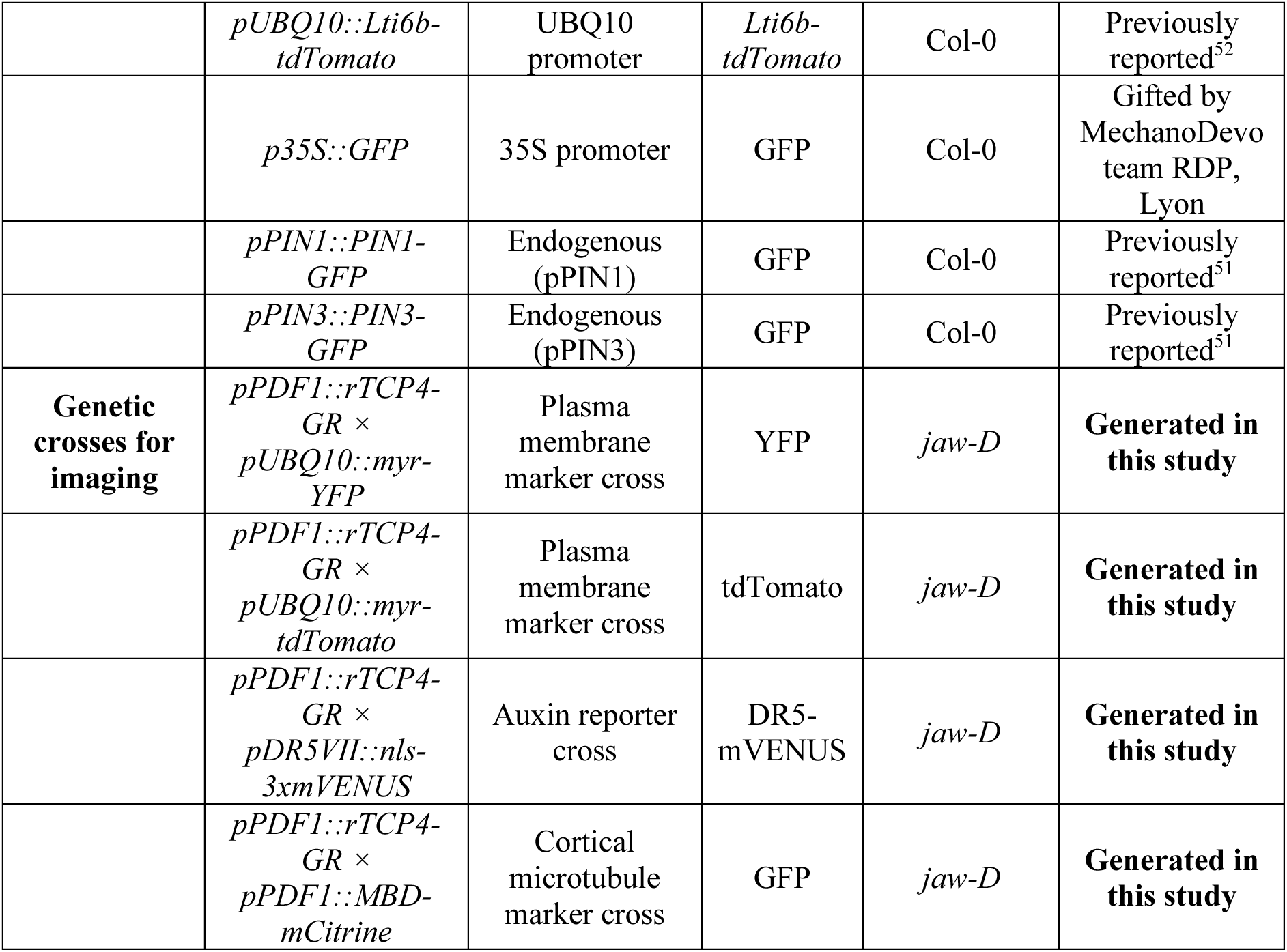
Plant materials used in this study.

**Supplementary Table S2:**
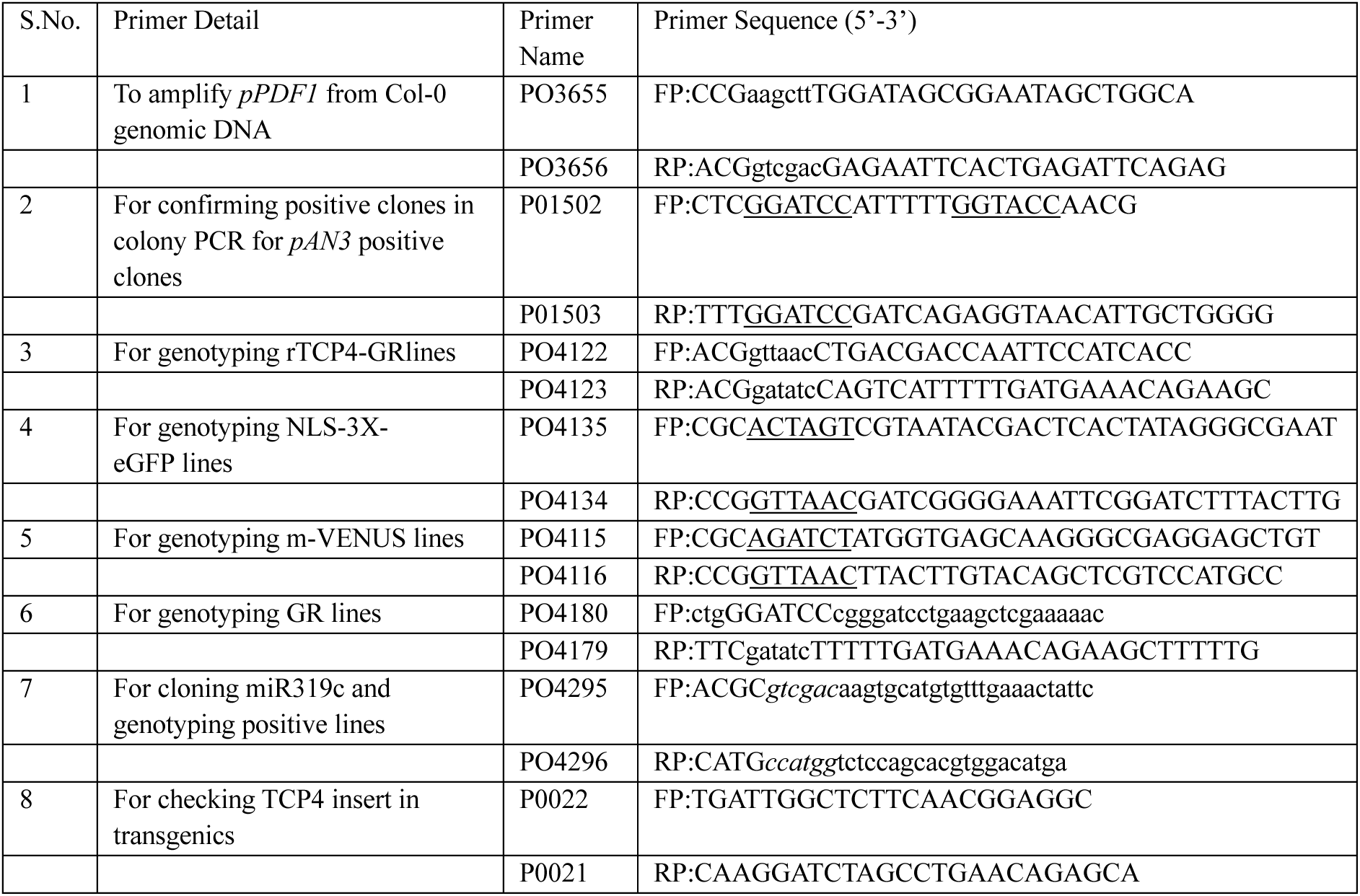
Primers used in this study.

**Supplementary Table S3:**
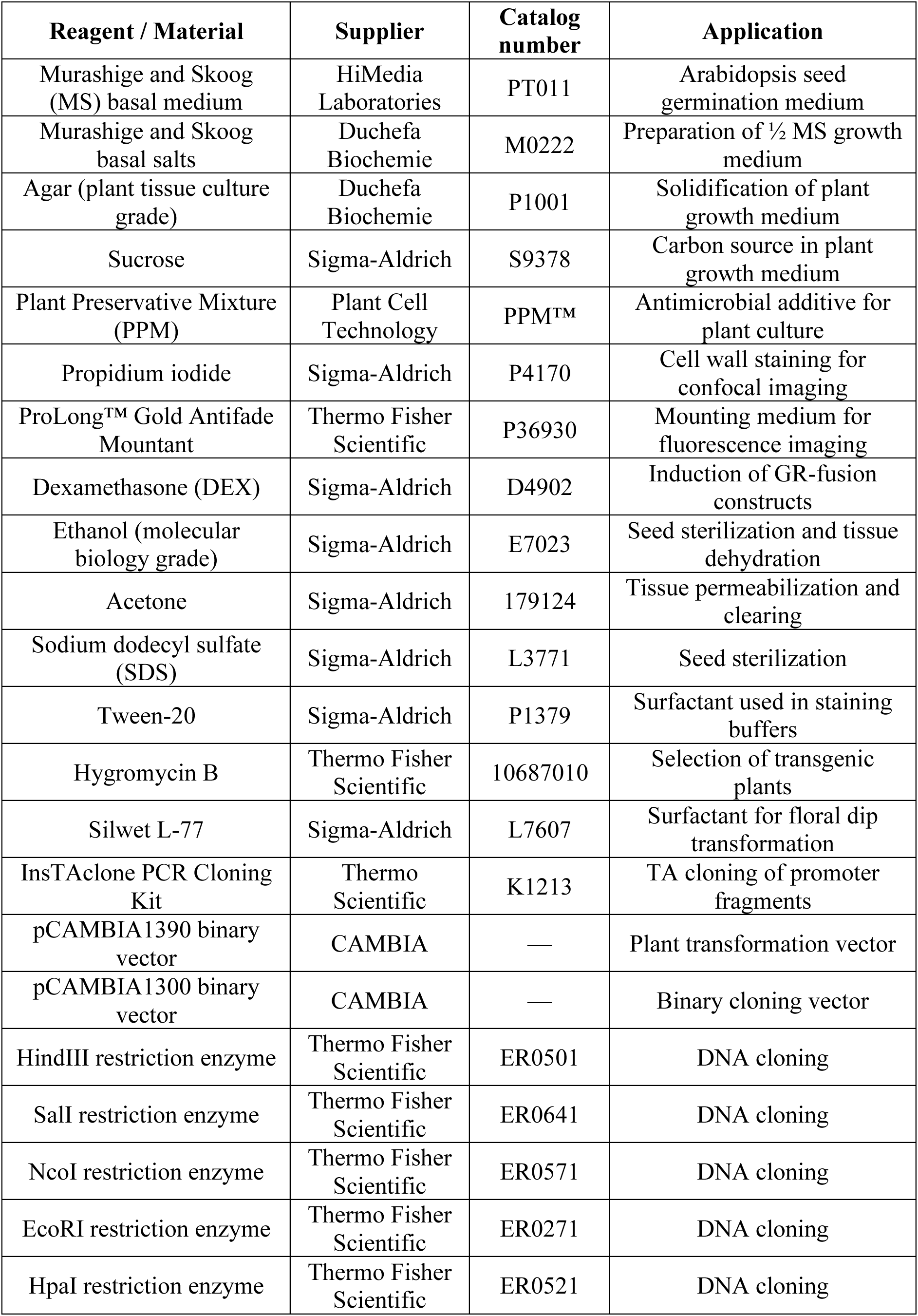
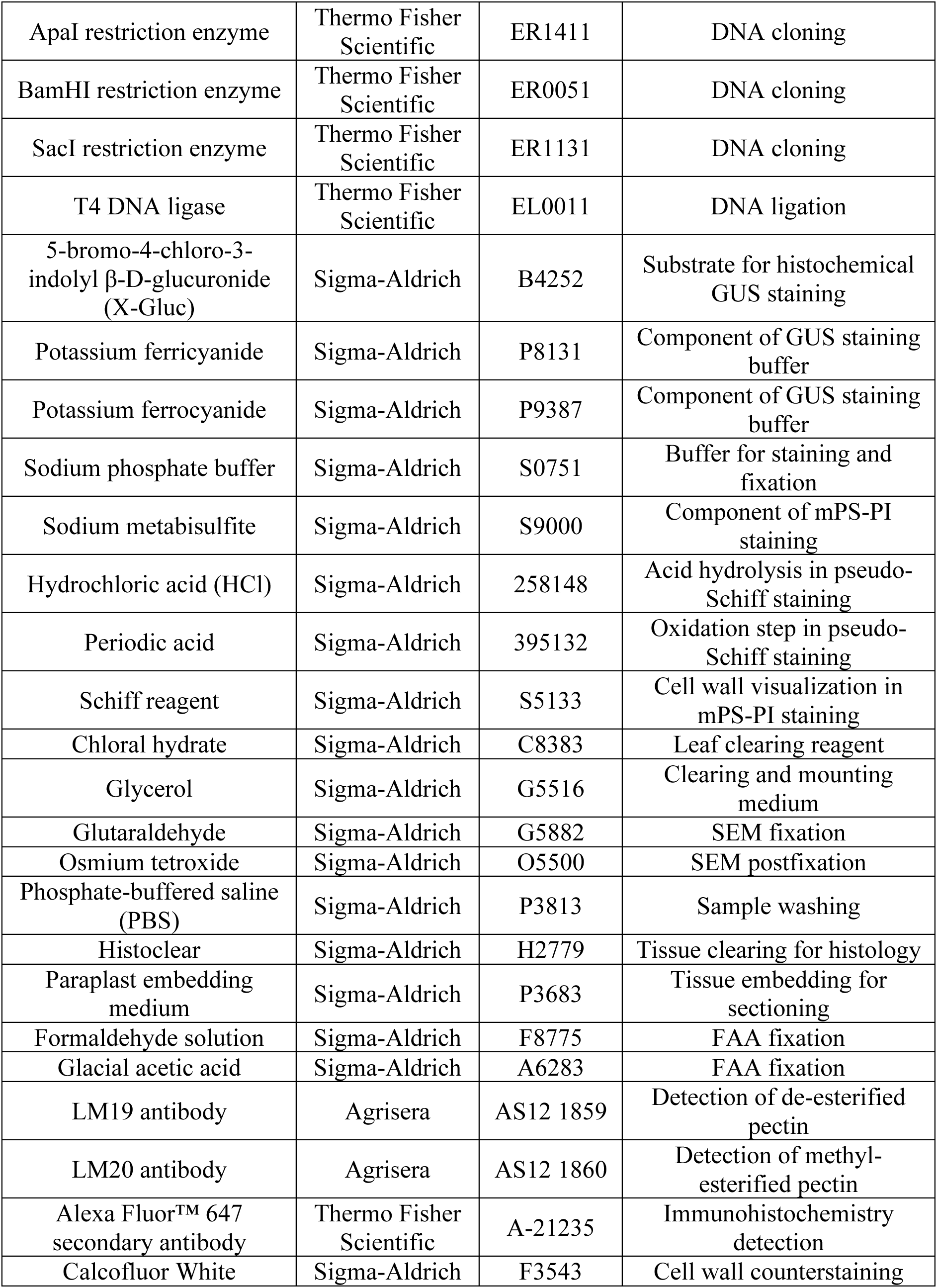
Reagents used in this study.

**Supplementary Table S4:**
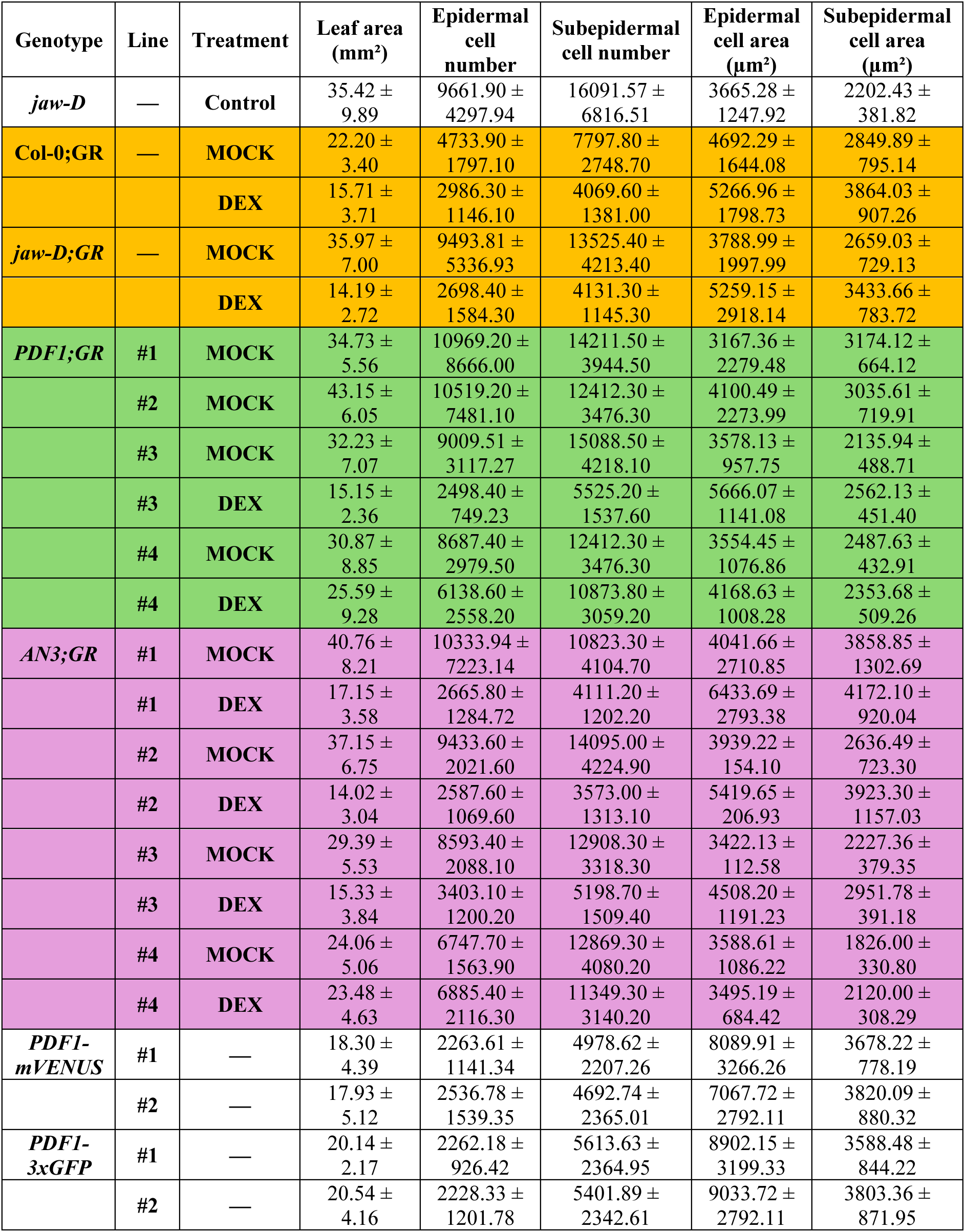
Morphometric measurements of leaf size, epidermal and subepidermal cellular parameters across genotypes and treatments (mean along with SD).

**Supplementary Table S5:**
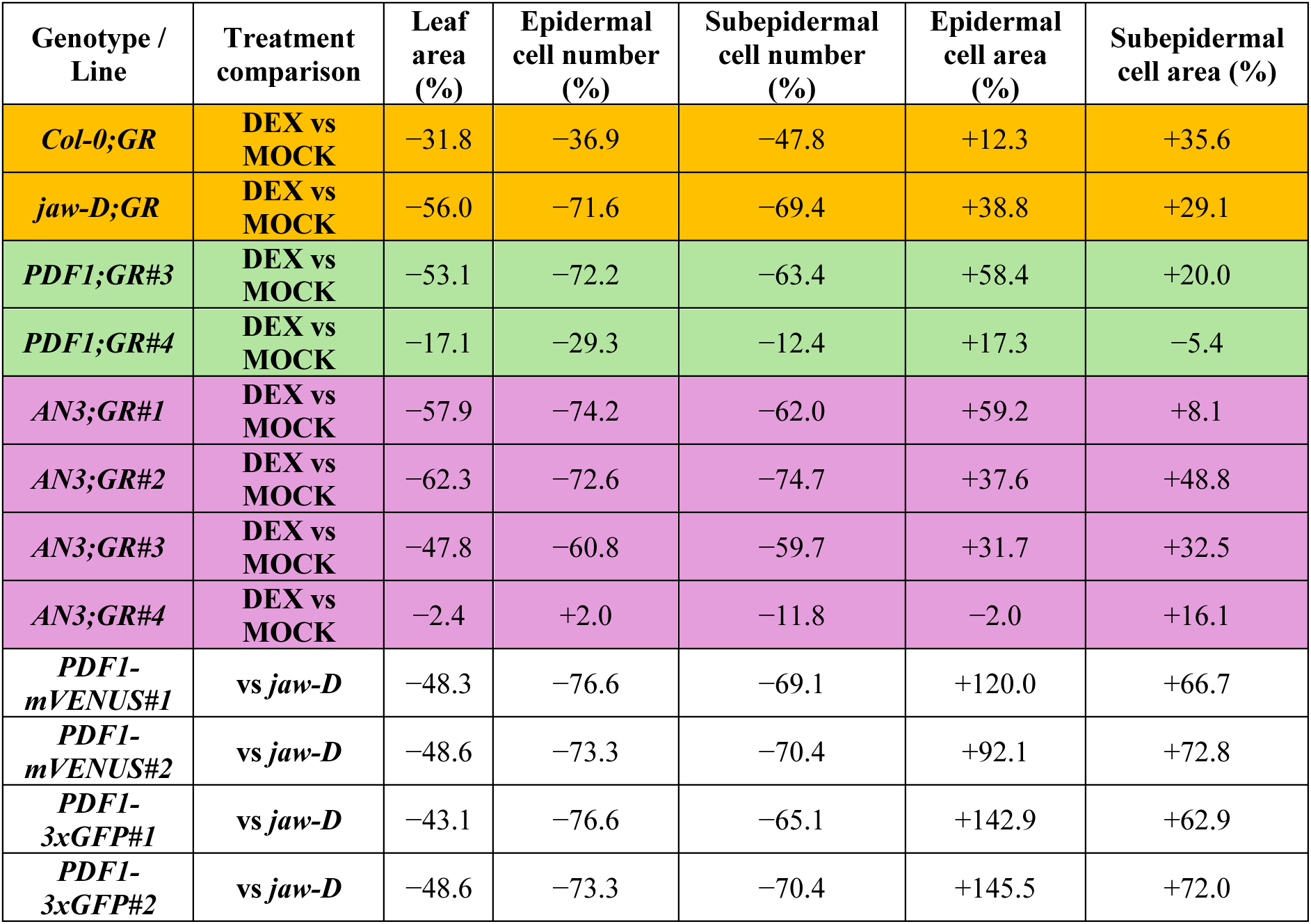
Percentage change in leaf morphometric parameters relative to respective controls.

**Supplementary Table S6:**
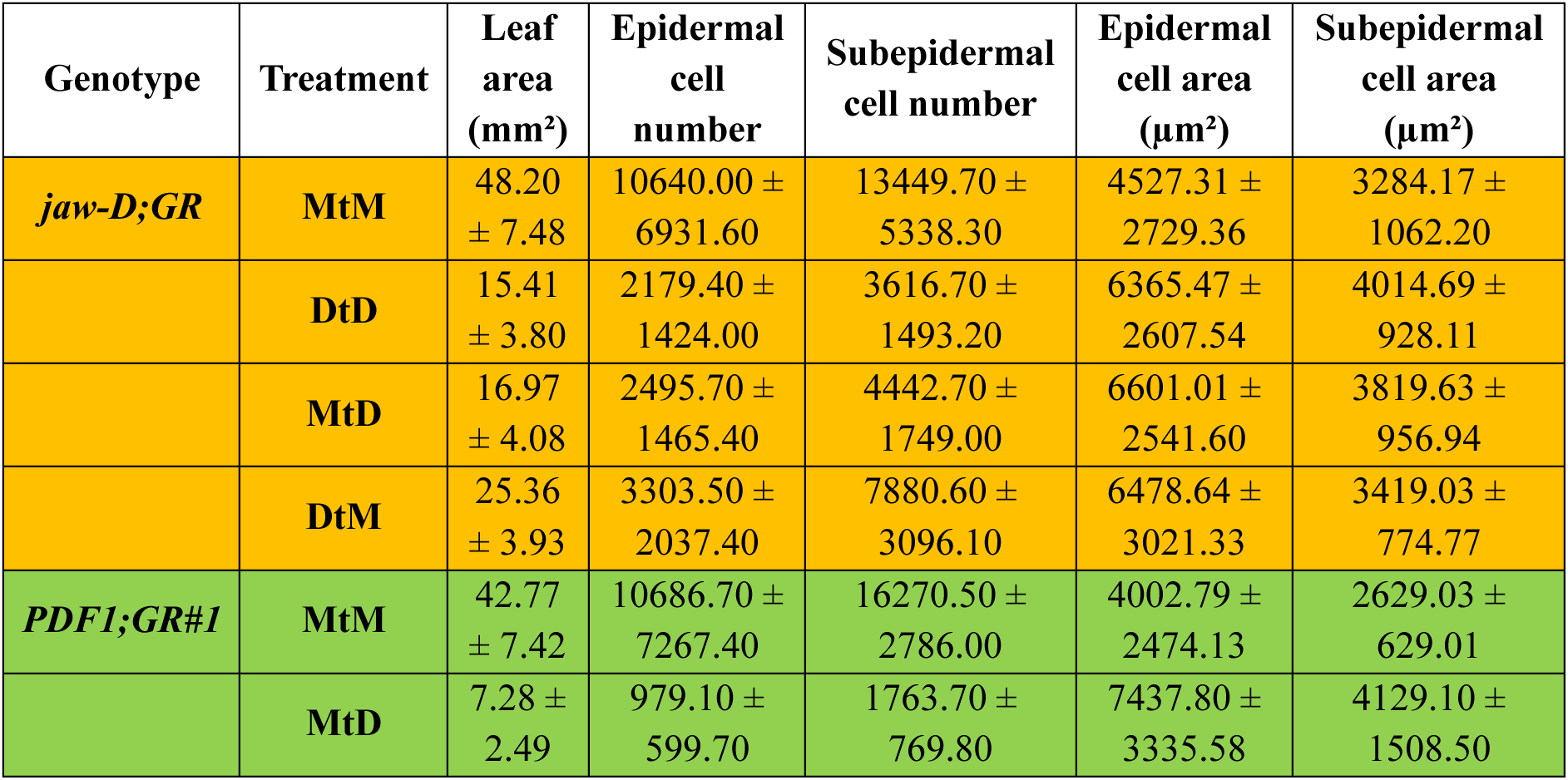
Absolute morphometric measurements for transfer experiments.

**Supplementary Table S7:**
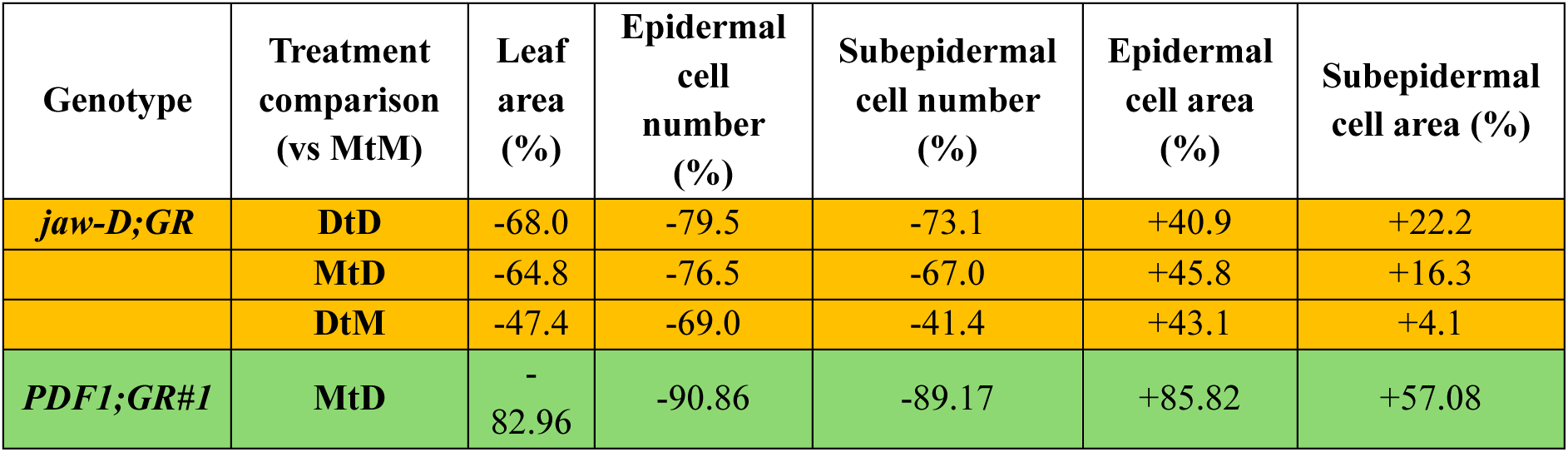
Percentage change in leaf morphometric parameters in transfer experiments relative to MtM control.

