## Supplementary files for "Control of leaf shape through non-cell autonomous TCP4 regulation"

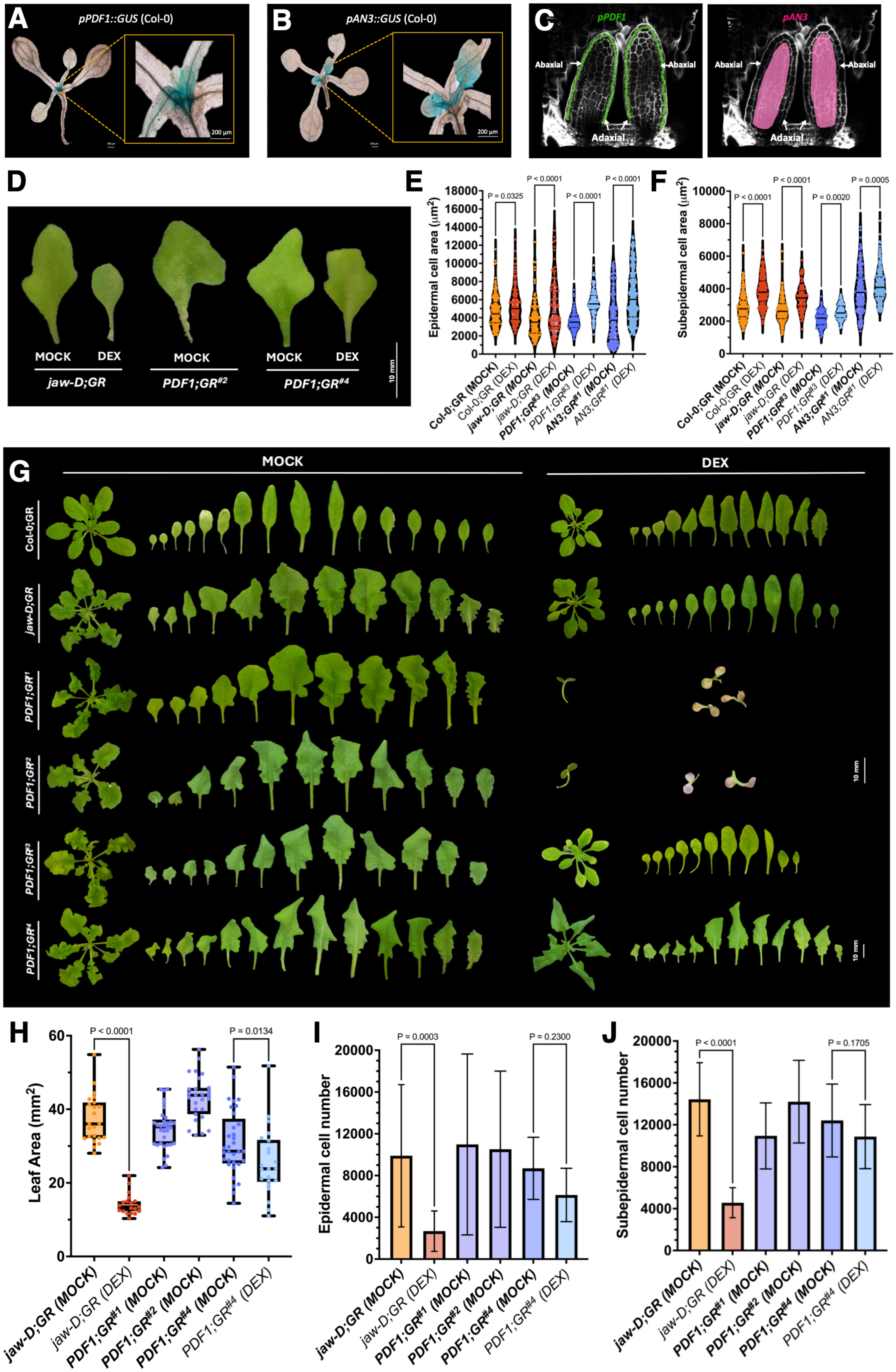
A
pPDF1::GUS (Col-0)
200 µm
B
pAN3::GUS (Col-0)
200 µm
C
pPDF1
Abaxial
Abaxial
Adaxial
pAN3
Abaxial
Abaxial
Adaxial
D
MOCK
DEX
jaw-D;GR
MOCK
PDF1;GR#2
MOCK
DEX
PDF1;GR#4
10 mm
E
Epidermal cell area (µm2)
P = 0.0325
P < 0.0001
P < 0.0001
P < 0.0001
Col-0;GR (MOCK)
Col-0;GR (DEX)
jaw-D;GR (MOCK)
jaw-D;GR (DEX)
PDF1;GR#3 (MOCK)
PDF1;GR#3 (DEX)
AN3;GR#1 (MOCK)
AN3;GR#1 (DEX)
F
Subepidermal cell area (µm2)
P < 0.0001
P < 0.0001
P = 0.0020
P = 0.0005
Col-0;GR (MOCK)
Col-0;GR (DEX)
jaw-D;GR (MOCK)
jaw-D;GR (DEX)
PDF1;GR#3 (MOCK)
PDF1;GR#3 (DEX)
AN3;GR#1 (MOCK)
AN3;GR#1 (DEX)
G
MOCK
DEX
Col-0;GR
jaw-D;GR
PDF1;GR1
PDF1;GR2
PDF1;GR3
PDF1;GR4
10 mm
10 mm
H
Leaf Area (mm2)
P < 0.0001
P = 0.0134
jaw-D;GR (MOCK)
jaw-D;GR (DEX)
PDF1;GR#1 (MOCK)
PDF1;GR#2 (MOCK)
PDF1;GR#4 (MOCK)
PDF1;GR#4 (DEX)
I
Epidermal cell number
P = 0.0003
P = 0.2300
jaw-D;GR (MOCK)
jaw-D;GR (DEX)
PDF1;GR#1 (MOCK)
PDF1;GR#2 (MOCK)
PDF1;GR#4 (MOCK)
PDF1;GR#4 (DEX)
J
Subepidermal cell number
P < 0.0001
P = 0.1705
jaw-D;GR (MOCK)
jaw-D;GR (DEX)
PDF1;GR#1 (MOCK)
PDF1;GR#2 (MOCK)
PDF1;GR#4 (MOCK)
PDF1;GR#4 (DEX)

**Fig. S1. Promoter-specific TCP4 induction reveals layer-dependent control of cell proliferation and expansion in leaf size rescue.** (A, B) Bright-field images of whole-mount leaves with corresponding magnified insets showing GUS reporter activity driven by the epidermis-specific promoter *pPDF1* (A) and the mesophyll-associated promoter *pAN3* (B) in 8-day-old seedlings. Scale bars, 200 µm. (C) Schematic representation of *pPDF1* (green) and *pAN3* (magenta) expression domains superimposed on an optical sagittal section of a developing leaf (grey), illustrating their relative tissue-layer localisation. (D) Representative images of the mature first leaf pair from 28-day-old plants of the indicated genotypes and chemical treatments. Scale bar, 10 mm. (E, F) Violin plots showing the distribution of epidermal (E) and subepidermal (F) cell areas for the indicated genotypes and treatments. N = 180-600 cells. Statistical significance was assessed by one-way ANOVA followed by a Šídák post hoc test (P-values indicated above the compared means). (G) Representative images of whole rosettes and individual rosette leaves from the indicated genotypes under MOCK and dexamethasone (DEX) treatments, corresponding to inducible TCP4 expression from the endogenous pTCP4 promoter or the epidermis-specific pPDF1 promoter. Scale bar, 10 mm. (H) Quantification of leaf area (Y-axis) for the mature first leaf pair of the indicated genotypes and treatments (X-axis). N = 20-34 leaves. Statistical analysis as in (E, F). Error bars represent SD. (I, J) Estimated epidermal (I) and subepidermal (J) cell numbers for the indicated genotypes and treatments. Cell number was calculated as the ratio of total leaf area to mean cell area (10-13 leaves per genotype). Statistical analysis as in (E, F); error bars represent SD. P-values are indicated above the compared means.

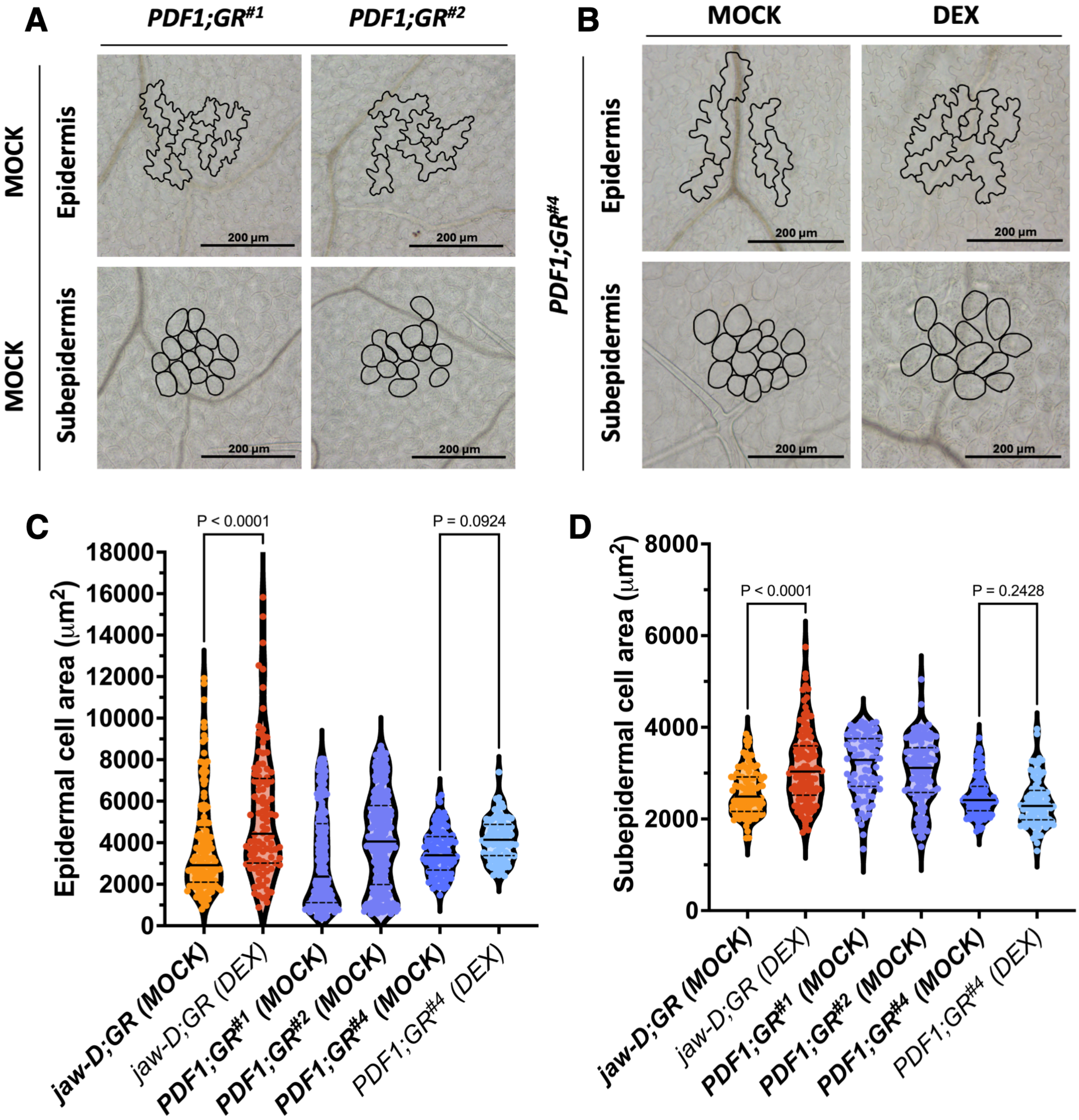


**Fig. S2. Epidermal TCP4 expression alters leaf development by modulating cell number and cell size.** (A, B) Differential interference contrast (DIC) images of epidermal pavement cells and subepidermal cells from mature first leaves of 28-day-old plants of the indicated genotypes and chemical treatments. Cell outlines, including pavement cells and stomata, are shown in black. Scale bars, 200 µm. (C, D) Violin plots showing the distribution of epidermal (C) and subepidermal (D) cell areas (Y-axis) for the indicated genotypes and treatments. N = 160-600 cells. Statistical significance was assessed by one-way ANOVA followed by a Šídák post hoc test (P-values indicated above the compared means).

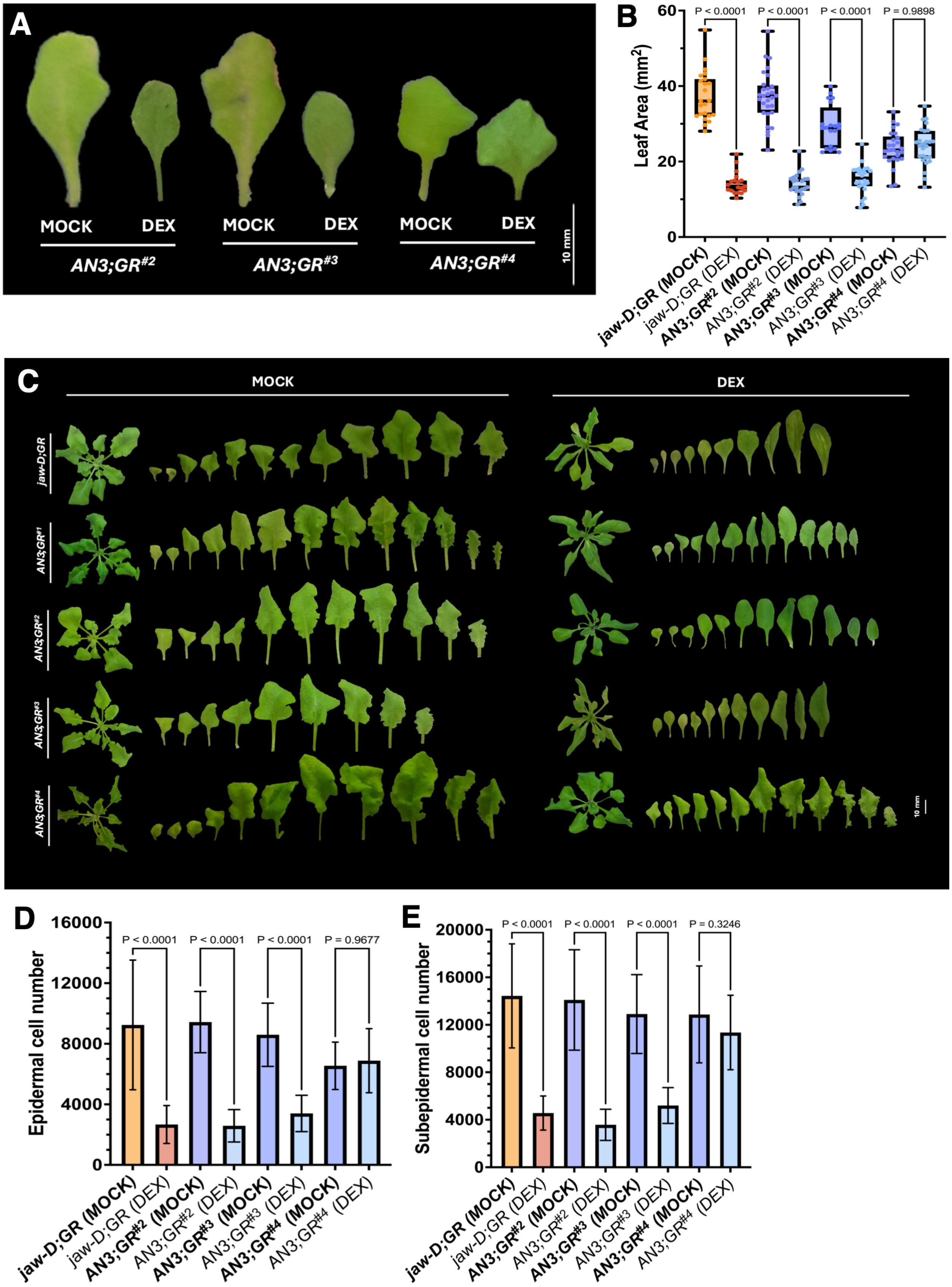

A
MOCK
DEX
MOCK
DEX
MOCK
DEX
AN3;GR#2
AN3;GR#3
AN3;GR#4
10 mm
B
Leaf Area (mm2)
P < 0.0001
P < 0.0001
P < 0.0001
P = 0.9898
jaw-D;GR (MOCK)
jaw-D;GR (DEX)
AN3;GR#2 (MOCK)
AN3;GR#2 (DEX)
AN3;GR#3 (MOCK)
AN3;GR#3 (DEX)
AN3;GR#4 (MOCK)
AN3;GR#4 (DEX)
C
MOCK
DEX
jaw-D;GR
AN3;GR#1
AN3;GR#2
AN3;GR#3
AN3;GR#4
10 mm
D
Epidermal cell number
P < 0.0001
P < 0.0001
P < 0.0001
P = 0.9677
E
Subepidermal cell number
P < 0.0001
P < 0.0001
P < 0.0001
P = 0.3246

**Fig. S3. TCP4 signalling coordinates growth bidirectionally between epidermal and subepidermal layers.** (A) Representative images of the mature first leaf pair from 28-day-old plants of the indicated genotypes and chemical treatments. Scale bar, 10 mm. (B) Quantification of leaf area (Y-axis) for the mature first leaf pair of the indicated genotypes and treatments (X-axis). N = 15-28 leaves. Statistical significance was assessed by one-way ANOVA followed by a Holm-Šídák post hoc test (P-values indicated above the compared means). Error bars represent SD. (C) Representative images of whole rosettes and individual rosette leaves from the indicated genotypes under MOCK and dexamethasone (DEX) treatments, corresponding to inducible TCP4 expression from the endogenous pTCP4 promoter or the subepidermal (L2) pAN3 promoter. Scale bar, 10 mm. (D, E) Estimated epidermal (D) and subepidermal (E) cell numbers for the indicated genotypes and treatments (X-axis). Cell number was calculated as the ratio of total leaf area to mean cell area (8-15 leaves per genotype). Statistical analysis as in (B); error bars represent SD.

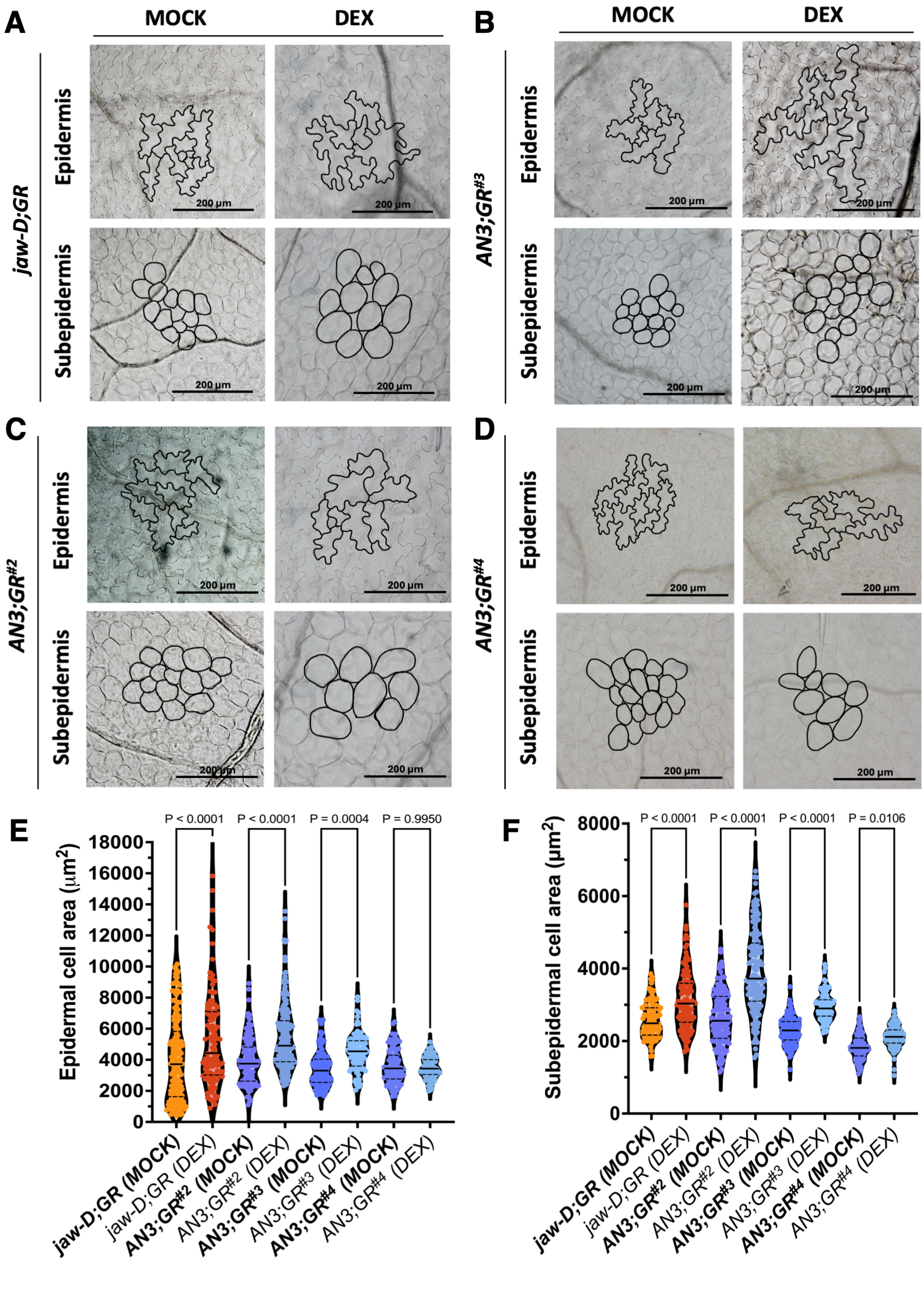
A
MOCK
DEX
jaw-D;GR
Epidermis
Subepidermis
200 µm
B
AN3;GR#3
C
AN3;GR#2
D
AN3;GR#4
E
Epidermal cell area (µm2)
P < 0.0001
P = 0.0004
P = 0.9950
F
Subepidermal cell area (µm2)
P = 0.0106
jaw-D;GR (MOCK)
jaw-D;GR (DEX)
AN3;GR#2 (MOCK)
AN3;GR#2 (DEX)
AN3;GR#3 (MOCK)
AN3;GR#3 (DEX)
AN3;GR#4 (MOCK)
AN3;GR#4 (DEX)

**Fig. S4. Subepidermal TCP4 expression non-autonomously regulates cell proliferation and expansion across tissue layers at near-endogenous levels.** (A-D) Differential interference contrast (DIC) images of epidermal pavement cells and subepidermal cells from mature first leaves of 28-day-old plants of the indicated genotypes and chemical treatments. Cell outlines, including pavement cells and stomata, are shown in black. Scale bars, 200 µm. (E, F) Violin plots showing the distribution of epidermal (E) and subepidermal (F) cell areas (Y-axis) for the indicated genotypes and treatments. N = 120-400 cells. Statistical significance was assessed by one-way ANOVA followed by a Holm-Šídák post hoc test (P-values indicated above the compared means).

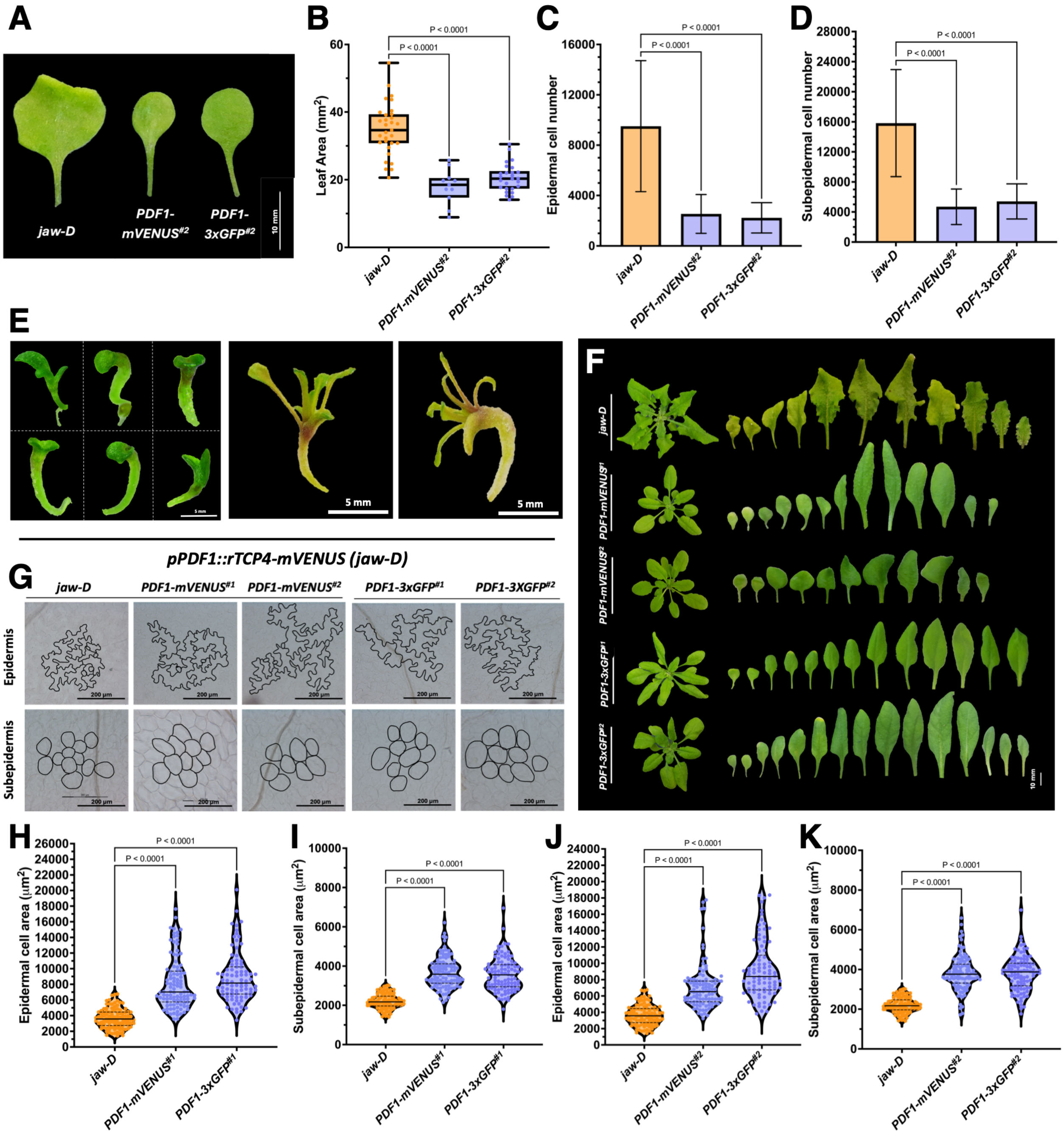


**Fig. S5. TCP4-mediated downstream signalling operates in a non-cell-autonomous manner.** (A) Representative images of the mature first leaf pair from 28-day-old plants of the indicated genotypes. Scale bar, 10 mm. (B) Quantification of leaf area (Y-axis) for the mature first leaf pair of the indicated genotypes (X-axis). N = 14-26 leaves. Statistical significance was assessed by one-way ANOVA followed by a Šídák post hoc test (P-values indicated above the compared means). Error bars represent SD. (C, D) Estimated epidermal (C) and subepidermal (D) cell numbers for the indicated genotypes and chemical treatments (X-axis). Cell number was calculated as the ratio of total leaf area to mean cell area (8-10 leaves per genotype). Statistical analysis as in

(B); error bars represent SD. (E) Representative T1 seedlings expressing *pPDF1::rTCP4-mVENUS* showing fused cotyledons, arrested leaf development, and seedling lethality upon constitutive epidermal TCP4 expression. Scale bar, 5 mm. (F) Representative whole rosettes and rosette leaves of homozygous weakest insertion lines *pPDF1::rTCP4-mVENUS* (#1 and #2) and *pPDF1::rTCP4-3xGFP* (#1 and #2), shown alongside the *jaw-D* control. (G) Differential interference contrast (DIC) images of epidermal pavement cells and subepidermal cells from mature first leaves of 28-day-old plants of the indicated genotypes. Cell outlines are shown in black. Scale bar, 200 µm. (H-K) Violin plots showing the distribution of epidermal (H, J) and subepidermal cell (I, K) areas for the indicated genotypes (Y-axis). N = 120-400 cells. Statistical significance was assessed by one-way ANOVA followed by a Šídák post hoc test (P-values indicated above the compared means).

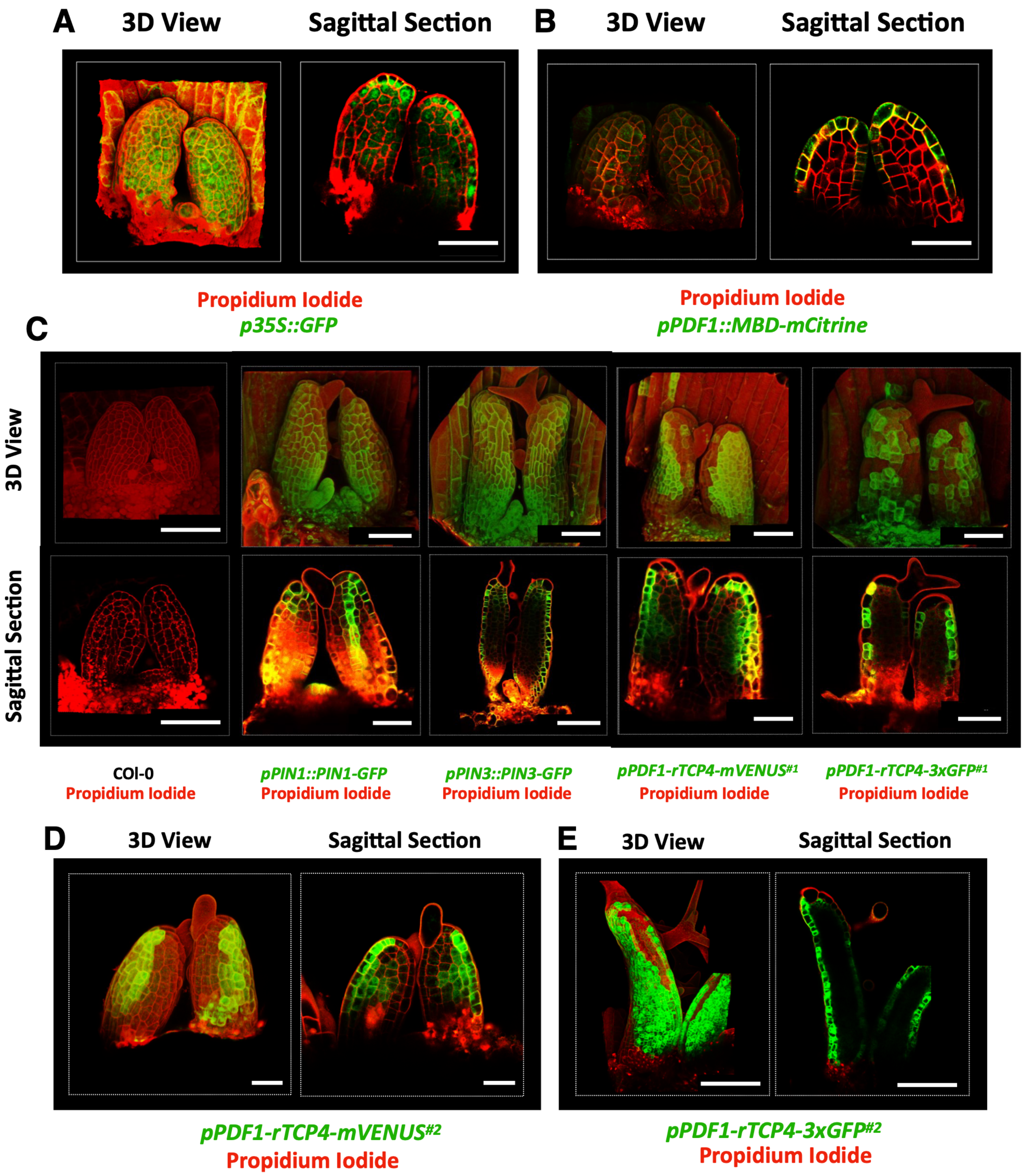


**Fig. S6. TCP4 protein is mobile across tissue layers, whereas a bulky fusion restricts its movement.** (A, B) Confocal images of propidium iodide-stained leaf primordia (2-3 DAI) from control lines. (A) *p35S::GFP* showing fluorescence in all tissue layers. (B) *pPDF1::MBD-mCitrine* showing fluorescence restricted to the epidermis. Three-dimensional reconstructions and optical mid-sagittal sections are shown. Scale bars, 50 µm. (C) Propidium iodide-stained confocal images of first leaf primordia (2-3 DAI) are shown as three-dimensional reconstructions and optical sagittal sections. Negative control (Col-0) shows no signal. Positive controls *pPIN1::PIN1-GFP* and *pPIN3::PIN3-GFP* display expected localisation patterns in L1/vasculature and all layers, respectively. In comparison, *pPDF1::rTCP4-mVENUS* (#1 and #2) shows fluorescence in the epidermis and adjacent subepidermal layers, whereas *pPDF1::rTCP4-3xGFP* (#1 and #2) fluorescence is confined to the epidermis. Scale bars, 50 µm. (D) Confocal images of propidium iodide-stained leaf primordia (2-3 DAI) from *pPDF1::rTCP4-mVENUS* insertion lines (#2). Three-dimensional views and optical sagittal show fluorescence in the epidermis and adjacent subepidermal layers, indicating limited inter-layer mobility of TCP4-mVENUS. Scale bars, 50 µm. (E) Propidium iodide-stained confocal images of first leaf primordia (5-6 DAI) from *pPDF1::rTCP4-3xGFP* (#2) are shown as three-dimensional views and optical mid-sagittal sections, confirming epidermis-restricted localisation at later developmental stages. Scale bars, 100 µm.

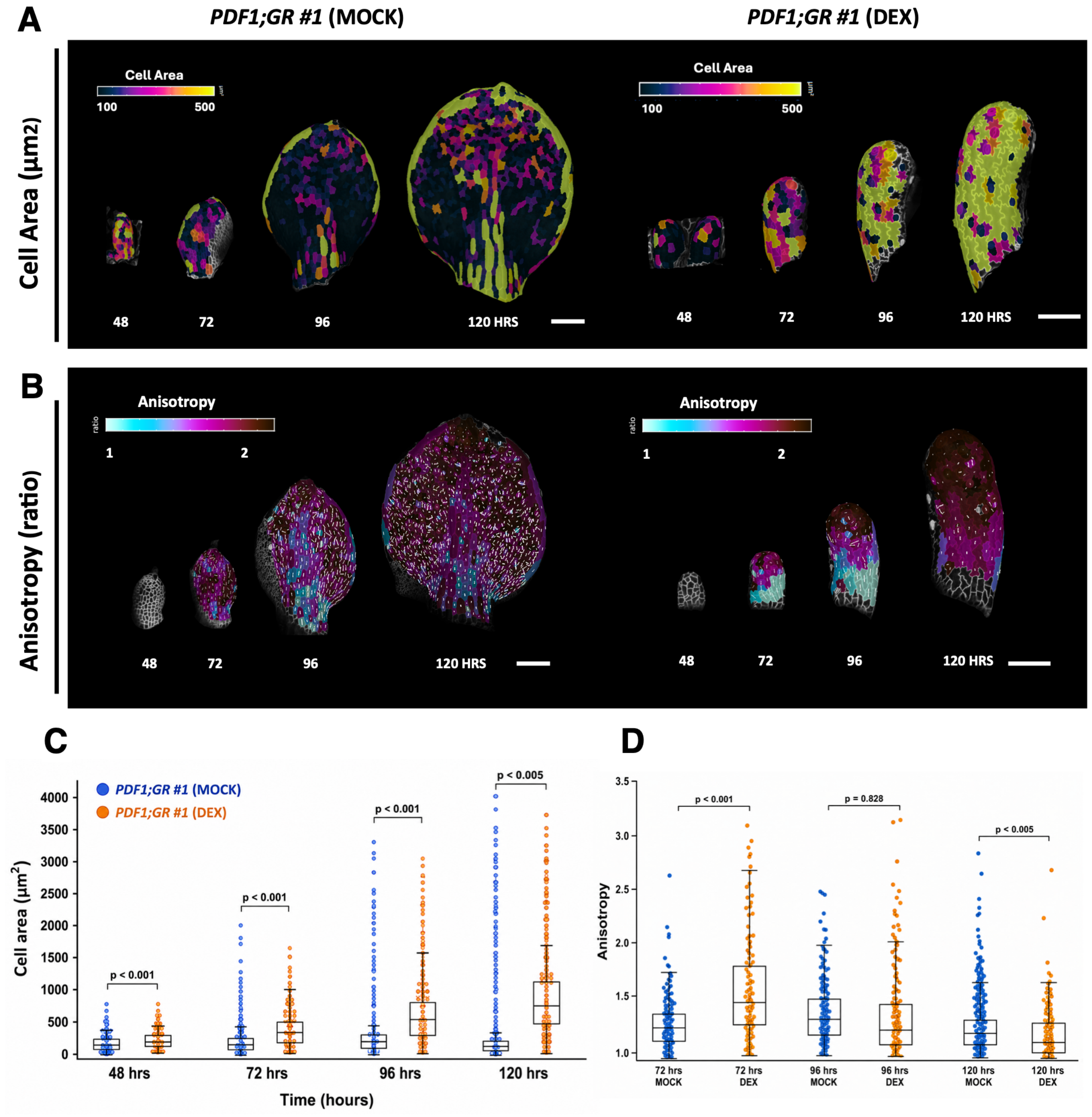


**Fig. S7. Epidermal TCP4 induction remodels cell area expansion and growth anisotropy during leaf morphogenesis.** (A) Heat maps depicting the area extension cellular area profile of the leaf primordia (*pPDF::rTCP4-GR #1* line in MOCK and DEX conditions) live-imaged from 48-120 hours after initiation. (B) Heat maps of growth anisotropy superimposed with vectors depicting principal directions of growth (max PDGs) calculated using live imaging (*pPDF1; GR*-MOCK and DEX) from 48-120 hours post leaf initiation. (C) Boxplots of area extension cell area (µm²) of three replicates of leaf primordium of *pPDF::rTCP4-GR #1* line in MOCK (blue) and DEX (orange) conditions. A Student's t-test was performed between the respective MOCK and DEX treatments across the time course, and p-values are indicated above the box plots. (D)

Boxplots showing growth anisotropy of three replicates of leaf primordium of *pPDF::rTCP4-GR* line in MOCK (blue) and DEX (orange) conditions. A Student's t-test was performed between the respective MOCK and DEX treatments across the time course, and p-values are indicated above the box plots.

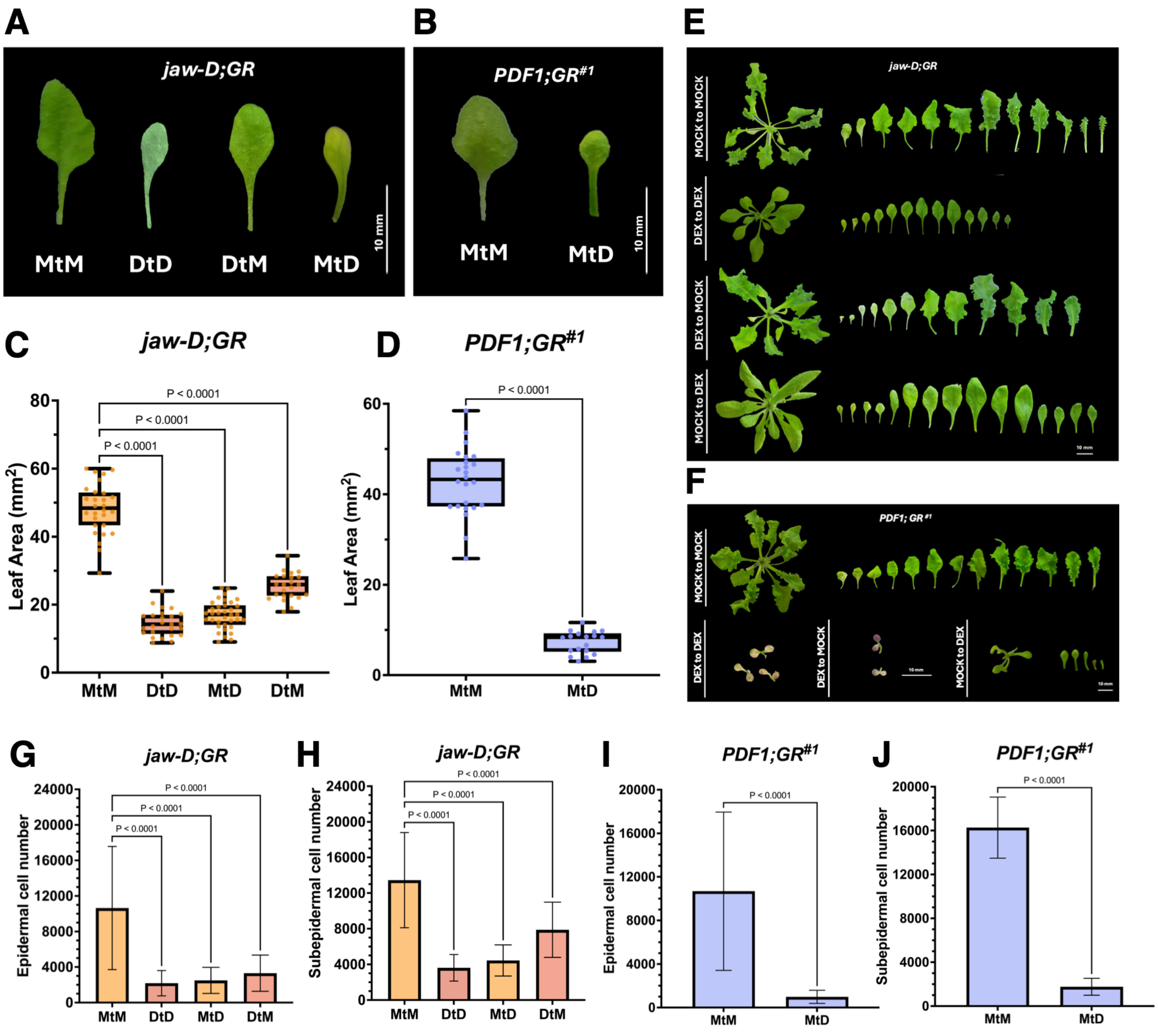


**Fig. S8. Ectopic epidermal TCP4 induction inhibits leaf formation in a temporally sensitive manner.** (A, B) Mature first leaves from 28-day-old plants of *jaw-D;GR* (*pTCP4::rTCP4-GR*) and *pPDF1;GR* (#1; *pPDF1::rTCP4-GR*) grown under staged dexamethasone (DEX; 12 µM) treatments. Seedlings were grown under MOCK conditions for the first 4 days and then transferred to MOCK (MtM) or DEX (MtD). Alternatively, seedlings were grown in DEX for the first 4 days and then transferred to DEX (DtD) or MOCK (DtM). Scale bar, 10 mm. (C, D) Quantification of leaf area (Y-axis) of the mature first leaf pair for *jaw-D;GR*, *pPDF1;GR* #1 under the indicated transfer treatments (X-axis). N = 20-34 leaves. Statistical significance was assessed by one-way ANOVA followed by a Šídák post hoc test (P-values indicated over the compared treatments). Error bars represent SD. (E) Representative whole rosettes and individual rosette leaves of *jaw-D;GR* plants under MOCK and continuous DEX treatment, corresponding to induction of TCP4 under its endogenous promoter (pTCP4) in the jaw-D background. Scale bar, 10 mm. (F) Representative whole rosettes and individual rosette leaves of *pPDF1;GR* #1 plants under MOCK and DEX and transfer treatments, corresponding to epidermis-specific induction of TCP4 in the *jaw-D* background. Scale bar, 10 mm. (G-I) Estimated epidermal and subepidermal cell numbers (Y-axis) for the indicated genotypes under the respective chemical transfer treatments (X-axis). Cell number was calculated as the ratio of total leaf area to mean cell area, based on 12-20 leaves per genotype. Statistical significance was assessed by one-way ANOVA followed by a Šídák post hoc test (P-values indicated over the compared treatments). Error bars represent SD.

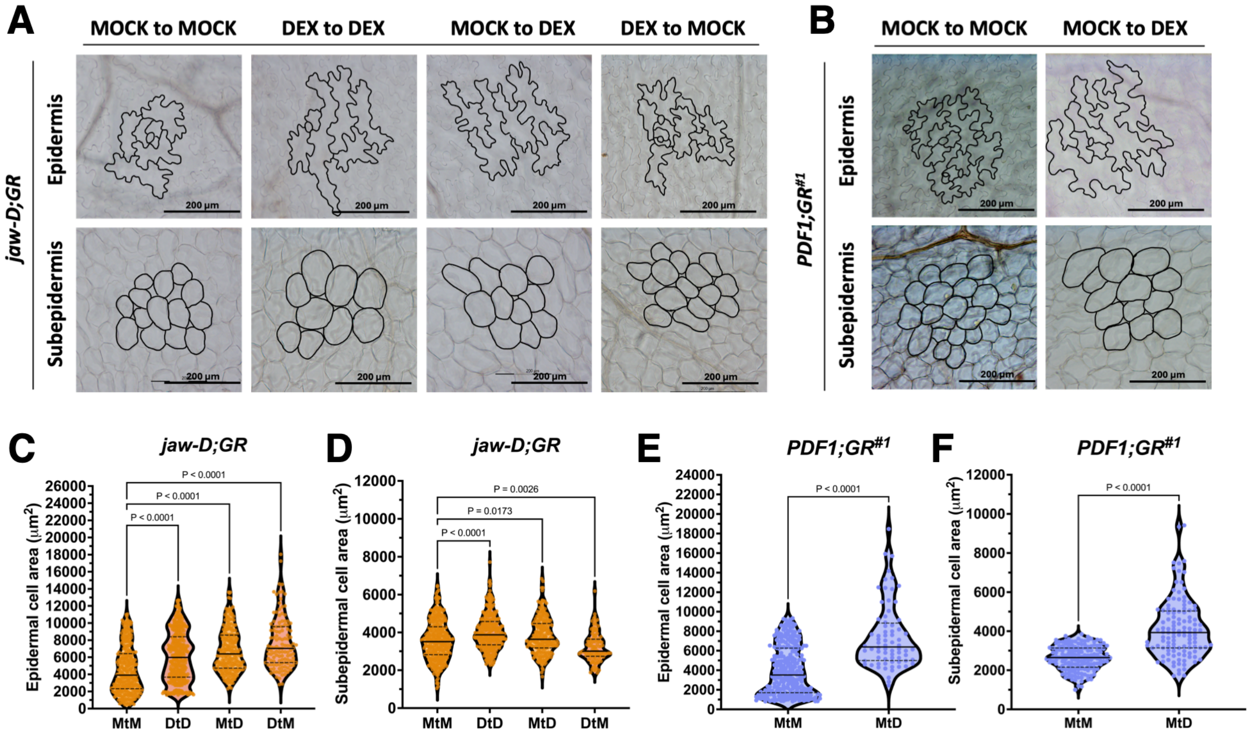


**Fig. S9. Epidermal TCP4 expression dominantly shapes cellular area through non-autonomous regulation across tissue layers**. (A, B) Differential interference contrast (DIC) images of epidermal pavement cells and subepidermal cells from mature first leaves of 28-day-old plants of the indicated genotypes and chemical transfer treatments. Cell outlines, including pavement cells and stomata, are shown in black. Scale bars, 200 µm. (C-F) Violin plots showing the distribution of epidermal and subepidermal cell areas (Y-axis) for the indicated genotypes and transfer treatments. N = 120-400 cells. Statistical significance was assessed by one-way ANOVA followed by a Šídák post hoc test (P-values indicated over the compared treatments). Error bars represent SD.

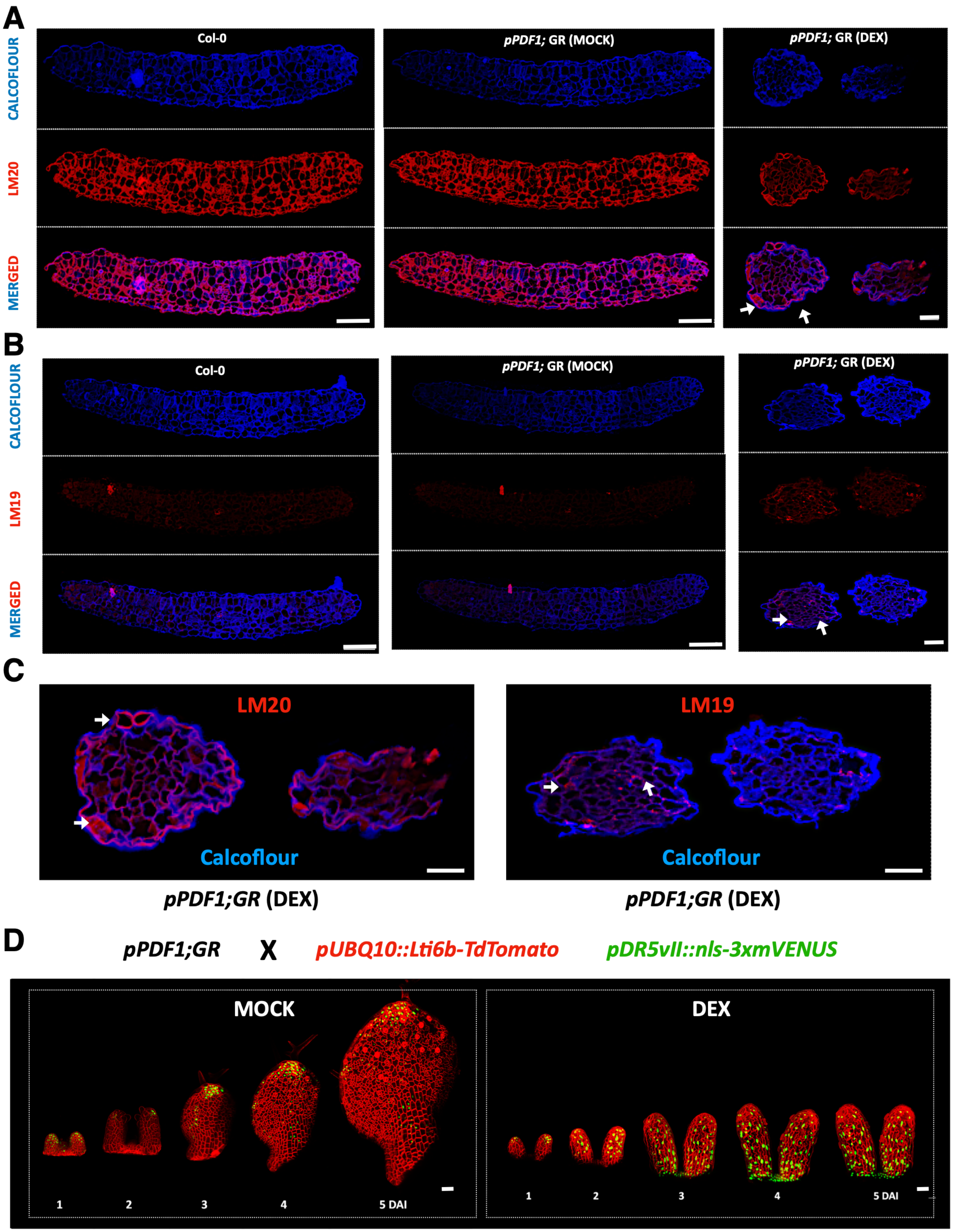
A
Col-0
pPDF1; GR (MOCK)
pPDF1; GR (DEX)
CALCOFLOUR
LM20
MERGED
B
Col-0
pPDF1; GR (MOCK)
pPDF1; GR (DEX)
CALCOFLOUR
LM19
MERGED
C
LM20
Calcoflour
LM19
Calcoflour
pPDF1;GR (DEX)
pPDF1;GR (DEX)
D
pPDF1;GR
X
pUBQ10::Lti6b-TdTomato
pDR5vII::nls-3xmVENUS
MOCK
DEX
1
2
3
4
5 DAI
1
2
3
4
5 DAI

**Fig. S10. Epidermal TCP4 induction triggers sustained auxin signalling and alters pectin-associated mechanical signatures.** (A, B) Confocal images of transverse sections of 6 DAI leaves (N=4) from Col-0, and *pPDF1::rTCP4-GR* leaves under MOCK and DEX conditions, immunolabeled with calcofluor (cell wall marker) and antibodies recognising pectin modifications. LM20 labels highly methylesterified pectin, whereas LM19 labels demethylesterified pectin. Merged images with calcofluor are shown. Arrows indicate ectopic deposition of LM20 in the epidermis of *pPDF1::rTCP4-GR DEX* samples (A) and increased LM19 signal in the subepidermal layer (B). Scale bars, 20 µm. (C) Zoomed inset of *pPDF1::rTCP4-GR* (DEX) depicting localisation of LM19 and LM20 antibodies/calcofluor. Arrows indicate ectopic deposition of LM20 in the epidermis of *pPDF1::rTCP4-GR DEX* samples and increased LM19 punctate signal in the subepidermal layer. Scale bars, 20 µm. (D) Time-lapse confocal imaging of the first leaf from *pPDF1::rTCP4-GR* crossed to *pUBQ10::Lti6b-TdTomato* (plasma membrane marker) and *pDR5vII::3xmVENUS* (auxin response reporter). Seedlings were imaged every 24 hours from 1 to 5 days after initiation (DAI) under MOCK and dexamethasone (DEX) treatments. DEX-treated primordia display sustained auxin response compared with MOCK controls. Scale bar, 50 µm.

**Supplementary Table S1:** Plant materials used in this study.

| Category | Line / Genotype | Genetic construct | Reporter / Tag | Background | Source |
|---|---|---|---|---|---|
| **Wild type control** | Col-0 | — | — | Col-0 | Standard laboratory accession |
| **Mutant control** | *jaw-D* | *MIR319A* overexpression (JAW-TCP knockdown) | — | Col-0 | Previously reported[40, 44] |
| **Promoter and protein reporters** | *pTCP4::GUS* | TCP4 promoter reporter | GUS | Col-0 | Previously reported[40, 44] |
| | *pTCP4::TCP4-GUS* | TCP4 translational fusion | GUS | Col-0 | Previously reported[40, 44] |
| | *pMIR319C::GUS* | MIR319C promoter reporter | GUS | Col-0 | Previously reported[40] |
| | *pPDF1::GUS* | Epidermis-specific promoter reporter | GUS | Col-0 | **Generated in this study** |
| | *pAN3::GUS* | Subepidermal promoter reporter | GUS | Col-0 | **Generated in this study** |
| **Inducible TCP4 lines** | *pTCP4::rTCP4-GR* | miR319-resistant *TCP4* fused to GR | GR | Col-0 | Previously reported[40, 44] |
| | *pTCP4::rTCP4-GR* | miR319-resistant *TCP4* fused to GR | GR | *jaw-D* | Previously reported[40, 44] |
| | *pPDF1::rTCP4-GR* | miR319-resistant *TCP4* | GR | *jaw-D* | **Generated in this study** |
| | *pAN3::rTCP4-GR* | miR319-resistant *TCP4* | GR | *jaw-D* | **Generated in this study** |
| **Constitutive TCP4 constructs** | *pPDF1::rTCP4-nls-mVENUS* | miR319-resistant *TCP4* | mVENUS | *jaw-D* | **Generated in this study** |
| | *pPDF1::rTCP4-nls-3xGFP* | miR319-resistant *TCP4* | 3×GFP | *jaw-D* | **Generated in this study** |
| **Reporter and marker lines** | *pDR5VII::nls-3xmVENUS* | Synthetic auxin-responsive promoter | mVENUS | Col-0 | Previously reported[51] |
| | *pUBQ10::myr-tdTomato* | Constitutive | tdTomato | Col-0 | Previously reported[51] |

| | | | | | |
|---|---|---|---|---|---|
| | *pUBQ10::myr-YFP* | UBQ10 promoter | myr-YFP | Col-0 | Previously reported[54] |
| | *pPDF1::MBD-mCitrine* | PDF1 promoter | MBD-mCitrine | Col-0 | Previously reported[52] |
| | *pUBQ10::Lti6b-tdTomato* | UBQ10 promoter | *Lti6b-tdTomato* | Col-0 | Previously reported[52] |
| | *p35S::GFP* | 35S promoter | GFP | Col-0 | Gifted by MechanoDevo team RDP, Lyon |
| | *pPIN1::PIN1-GFP* | Endogenous (pPIN1) | GFP | Col-0 | Previously reported[51] |
| | *pPIN3::PIN3-GFP* | Endogenous (pPIN3) | GFP | Col-0 | Previously reported[51] |
| **Genetic crosses for imaging** | *pPDF1::rTCP4-GR × pUBQ10::myr-YFP* | Plasma membrane marker cross | YFP | *jaw-D* | **Generated in this study** |
| | *pPDF1::rTCP4-GR × pUBQ10::myr-tdTomato* | Plasma membrane marker cross | tdTomato | *jaw-D* | **Generated in this study** |
| | *pPDF1::rTCP4-GR × pDR5VII::nls-3xmVENUS* | Auxin reporter cross | DR5-mVENUS | *jaw-D* | **Generated in this study** |
| | *pPDF1::rTCP4-GR × pPDF1::MBD-mCitrine* | Cortical microtubule marker cross | GFP | *jaw-D* | **Generated in this study** |

**Supplementary Table S2:** Primers used in this study.

| S.No. | Primer Detail | Primer Name | Primer Sequence (5'-3') |
|---|---|---|---|
| 1 | To amplify *pPDF1* from Col-0 genomic DNA | PO3655 | FP:CCGaagcttTGGATAGCGGAATAGCTGGCA |
| | | PO3656 | RP:ACGgtcgacGAGAATTCACTGAGATTCAGAG |
| 2 | For confirming positive clones in colony PCR for *pAN3* positive clones | P01502 | FP:CTCGGATCCATTTTTGGTACCAACG |
| | | P01503 | RP:TTTGGATCCGATCAGAGGTAACATTGCTGGGG |
| 3 | For genotyping rTCP4-GRlines | PO4122 | FP:ACGgttaacCTGACGACCAATTCCATCACC |
| | | PO4123 | RP:ACGgatatcCAGTCATTTTTGATGAAACAGAAGC |
| 4 | For genotyping NLS-3X-eGFP lines | PO4135 | FP:CGCACTAGTCGTAATACGACTCACTATAGGGCGAAT |
| | | PO4134 | RP:CCGGTTAACGATCGGGGAAATTCGGATCTTTACTTG |
| 5 | For genotyping m-VENUS lines | PO4115 | FP:CGCAGATCTATGGTGAGCAAGGGCGAGGAGCTGT |
| | | PO4116 | RP:CCGGTTAACTTACTTGTACAGCTCGTCCATGCC |
| 6 | For genotyping GR lines | PO4180 | FP:ctgGGATCCcgggatcctgaagctcgaaaaac |
| | | PO4179 | RP:TTCgatatcTTTTTGATGAAACAGAAGCTTTTTG |
| 7 | For cloning miR319c and genotyping positive lines | PO4295 | FP:ACGC*gtcgac*aagtgcatgtgtttgaaactattc |
| | | PO4296 | RP:CATG*ccatgg*tctccagcacgtggacatga |
| 8 | For checking TCP4 insert in transgenics | P0022 | FP:TGATTGGCTCTTCAACGGAGGC |
| | | P0021 | RP:CAAGGATCTAGCCTGAACAGAGCA |

**Supplementary Table S3:** Reagents used in this study.

| Reagent / Material | Supplier | Catalog number | Application |
|---|---|---|---|
| Murashige and Skoog (MS) basal medium | HiMedia Laboratories | PT011 | Arabidopsis seed germination medium |
| Murashige and Skoog basal salts | Duchefa Biochemie | M0222 | Preparation of ½ MS growth medium |
| Agar (plant tissue culture grade) | Duchefa Biochemie | P1001 | Solidification of plant growth medium |
| Sucrose | Sigma-Aldrich | S9378 | Carbon source in plant growth medium |
| Plant Preservative Mixture (PPM) | Plant Cell Technology | PPM™ | Antimicrobial additive for plant culture |
| Propidium iodide | Sigma-Aldrich | P4170 | Cell wall staining for confocal imaging |
| ProLong™ Gold Antifade Mountant | Thermo Fisher Scientific | P36930 | Mounting medium for fluorescence imaging |
| Dexamethasone (DEX) | Sigma-Aldrich | D4902 | Induction of GR-fusion constructs |
| Ethanol (molecular biology grade) | Sigma-Aldrich | E7023 | Seed sterilization and tissue dehydration |
| Acetone | Sigma-Aldrich | 179124 | Tissue permeabilization and clearing |
| Sodium dodecyl sulfate (SDS) | Sigma-Aldrich | L3771 | Seed sterilization |
| Tween-20 | Sigma-Aldrich | P1379 | Surfactant used in staining buffers |
| Hygromycin B | Thermo Fisher Scientific | 10687010 | Selection of transgenic plants |
| Silwet L-77 | Sigma-Aldrich | L7607 | Surfactant for floral dip transformation |
| InsTAclone PCR Cloning Kit | Thermo Scientific | K1213 | TA cloning of promoter fragments |
| pCAMBIA1390 binary vector | CAMBIA | — | Plant transformation vector |
| pCAMBIA1300 binary vector | CAMBIA | — | Binary cloning vector |
| HindIII restriction enzyme | Thermo Fisher Scientific | ER0501 | DNA cloning |
| SalI restriction enzyme | Thermo Fisher Scientific | ER0641 | DNA cloning |
| NcoI restriction enzyme | Thermo Fisher Scientific | ER0571 | DNA cloning |
| EcoRI restriction enzyme | Thermo Fisher Scientific | ER0271 | DNA cloning |

| | | | |
|---|---|---|---|
| HpaI restriction enzyme | Thermo Fisher Scientific | ER0521 | DNA cloning |
| ApaI restriction enzyme | Thermo Fisher Scientific | ER1411 | DNA cloning |
| BamHI restriction enzyme | Thermo Fisher Scientific | ER0051 | DNA cloning |
| SacI restriction enzyme | Thermo Fisher Scientific | ER1131 | DNA cloning |
| T4 DNA ligase | Thermo Fisher Scientific | EL0011 | DNA ligation |
| 5-bromo-4-chloro-3-indolyl β-D-glucuronide (X-Gluc) | Sigma-Aldrich | B4252 | Substrate for histochemical GUS staining |
| Potassium ferricyanide | Sigma-Aldrich | P8131 | Component of GUS staining buffer |
| Potassium ferrocyanide | Sigma-Aldrich | P9387 | Component of GUS staining buffer |
| Sodium phosphate buffer | Sigma-Aldrich | S0751 | Buffer for staining and fixation |
| Sodium metabisulfite | Sigma-Aldrich | S9000 | Component of mPS-PI staining |
| Hydrochloric acid (HCl) | Sigma-Aldrich | 258148 | Acid hydrolysis in pseudo-Schiff staining |
| Periodic acid | Sigma-Aldrich | 395132 | Oxidation step in pseudo-Schiff staining |
| Schiff reagent | Sigma-Aldrich | S5133 | Cell wall visualization in mPS-PI staining |
| Chloral hydrate | Sigma-Aldrich | C8383 | Leaf clearing reagent |
| Glycerol | Sigma-Aldrich | G5516 | Clearing and mounting medium |
| Glutaraldehyde | Sigma-Aldrich | G5882 | SEM fixation |
| Osmium tetroxide | Sigma-Aldrich | O5500 | SEM postfixation |
| Phosphate-buffered saline (PBS) | Sigma-Aldrich | P3813 | Sample washing |
| Histoclear | Sigma-Aldrich | H2779 | Tissue clearing for histology |
| Paraplast embedding medium | Sigma-Aldrich | P3683 | Tissue embedding for sectioning |
| Formaldehyde solution | Sigma-Aldrich | F8775 | FAA fixation |
| Glacial acetic acid | Sigma-Aldrich | A6283 | FAA fixation |
| LM19 antibody | Agrisera | AS12 1859 | Detection of de-esterified pectin |
| LM20 antibody | Agrisera | AS12 1860 | Detection of methyl-esterified pectin |
| Alexa Fluor™ 647 secondary antibody | Thermo Fisher Scientific | A-21235 | Immunohistochemistry detection |
| Calcofluor White | Sigma-Aldrich | F3543 | Cell wall counterstaining |

**Supplementary Table S4:** Morphometric measurements of leaf size, epidermal and subepidermal cellular parameters across genotypes and treatments (mean along with SD).

| Genotype | Line | Treatment | Leaf area (mm²) | Epidermal cell number | Subepidermal cell number | Epidermal cell area (µm²) | Subepidermal cell area (µm²) |
|---|---|---|---|---|---|---|---|
| *jaw-D* | — | Control | 35.42 ± 9.89 | 9661.90 ± 4297.94 | 16091.57 ± 6816.51 | 3665.28 ± 1247.92 | 2202.43 ± 381.82 |
| Col-0;GR | — | MOCK | 22.20 ± 3.40 | 4733.90 ± 1797.10 | 7797.80 ± 2748.70 | 4692.29 ± 1644.08 | 2849.89 ± 795.14 |
| | | DEX | 15.71 ± 3.71 | 2986.30 ± 1146.10 | 4069.60 ± 1381.00 | 5266.96 ± 1798.73 | 3864.03 ± 907.26 |
| *jaw-D;GR* | — | MOCK | 35.97 ± 7.00 | 9493.81 ± 5336.93 | 13525.40 ± 4213.40 | 3788.99 ± 1997.99 | 2659.03 ± 729.13 |
| | | DEX | 14.19 ± 2.72 | 2698.40 ± 1584.30 | 4131.30 ± 1145.30 | 5259.15 ± 2918.14 | 3433.66 ± 783.72 |
| *PDF1;GR* | #1 | MOCK | 34.73 ± 5.56 | 10969.20 ± 8666.00 | 14211.50 ± 3944.50 | 3167.36 ± 2279.48 | 3174.12 ± 664.12 |
| | #2 | MOCK | 43.15 ± 6.05 | 10519.20 ± 7481.10 | 12412.30 ± 3476.30 | 4100.49 ± 2273.99 | 3035.61 ± 719.91 |
| | #3 | MOCK | 32.23 ± 7.07 | 9009.51 ± 3117.27 | 15088.50 ± 4218.10 | 3578.13 ± 957.75 | 2135.94 ± 488.71 |
| | #3 | DEX | 15.15 ± 2.36 | 2498.40 ± 749.23 | 5525.20 ± 1537.60 | 5666.07 ± 1141.08 | 2562.13 ± 451.40 |
| | #4 | MOCK | 30.87 ± 8.85 | 8687.40 ± 2979.50 | 12412.30 ± 3476.30 | 3554.45 ± 1076.86 | 2487.63 ± 432.91 |
| | #4 | DEX | 25.59 ± 9.28 | 6138.60 ± 2558.20 | 10873.80 ± 3059.20 | 4168.63 ± 1008.28 | 2353.68 ± 509.26 |
| *AN3;GR* | #1 | MOCK | 40.76 ± 8.21 | 10333.94 ± 7223.14 | 10823.30 ± 4104.70 | 4041.66 ± 2710.85 | 3858.85 ± 1302.69 |
| | #1 | DEX | 17.15 ± 3.58 | 2665.80 ± 1284.72 | 4111.20 ± 1202.20 | 6433.69 ± 2793.38 | 4172.10 ± 920.04 |
| | #2 | MOCK | 37.15 ± 6.75 | 9433.60 ± 2021.60 | 14095.00 ± 4224.90 | 3939.22 ± 154.10 | 2636.49 ± 723.30 |
| | #2 | DEX | 14.02 ± 3.04 | 2587.60 ± 1069.60 | 3573.00 ± 1313.10 | 5419.65 ± 206.93 | 3923.30 ± 1157.03 |
| | #3 | MOCK | 29.39 ± 5.53 | 8593.40 ± 2088.10 | 12908.30 ± 3318.30 | 3422.13 ± 112.58 | 2227.36 ± 379.35 |
| | #3 | DEX | 15.33 ± 3.84 | 3403.10 ± 1200.20 | 5198.70 ± 1509.40 | 4508.20 ± 1191.23 | 2951.78 ± 391.18 |
| | #4 | MOCK | 24.06 ± 5.06 | 6747.70 ± 1563.90 | 12869.30 ± 4080.20 | 3588.61 ± 1086.22 | 1826.00 ± 330.80 |
| | #4 | DEX | 23.48 ± 4.63 | 6885.40 ± 2116.30 | 11349.30 ± 3140.20 | 3495.19 ± 684.42 | 2120.00 ± 308.29 |
| *PDF1-mVENUS* | #1 | — | 18.30 ± 4.39 | 2263.61 ± 1141.34 | 4978.62 ± 2207.26 | 8089.91 ± 3266.26 | 3678.22 ± 778.19 |
| | #2 | — | 17.93 ± 5.12 | 2536.78 ± 1539.35 | 4692.74 ± 2365.01 | 7067.72 ± 2792.11 | 3820.09 ± 880.32 |
| *PDF1-3xGFP* | #1 | — | 20.14 ± 2.17 | 2262.18 ± 926.42 | 5613.63 ± 2364.95 | 8902.15 ± 3199.33 | 3588.48 ± 844.22 |
| | #2 | — | 20.54 ± 4.16 | 2228.33 ± 1201.78 | 5401.89 ± 2342.61 | 9033.72 ± 2792.11 | 3803.36 ± 871.95 |

**Supplementary Table S5:** Percentage change in leaf morphometric parameters relative to respective controls.

| Genotype / Line | Treatment comparison | Leaf area (%) | Epidermal cell number (%) | Subepidermal cell number (%) | Epidermal cell area (%) | Subepidermal cell area (%) |
|---|---|---|---|---|---|---|
| ***Col-0;GR*** | **DEX vs MOCK** | −31.8 | −36.9 | −47.8 | +12.3 | +35.6 |
| ***jaw-D;GR*** | **DEX vs MOCK** | −56.0 | −71.6 | −69.4 | +38.8 | +29.1 |
| ***PDF1;GR#3*** | **DEX vs MOCK** | −53.1 | −72.2 | −63.4 | +58.4 | +20.0 |
| ***PDF1;GR#4*** | **DEX vs MOCK** | −17.1 | −29.3 | −12.4 | +17.3 | −5.4 |
| ***AN3;GR#1*** | **DEX vs MOCK** | −57.9 | −74.2 | −62.0 | +59.2 | +8.1 |
| ***AN3;GR#2*** | **DEX vs MOCK** | −62.3 | −72.6 | −74.7 | +37.6 | +48.8 |
| ***AN3;GR#3*** | **DEX vs MOCK** | −47.8 | −60.8 | −59.7 | +31.7 | +32.5 |
| ***AN3;GR#4*** | **DEX vs MOCK** | −2.4 | +2.0 | −11.8 | −2.0 | +16.1 |
| ***PDF1-mVENUS#1*** | **vs *jaw-D*** | −48.3 | −76.6 | −69.1 | +120.0 | +66.7 |
| ***PDF1-mVENUS#2*** | **vs *jaw-D*** | −48.6 | −73.3 | −70.4 | +92.1 | +72.8 |
| ***PDF1-3xGFP#1*** | **vs *jaw-D*** | −43.1 | −76.6 | −65.1 | +142.9 | +62.9 |
| ***PDF1-3xGFP#2*** | **vs *jaw-D*** | −48.6 | −73.3 | −70.4 | +145.5 | +72.0 |

**Supplementary Table S6:** Absolute morphometric measurements for transfer experiments.

| Genotype | Treatment | Leaf area (mm²) | Epidermal cell number | Subepidermal cell number | Epidermal cell area (µm²) | Subepidermal cell area (µm²) |
|---|---|---|---|---|---|---|
| ***jaw-D;GR*** | **MtM** | 48.20 ± 7.48 | 10640.00 ± 6931.60 | 13449.70 ± 5338.30 | 4527.31 ± 2729.36 | 3284.17 ± 1062.20 |
| | **DtD** | 15.41 ± 3.80 | 2179.40 ± 1424.00 | 3616.70 ± 1493.20 | 6365.47 ± 2607.54 | 4014.69 ± 928.11 |
| | **MtD** | 16.97 ± 4.08 | 2495.70 ± 1465.40 | 4442.70 ± 1749.00 | 6601.01 ± 2541.60 | 3819.63 ± 956.94 |
| | **DtM** | 25.36 ± 3.93 | 3303.50 ± 2037.40 | 7880.60 ± 3096.10 | 6478.64 ± 3021.33 | 3419.03 ± 774.77 |
| ***PDF1;GR#1*** | **MtM** | 42.77 ± 7.42 | 10686.70 ± 7267.40 | 16270.50 ± 2786.00 | 4002.79 ± 2474.13 | 2629.03 ± 629.01 |
| | **MtD** | 7.28 ± 2.49 | 979.10 ± 599.70 | 1763.70 ± 769.80 | 7437.80 ± 3335.58 | 4129.10 ± 1508.50 |

**Supplementary Table S7:** Percentage change in leaf morphometric parameters in transfer experiments relative to MtM control.

| Genotype | Treatment comparison (vs MtM) | Leaf area (%) | Epidermal cell number (%) | Subepidermal cell number (%) | Epidermal cell area (%) | Subepidermal cell area (%) |
|---|---|---|---|---|---|---|
| ***jaw-D;GR*** | **DtD** | -68.0 | -79.5 | -73.1 | +40.9 | +22.2 |
| | **MtD** | -64.8 | -76.5 | -67.0 | +45.8 | +16.3 |
| | **DtM** | -47.4 | -69.0 | -41.4 | +43.1 | +4.1 |
| ***PDF1;GR#1*** | **MtD** | -82.96 | -90.86 | -89.17 | +85.82 | +57.08 |
